# TrIdent - An R package to automate transductomics analysis of virus-like particle mediated DNA mobilization

**DOI:** 10.64898/2026.03.31.715651

**Authors:** Jessie Maier, Craig Gin, Jorden Rabasco, Avery Bass, Wynter Spencer, Breck A. Duerkop, Benjamin Callahan, Manuel Kleiner

## Abstract

**Background:** Transduction is a form of horizontal gene transfer in which bacterial DNA is packaged and transferred by virus-like particles (VLPs). Transductomics is a sequencing-based method used to detect DNA carried by VLPs. During transductomics analysis, reads from a sample’s ultra-purified VLPs are mapped to metagenomic contigs assembled from the same sample’s whole-community. The read mapping produces coverage patterns that require a time-consuming manual inspection and classification process which makes the method’s use unfeasible for datasets with many samples.

**Results:** We developed a novel algorithm, TrIdent (**Tr**ansduction **Ident**ification), that uses pattern-matching to automate the transductomics data analysis and that is available as an R package (https://jlmaier12.github.io/TrIdent/). There is no software equivalent to TrIdent so we compared TrIdent’s classifications of transductomics datasets to classifications made by human classifiers. TrIdent’s classifications were generally comparable to the manual classifications on a previously generated, manually classified transductomics dataset. When applied to newly generated transductomics data from the murine microbiota, TrIdent agreed with two independent human classifiers as much as the two independent human classifications agreed with each other. TrIdent classified transductomics datasets in a fraction of the time needed by human classifiers, and the classifications produced by TrIdent are fully reproducible. We used TrIdent to explore three murine gut transductomes and found that bacterial DNA associated with the Oscillospiraceae and Turicibacteraceae families was highly enriched in the DNA packaged by VLPs as compared to the whole community metagenomes.

**Conclusions:** The TrIdent software is a more accessible, more efficient, and more reproducible alternative to the manual inspection of read coverage patterns previously required for transductomics data analysis. To demonstrate the application of TrIdent, we analyzed transductomics datasets from murine fecal pellets and showed that specific low abundance bacterial families appear to be heavily involved in transduction.

## Introduction

Horizontal gene transfer (HGT), the exchange of genetic material between cells by means other than reproduction, is a major driver of prokaryotic evolution^1–4^. The three best described mechanisms of HGT are conjugation, transformation and transduction. Conjugation, the transfer of nucleic acid between cells via a pilus or dedicated secretion system, and transformation, the acquisition of free nucleic acid from the environment, only require two participants - the donor and recipient bacterial cells. Conversely, transduction involves three entities-the donor and recipient cells in addition to an extracellular particle that transports nucleic acid between the donor and recipient. The extracellular particle involved in transduction is typically a virus (bacteriophage in the context of prokaryotes), however other virus-like particles (VLPs) like gene transfer agents (GTAs)^5,6^ and particles produced by phage inducible chromosomal islands (PICIs)^7,8^ also serve as vectors for DNA transfer. Membrane vesicles (MVs) have also been implicated in DNA transfer between cells^9–12^ and while they are not VLPs, they are often co-purified with VLPs^13,14^ and therefore we will include MVs as vectors of transduction in the context of this manuscript.

The classical forms of transduction are generalized and specialized transduction. Generalized transduction occurs when a lytic phage “accidentally” initiates packaging on its bacterial host’s genome and generates phage particles carrying exclusively bacterial DNA^15^. Specialized transduction occurs when a prophage, a phage genome integrated into a bacterial genome, excises and removes a region of bacterial DNA directly adjacent to the prophage insertion site and packages it into a phage particle together with the phage genome^16^. Despite generalized and specialized being two of the most well-studied forms of transduction, many other forms exist such as lateral, GTA-mediated, and transduction mediated by selfish mobile genetic elements like PICIs and some plasmids^17,18^. While there are ample opportunities for gene exchange to occur via transduction, little is known about the impact that transduction has on genetic diversity and subsequent evolution in complex microbial communities.

Most methods available to detect HGT in microbiome samples investigate it from a “historical” perspective by looking for signatures of past HGT events in sequencing data^19,20^. These methods often infer HGT from the detection of nearly identical DNA sequences in unrelated genomes^21–23^, however, they cannot differentiate between the mechanism or timescale of transfer thus limiting our understanding of the individual contributions that transduction, transformation, and conjugation have to gene transfer in complex microbial communities. One method that can provide insights into potential actively occurring HGT is metagenomic high throughput chromatin conformation capture (MetaHi-C), which physically links the DNA within cells prior to sequencing and can be used to detect extrachromosomal mobile genetic elements (MGEs), however the method provides no context for how the MGEs are spreading^24–26^. In summary, there is a strong need for methods that measure active HGT in microbiomes and determine the respective mechanism of HGT.

Transductomics is a sequencing based technique that allows for the characterization of prokaryotic DNA carried by VLPs^27^. While carriage of prokaryotic DNA in VLPs does not conclusively show integration into a recipient genome, it represents the first half of the transduction process, specifically the mobilization of DNA from a prokaryote in a VLP. In transductomics, prokaryotic DNA carried by VLPs is identified by assessing read coverage patterns formed when sequencing reads from ultra-purified VLPs of a sample are mapped to contigs generated from the same sample’s whole-community (WC). Since transduction is generally a non-random process, different mechanisms of DNA packaging or encapsulation produce distinct read coverage patterns that can be classified^5,10,28–31^. Transductomics generates a ‘snapshot’ of a sample’s transductome-a term we use to define the genetic material actively carried by VLPs in a sample. Using the transductomics approach, Kleiner et al. (2020)^27^ showed evidence that a diversity of transduction events including generalized and specialized transduction, gene-transfer agent (GTA) mediated transduction, chromosomal island transduction, and several unknown transduction mechanisms were occurring in the murine intestinal microbiome^27^.

The transductomics approach described by Kleiner et al. (2020) relies on visual inspection and manual classification of hundreds to thousands of read coverage patterns and is extremely time consuming. The need for manual classification has prevented larger transductomics studies with high sample numbers that are necessary for translational and clinical research. Additionally, the inherent subjectivity introduced by manual classifications prohibits reproducible analysis. While most read coverage patterns produced by transductomics are distinct, some patterns possess characteristics of multiple types of transduction events thus allowing for different interpretations. Furthermore, coverage patterns produced by unknown packaging and/or transduction mechanisms could cause confusion and inconsistency amongst manual classifiers. Despite the impact that transductomics analysis could have on microbial ecology research, the described limitations are a barrier to any meaningful use of transductomics. We developed an automated transductomics analysis software that overcomes the constraints imposed by manual classification. Our software, TrIdent (**Tr**ansduction **Ident**ification), uses a novel algorithm to automatically inspect and classify read coverage patterns indicative of transduction events. TrIdent’s ability to quickly and reproducibly classify transductomics datasets improves the accessibility of transductomics and allows for the exploration of transductomes in diverse environments. Transductomics and TrIdent will improve our understanding of how transduction contributes to genetic diversity generation and subsequent evolution in microbial communities.

## Results and discussion

### The TrIdent software and its integration with the transductomics analysis workflow

We developed software to automate the transductomics pattern detection and classification process. The TrIdent R package^32^ automates the transductomics analysis by detecting, classifying, and computing read coverage pattern characteristics associated with potential transduction events (Fig. 1). TrIdent detects transduction events with a novel pattern-matching algorithm which uses predefined patterns that correspond to different mechanisms of DNA packaging. Predefined patterns include ‘Prophage-like’, ‘Sloping’ and ‘NoPattern’ patterns. The Prophage-like patterns, defined by rectangular shapes, represent read coverage patterns associated with integrated genetic elements like prophage, PICIs, and other MGEs. The ‘Sloping’ patterns, defined by triangular shapes, represent read coverage patterns associated with transduction events that transfer large regions of host DNA like generalized, lateral and GTA-mediated transduction. The ‘NoPattern’ pattern is defined by a horizontal line and represents contigs with no distinct read coverage pattern. Patterns are translated across each contig in the dataset and at each translation, a match-score is calculated which indicates how well the pattern matches the contig’s read coverage. After a pattern is fully translated across a contig, certain aspects of the pattern are changed (i.e. height, width, slope) and translation is repeated. This process of translation and pattern re-scaling cycles until a large number of pattern variations are tested, as determined by the formula in Fig. S1. The number of patterns tested depends on contig length, for example, for a single 50,000 bp contig, 59,452 pattern variations are tested in total. While the pattern-matching computation time increases quadratically with contig length, a 525 kbp contig can be classified in an average of 51 seconds and therefore overall computational time remains low (Fig. S2). After the pattern-matching process is complete, the pattern associated with the best match-score is used to classify contigs as either ‘Prophage-like’, ‘Sloping’, or ‘NoPattern’. After pattern-matching is complete, contigs classified as ‘NoPattern’ proceed to an additional classification step in which contigs with a median 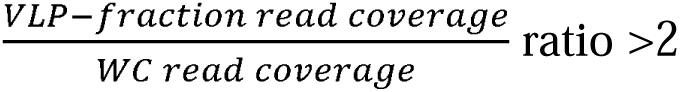 are reclassified as ‘HighCovNoPattern’. Contigs with large amounts of VLP-fraction read coverage relative to the WC may represent the ‘tails’ of generalized transduction events (see Fig. S2 Kleiner et al. (2020)), MV mediated transduction^10^ (also known as vesiduction^33^ or MV-mediated HGT^12^), GTAs^34^ or abundant viruses whose genomes have assembled into contigs. The pattern-matching evaluated in this manuscript is performed with TrIdent’s main function - TrIdentClassifier() and subsequent results are visualized with the plotTrIdentResults() function. The specializedTransductionID() function allows users to search for specialized transduction associated with Prophage-like classifications and while we did not evaluate the performance of this function, it is available for expert users. Detailed instructions on input data and usage of TrIdent is located at https://jlmaier12.github.io/TrIdent/.

**Figure 1.**
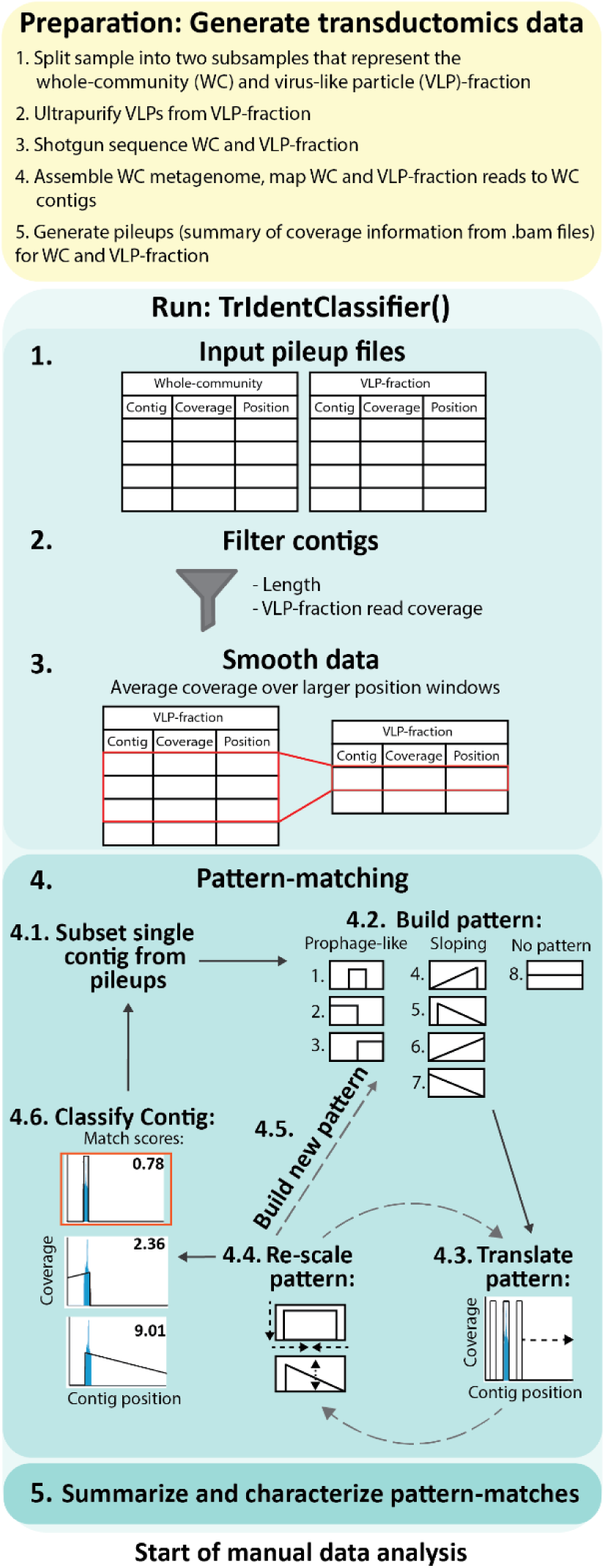
Overview of the TrIdent software. The yellow shaded box highlights the main steps for generating sequencing data compatible with the transductomics approach. Details in Kleiner et al. (2020). The teal shaded boxes highlight the various steps in the TrIdentClassifier() function in the TrIdent R package.

### Testing TrIdent against previously established transductomics data

We applied TrIdent to the dataset generated in the Kleiner et al. (2020) manuscript that introduced the transductomics method^27^ and compared TrIdent’s classifications to the manual classifications reported in that foundational manuscript. The Kleiner et al. dataset was generated from murine colon samples and the data was manually classified based on the contigs’ coverage patterns and gene annotations. We re-labeled the original classifications to be consistent with TrIdent’s pattern classification terms (re-labeling described in methods). The original classifications were highly descriptive and in many cases, mentioned specific MGEs or transduction mechanisms rather than general coverage patterns alone (S1 Table).

TrIdent classified the Kleiner et al. dataset, which consisted of 2,143 contigs, in ∼13 minutes. TrIdent’s classifications agreed with the manual ‘human’ classifications (excluding the ‘Unknowns’) 94.9% of the time. Transductomics data is highly imbalanced; the majority of contigs are not expected to show signs of transduction. Most of the agreements between TrIdent and human classifications reported in Kleiner et al. came from the negative ‘NoPattern’ class (Fig. 2). We further inspected the classification agreement between the human classifier and TrIdent in the positive set of contigs where some form of potential transduction was identified. TrIdent agreed with the human classifier’s ‘positive’ (Prophage-like, Sloping, and HighCovNoPattern) classifications 40.6% of the time. TrIdent was more conservative with its classifications, classifying 52% of the human classifier’s positive classifications (including the ‘Unknowns’) as NoPattern. We reviewed the coverage patterns of contigs with mismatched classifications (i.e. any contig where TrIdent and the human classifier did not agree) to better understand why the mismatches occurred (Fig. S3). Many of the human classifiers ‘Prophage-like’ classifications that were classified as NoPattern by TrIdent were smaller than TrIdent’s default minimum Prophage-like pattern match size (default is 10 kbp). Many of the human classifier’s HighCovNoPattern and Sloping classifications that were classified as NoPattern by TrIdent were associated with contigs with complete VLP-fraction coverage, however the coverage was low in respect to the WC and therefore did not meet TrIdent’s filter for HighCovNoPattern classifications 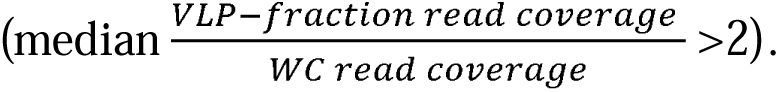 There were only 15 contigs in which the human classifier made NoPattern classification whereas TrIdent detected patterns. These contigs all had coverage patterns in the VLP-fraction (e.g. NODE_1269, NODE_649, and NODE_ 965 in Fig. S3), however, most were not associated with a known transduction mechanism, which may have been why they were classified as ‘NoPattern’ by the human classifier. Despite the classification differences, TrIdent still matched nearly half of the manual classifier’s positive classifications with the added benefits of speed and reproducibility. The time spent performing manual classification was not reported in Kleiner et al. 2020, but the manual inspection of 2,000+ contigs undoubtedly took longer than TrIdent’s 13 minute analysis. Furthermore, if the same human classifier performed the same classification task on the same contigs, there would almost certainly have been numerous differences in contig classification, whereas TrIdent produces the same classifications each time. These results demonstrated TrIdent’s ability to classify transductomics data in a manner comparable to humans with much higher reproducibility and lower effort.

**Figure 2.**
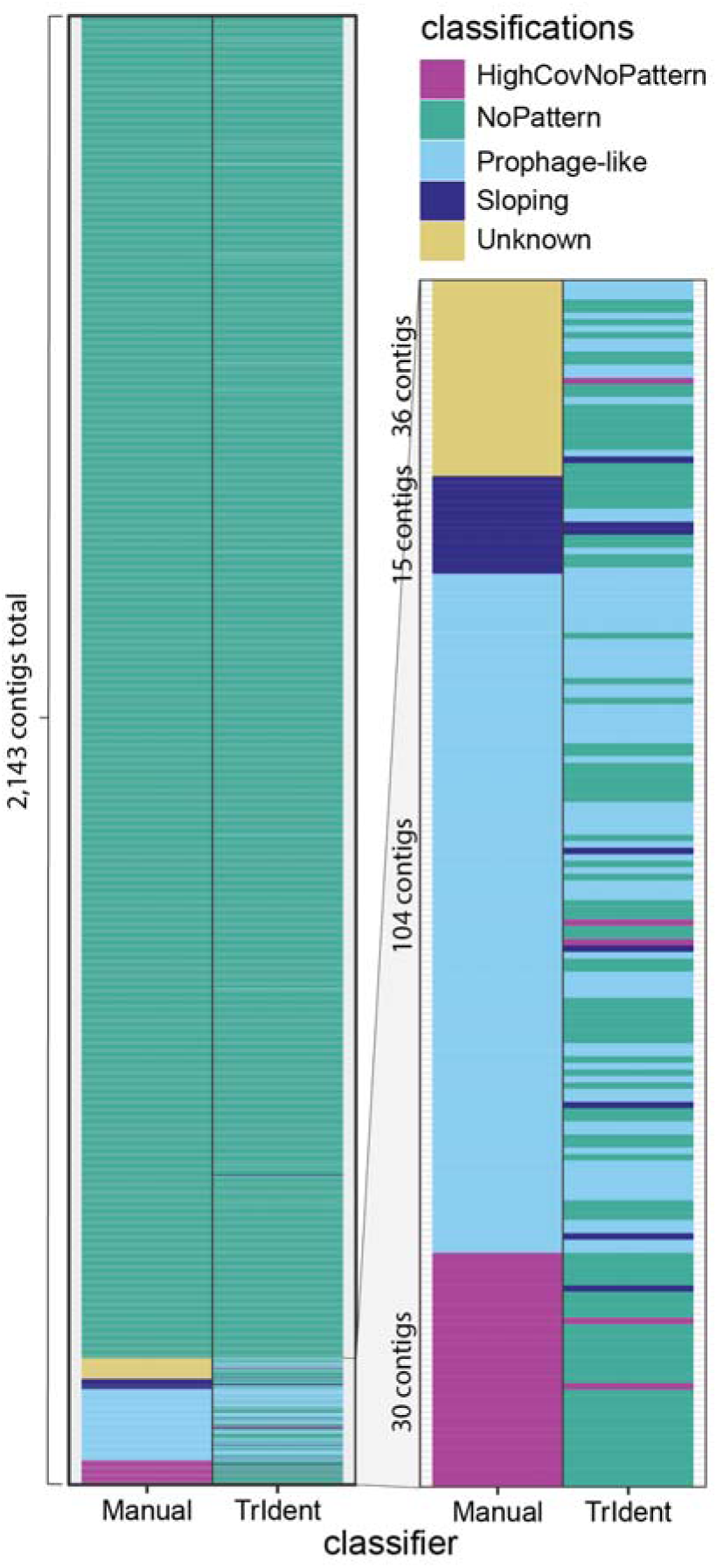
Comparison of manual and TrIdent classifications of a previously generated transductomics dataset^27^. The prior manual classifications were re-labeled for consistent use of classification terms in line with the ones used by TrIdent (Table S1). We ran TrIdent on the dataset with default parameters and compared the classifications (Fig. 2).

### Benchmarking TrIdent using human classifiers

To benchmark TrIdent’s performance, we compared it to the performance of two human classifiers on a new transductomics dataset. The human classifications represent the inherent variability introduced with manual analysis and set the baseline level of reproducibility and accuracy which currently exist for transductomics, for which there are no other computational tools. We generated new transductomics datasets from a mouse subjected to two different gut microbiome perturbations designed to induce variation in the observable transductomes. We collected fecal pellets from the mouse at a base-line condition (Pre-ABX), immediately after treatment with the antibiotic cefoperazone (ABX), which leads to a strong reduction in gut microbiota diversity^35^, and during *Clostridioides difficile* infection (CDI), which leads to severe toxin-mediated gut inflammation^36,37^. We shotgun sequenced the WC and VLP-fraction fecal metagenomes from all three samples. We assembled the WC samples individually, filtered for contigs greater than 30 kbp, and mapped each sample’s WC and VLP-fraction reads against their respective filtered assemblies to generate transductomics datasets for benchmarking (details on procedures can be found in https://jlmaier12.github.io/TrIdent/articles/TrIdent-vignette.html). Two human classifiers (CL1 and CL2) uninvolved in TrIdent’s development classified the three transductomics datasets manually after receiving extensive training (see Methods). The three datasets consisted of 1345, 26, and 682 contigs >30 kbp, respectively.

After the classifiers were finished, the datasets were analyzed with TrIdent using default parameters. Each of the human classifiers (CL1 and CL2) maintained consistent ratios of classifications across the two largest datasets (Pre-ABX and CDI) with CL1 classifying less contigs as NoPattern compared to CL2 (Fig. 3B). All three datasets were dominated by NoPattern classifications which highlights the large imbalance of ‘positive’ (Prophage-like, Sloping, and HighCovNoPattern) to ‘negative’ (NoPattern) classifications in transductomics datasets. To determine how TrIdent’s pattern-matching functionality compared to the human classifiers, we calculated the percentage of times that the classifiers (CL1, CL2, and TrIdent) made classifications that agreed with each other on each of the three datasets (Fig. 3B). We found that there were no statistically significant differences (Wilcoxon, p < 0.05) between comparison groups (CL1 + CL2 agree, CL1 + TrIdent agree, CL2 + TrIdent agree, and all agree) indicating that TrIdent agreed with the humans no more or less than they agreed with each other. We choose to use percent agreement rather than more ‘traditional’ performance metrics such as accuracy, precision, and recall as these metrics all require a ‘ground truth’ which neither CL1, CL2, or TrIdent represent. We also calculated Cohen’s Kappa, a metric that measures inter-rater reliability, between each classifier (CL1 + CL2 agree, CL1 + TrIdent agree, CL2 + TrIdent agree) for each of the three datasets and found that, like the percent agreements, the values were not significantly different (Wilcoxon, p < 0.05) across comparison groups (Fig. S4). Finally, we compared TrIdent’s analysis time to the human classifiers and found that TrIdent classified the Pre-ABX, ABX and CDI datasets order-of-magnitude faster than the human counterparts. TrIdent’s classifications for the three datasets took 9.36 minutes, 43 seconds, and 2.1 minutes, respectively, whereas it took CL1 and CL2 hours to complete their classifications manually (Fig. 3C).

**Figure 3.**
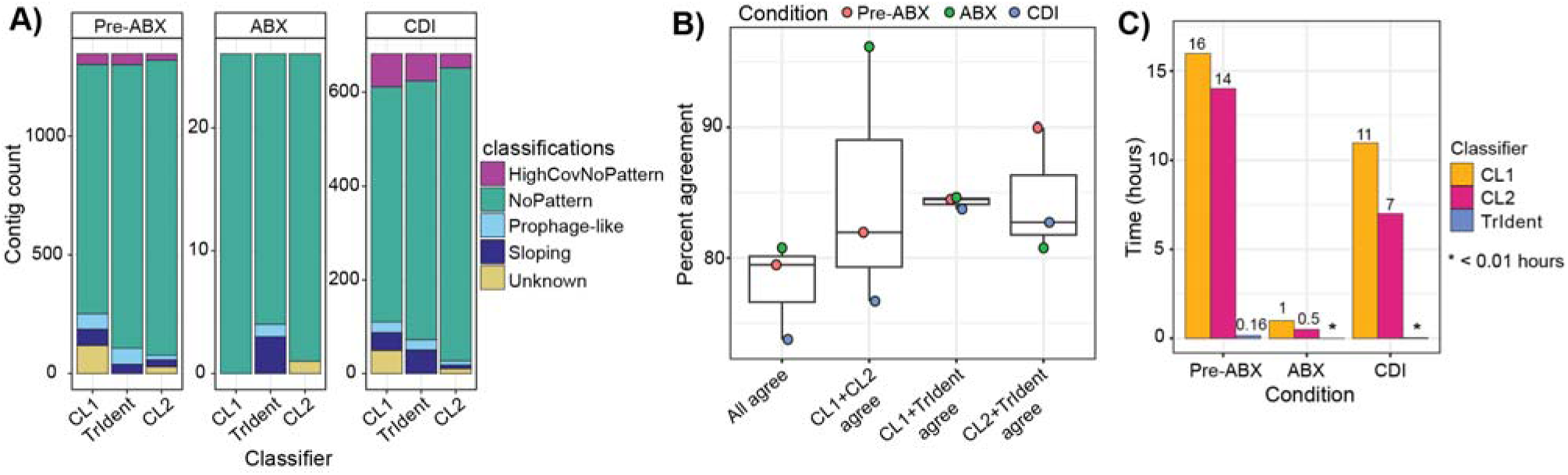
Evaluation of TrIdent’s pattern-matching performance against two human classifiers (CL1 and CL2). A) The composition of classifications made by each classifier (CL1, CL2 and TrIdent) for the three datasets analyzed. B) The percent agreement between classifications made by different classifiers across pre-antibiotics (Pre-ABX), antibiotics (ABX), and *C. difficile* infection (CDI) datasets. Percent agreement was calculated based on contigs that received the same specific classification (Prophage-like, Sloping, HighCovNoPattern, or NoPattern) between classifiers. C) A comparison of the time spent (in hours) classifying the Pre-ABX, ABX, and CDI datasets by each of the three classifiers (CL1, CL2 and TrIdent).

Since the datasets are imbalanced (majority NoPattern classifications, Fig. 3A), we also compared the three classifiers’ classifications at the individual class level (Prophage-like, Sloping, HighCovNoPattern, and NoPattern) to gain a better understanding of where and why differences in classifications occurred (Fig. 4). Unsurprisingly, CL1, CL2 and TrIdent overwhelmingly agreed with each other for the ‘NoPattern’ classifications which are relatively easy to detect both visually and computationally based on the near absence of VLP-fraction read coverage. As expected, there was more disagreement between both the human classifiers and TrIdent and between the human classifiers themselves for the Prophage-like, Sloping and HighCovNoPattern classes. As previously mentioned, the read coverage pattern classification can be a subjective process and interestingly, TrIdent and CL1 seem to agree with each other more than CL2 (Fig. 4A). Despite receiving the same training, CL1 and CL2 interpreted the pattern classes differently leading to differences in their classifications. This exemplifies the lack of reproducibility due to subjectivity during manual classification of transductomics data and highlights the need for an automated solution like TrIdent.

**Figure 4.**
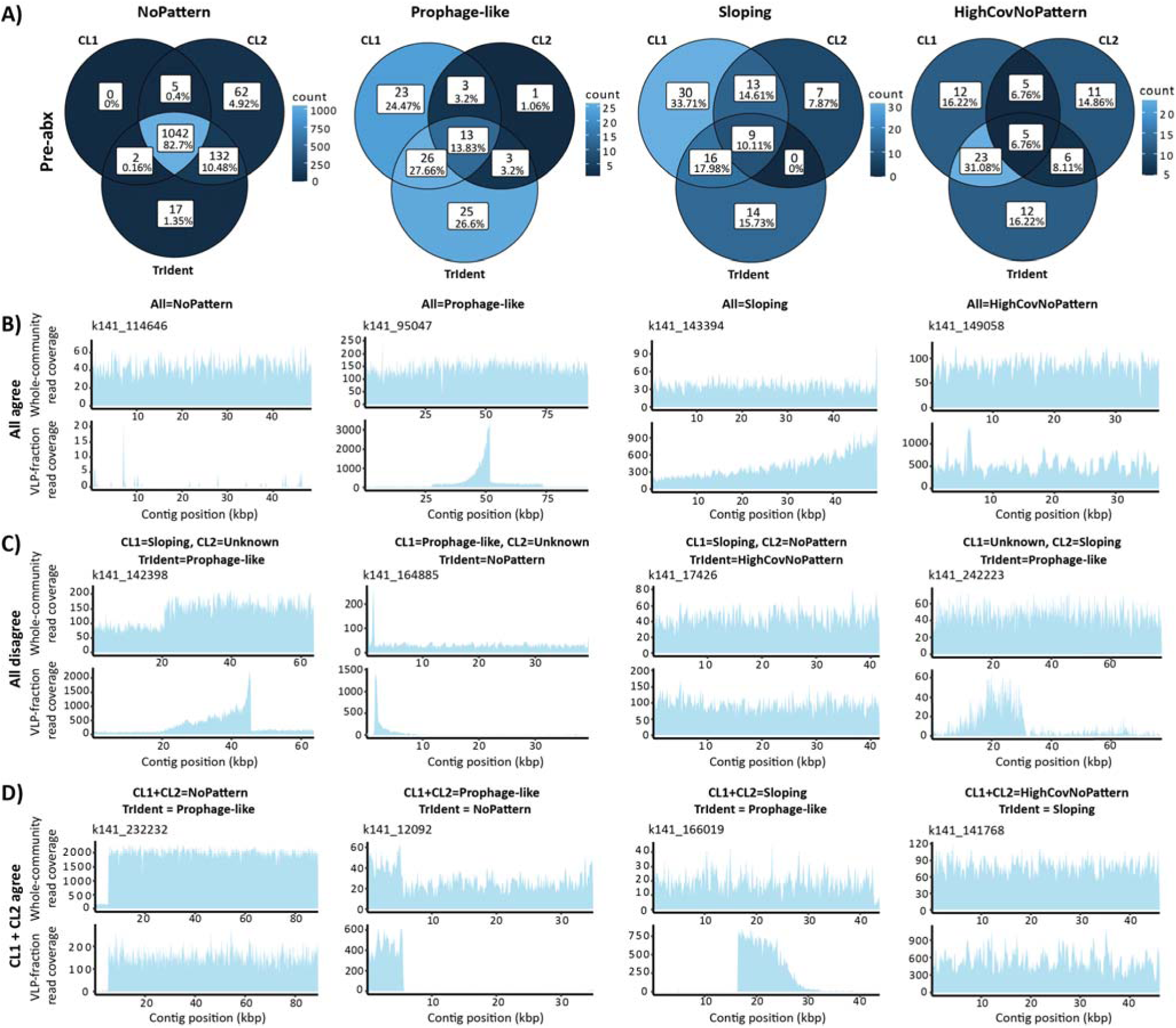
Comparison of classifications made by TrIdent and manual classifiers. A) Comparison of classifications made by the three classifiers (CL1, CL2, and TrIdent) in the pre-ABX dataset for each of the four TrIdent pattern-classes - NoPattern, Prophage-like, Sloping, and HighCovNoPattern. B) Examples of contigs in each of the four pattern classes in which all three classifiers agreed on the classification. C) Examples of contigs in which all three classifiers disagreed with each other on the classification. D) Example of contigs in which CL1 and CL2 agreed but TrIdent did not.

While only the comparisons from the pre-ABX dataset are visualized in Figure 4A, the Venn diagram patterns between classifiers are largely consistent across the post-ABX and CDI datasets (Fig. S5 and Fig. S6). The contigs in which all three classifiers agreed tend to have distinct read coverage patterns that are highly characteristic of the respective pattern classes (Fig. 4B). The contigs in which there were disagreements between classifiers often had read coverage patterns with characteristics of multiple different pattern classes which likely led to confusion amongst the human classifiers (Fig. 4C and 4D). For example, there were several contigs in which the human classifiers disagreed with TrIdent’s Prophage-like classification and each of these contigs had a defined block of coverage (characteristic of the Prophage-like pattern class) that sloped inwards or outwards (characteristic of the Sloping pattern class). These coverage patterns are typical of *cos* prophage whose use of 5’ and 3’ cohesive ends during packaging produce deceptive sloping patterns^38^. The sloping produced by *cos* phage can be differentiated from the sloping patterns produced by generalized, lateral and GTA-mediated transduction based on the length of the read coverage pattern. Generalized, lateral and GTA-mediated transduction typically spans tens to hundreds of kbps whereas the sloping associated with *cos* phage will generally be shorter than or similar length to the associated phage genome^27,28,31,38^. TrIdent considers these size differences during pattern-matching and classification, whereas the human classifications appear to be influenced more by the visual aspect of the pattern alone. Classification disagreements also often had to do with modifiable parameters in TrIdent’s pattern-matching functionality. For example, in Figure 4D, TrIdent classified the ∼5 kbp Prophage-like pattern on contig k141_232232 as NoPattern because TrIdent’s default minimum Prophage-like pattern match size is 10 kbp. When this parameter (minBlockSize) is changed to 5 kbp, TrIdent classifies this contig as Prophage-like.

### Case study to demonstrate TrIdent’s application

We generated transductomics datasets (see Methods) from the fecal pellets of three Pre-ABX mice to demonstrate TrIdent’s usage and exemplify the information that results from a transductomics study. We ran TrIdent with default parameters on each of the datasets and merged the output summaries to compare classifications of contigs across replicates. We ran geNomad^39^ on each WC assembly to detect plasmid, viral, and prophage-containing contigs. The geNomad software detects viruses, plasmids and prophage within metagenomic contig sets based on genomic features and gene annotations characteristic of each of these genetic elements. An average of 57% of the contigs classified by TrIdent as Prophage-like were classified by geNomad as viral or prophage-containing (Fig. 5A) which highlights the ability of transductomics to detect lytically active prophage present in the VLP-fraction, regardless if they are associated with transduction or not. Additionally, since TrIdent relies only on coverage data for its classifications rather than gene annotations, it can detect novel prophage or prophage-like elements, like PICIs for example, not present in sequence databases. An average of 27% of the contigs classified by TrIdent as Sloping were classified by geNomad as viral or prophage containing. These contigs likely contained coverage patterns associated with large *cos* phage which produced a sloping pattern longer than TrIdent’s default minimum Sloping pattern size (default is 20 kbp) ^38^.

**Figure 5.**
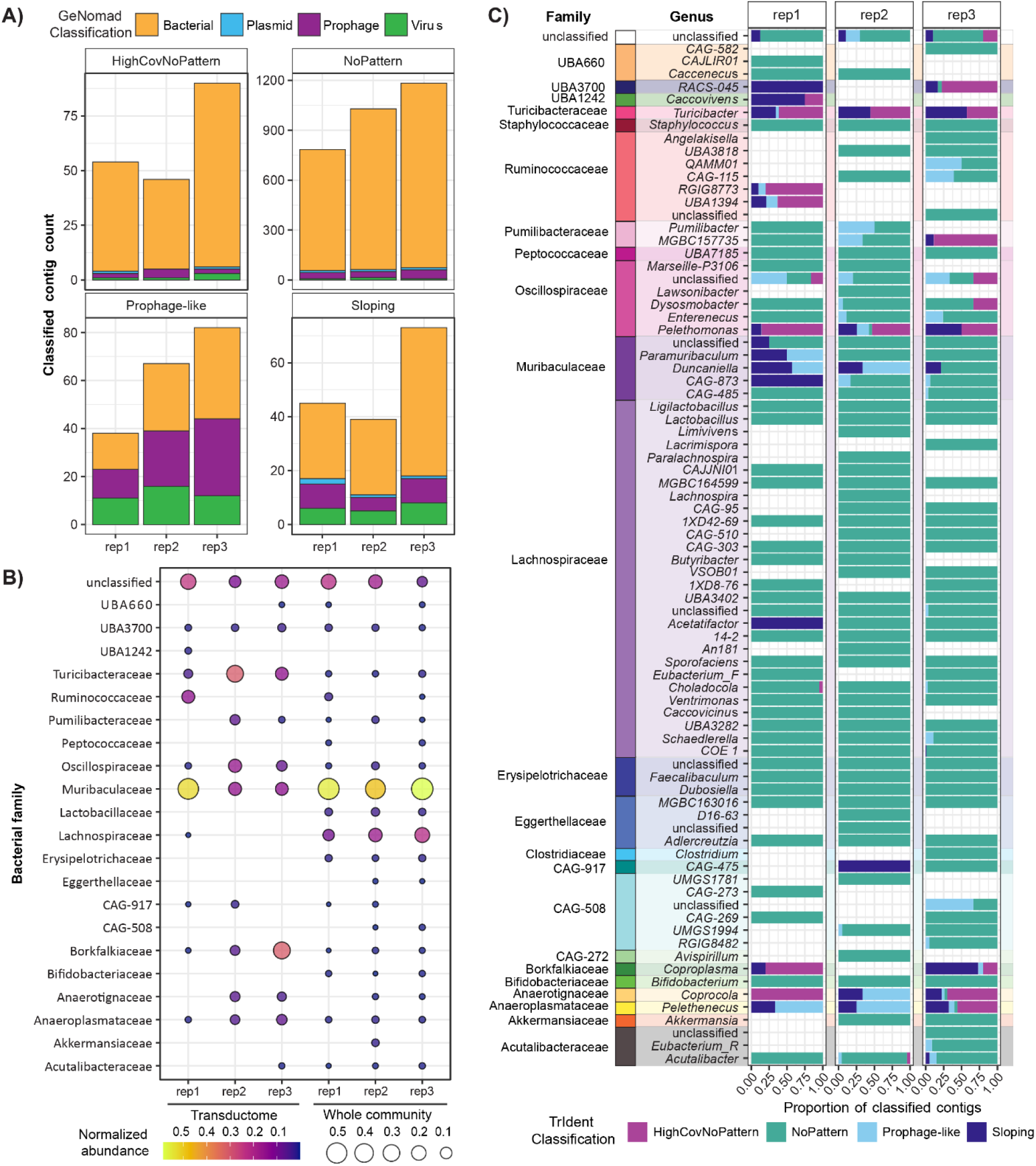
Composition of transductomes in murine feces case study. NoPattern classification counts contain both contigs that were filtered out prior to pattern-matching due to low read coverage and contigs which proceeded to pattern-matching but did not have a detectable read coverage pattern. A) Counts of contigs classified by TrIdent and their corresponding geNomad classifications. B) Comparison of bacterial family abundances in the WC vs the transductome. Contigs for which geNomad provided a virus or prophage prediction were removed prior to the relative abundance calculation. Abundance values were generated by mapping WC and VLP-fraction reads to WC MAGs and dividing by the total number of mapping reads for the respective WC or VLP sample. MAGs with less than 0.05% relative abundance in a replicate were removed in the associated replicate. The color and dot size scales represent the relative abundance values normalized to one in each individual replicate. C) Composition of TrIdent classifications (only for non-viral or prophage containing contigs) for all of the bacterial families and associated genera represented in the transductomes.

Prior to further analysis of the transductomes, we removed all contigs that geNomad predicted were viral or prophage containing to focus on pattern matches associated exclusively with packaging of bacterial DNA into VLPs. We determined the taxonomic compositions of the transductomes and compared them to the taxonomic compositions of the WC samples (Fig. 5B and 5C). We quantified taxa abundances by mapping WC and VLP-fraction reads from each sample to metagenome-assembled genomes (MAGs) associated with the respective WC. The WCs were composed of 54% Bacillota and 30% Bacteroidota on average which was in line with expected ratios for these phyla in male C57BL/6J mouse gut microbiota^40,41^. The composition of bacterial families in the transductomes were very different from taxonomic composition of the WC samples. Specific bacterial families were enriched in the transductomes relative to their abundance in the WC samples, whereas other families were not represented in the transductome at all (Fig. 5B). The Turicibacteraceae and Oscillospiraceae were some of the largest contributors to the gut transductomes making up averages of 16% and 6%, respetively, of the transductomes compared to their WC sample averages of 1%. Further, we found that only a few specific genera were responsible for the majority of transduction events associated with a family (Fig. 5C). There was only one genus, *Turicibacter*, associated with the Turicibacteraceae family and all of its contigs were classified by TrIdent as either Sloping, Prophage-like or HighCovNoPattern. There were at least six genera (including one unclassified) associated with the Oscillospiraceae family, however, the *Pelethomonas* and the unclassified genera were associated with the majority of the positively classified contigs. The majority of TrIdent classifications for both the *Turicibacter* and *Pelethomonas* genera are Sloping and HighCovNoPattern.

The Oscillospiraceae have been previously identified as a prevalent source of phage hosts in several gut microbiome studies^42,43^ and correspondingly, the Oscillospiraceae family is associated with the second largest number of prophage-containing contigs in our datasets (Fig. S7). It is conceivable that one or more of these phage is involved in transduction, likely using a method like generalized or lateral transduction which produces the coverage patterns typically associated with Sloping and HighCovNoPattern classifications. These types of coverage patterns are also produced by GTAs and due to their prophage origin, GTA clusters are often misclassified as phage. Interestingly, a recent study found large amounts of Oscillospiraceae DNA in the VLP-fractions of fecal homogenates which the authors hypothesized was the result of GTA producers in the Oscillospiraceae family^44^. There is little literature regarding the relationship between Turicibacteraceae and VLPs, however Turicibacteraceae represent an important component of the mammalian gut microbiome due to their role in bile acid and host lipid metabolism which aid in prevention of host obesity^45,46^. The Oscillospiraceae have been associated with beneficial short-chain fatty acid production and, like the Turicibacteraceae, their presence in the gut microbiome is correlated with host leanness^46,47^. The enrichment of Turicibacteraceae and Oscillospiraceae DNA in the transductomes suggests their involvement in transduction that may be influencing the respective roles these community members play in the murine gut microbiome. While we focused on the Oscillospiraceae and Turicibacteraceae due to the abundance of associated transduction events, contigs associated with other bacterial families like the Anaerotignaceae and Anaeroplasmataceae, for example, also had majority positive classifications by TrIdent and therefore warrant further investigation (Fig. 5C).

## Conclusions

Transduction is a major route of HGT yet research on the transductome is relatively sparse, largely due to a lack of accessible methods. Transductomics is a sequencing-based method used to detect bacterial genomic DNA actively carried by VLPs and make inferences about the packaging mechanism used^27^. We automated the transductomics analysis with a novel pattern-matching algorithm that detects read coverage patterns associated with potential transduction events. We integrated the algorithm into an R package, TrIdent^32^, which makes transductomics data analysis fast, efficient, and user-friendly. We compared TrIdent’s performance to that of human classifiers and found that TrIdent’s classifications were comparable to the humans’ classifications. Most of TrIdent’s misclassifications could be explained by parameters in the pattern-matching algorithm, many of which are user-defined and can therefore be altered to the user’s specific needs (i.e. minimum and maximum pattern sizes). That being said, TrIdent also made misclassifications that were caused by limitations in the pattern-matching algorithm itself. TrIdent can currently only detect one pattern per contig which means that long contigs with multiple patterns can lead to incorrect classifications. Additionally, coverage patterns that have characteristics of multiple pattern classes (i.e Prophage-like patterns with sloping regions at the sides or middle) can lead to misclassifications or pattern matches that do not fully match the contig’s coverage pattern (e.g. contigs k141_17365, k141_49595, k141_95047, Fig. S8). In a future version of TrIdent, these limitations could be addressed by implementing chunking of contigs to allow for the detection of multiple patterns per contig, adding additional pattern types, and further refining the pattern-matching algorithm with additional transductomics datasets. However, despite these limitations, TrIdent still outperforms human classifiers in terms of overall efficiency and reproducibility. To demonstrate TrIdent’s usage, we used it to analyze transductomics datasets generated from murine fecal pellets and found enrichments of bacterial DNA associated with two low abundance families - the Oscillospiraceae and Turicibactericeae, both of which have strong associations with host health. These preliminary results indicate that members of these bacterial families are heavily involved in transduction but further investigation and additional transductomics experiments are needed to better understand if the associated transduction affects the functions of these bacteria in the mammalian gut microbiome. The transductome is currently an unexplored component of every microbiome on Earth. Transductomics and TrIdent provide researchers with a toolset to investigate the transductome and better understand the role of transduction in microbial ecology.

## Methods

### TrIdent software development

#### Preparing data for pattern-matching

TrIdent detects read coverage patterns using a pattern-matching algorithm that operates on pileup files. A pileup file is a file format where each row contains the average number of mapped sequencing reads at a specific genomic location. Using a rolling mean of read coverages both smooths read coverage patterns and reduces data size relative to the associated.bam file. We use BBMap’s pileup.sh script to generate pileup files with a bin, or window, size of 100 bp as input for TrIdent. We found that averaging read coverage over 100 bp windows provides a balance between file size and read coverage pattern resolution, both of which increase when window size decreases. After generating the pileup files, we filter out contigs that are too short or have little to no read coverage. TrIdent filters out contigs that do not have at least 10x coverage on a total of 5 kbp across the whole contig. The read coverage filtering was done in this way to avoid filtering out long contigs with short read coverage patterns that might get removed if filtering was done with read coverage averages or medians. Additionally, TrIdent filters out contigs less than 30 kbp by default as they are often not long enough to capture complete transduction patterns. The contigs with sufficient VLP-fraction read coverage and length proceed to pattern-matching.

#### Pattern-matching algorithm

After filtering, the pileup files are re-formatted to increase the window size. While 100 bp windows provides the resolution needed to detect read coverage patterns associated with transduction events that only span a few thousand basepairs, like specialized transduction, read coverage patterns associated with other types of transduction are generally much larger and don’t require this level of resolution for detection. TrIdent uses 1 kbp windows by default to reduce the dataset size while preserving the read coverage patterns.

We broke down the transductomics classifications into their most basic characteristics-integrative MGEs, like prophages and chromosomal islands, are defined by ‘blocks’ of read coverage that represent the element’s reads mapping back to its integration site in the bacterial host genome. Generalized, lateral and GTA transduction events are defined by sloping read coverage patterns that correspond with the decreasing frequency of DNA packaging as the distances from the packaging initiation site increases. We designed two simple pattern classes to detect both types of read coverage patterns-a rectangle or ‘block’ pattern and a triangle or ‘sloping’ pattern. We also designed variations of these patterns to represent transduction events that are only partially captured on a single contig (i.e. patterns that ‘trail off the side’). Finally, we made a pattern which represents contigs with no read coverage patterns.

Next, we generated a metric to assess how well a specific pattern matched a contig’s read coverage. This metric, termed the ‘match-score’, is the mean absolute difference between the pattern values and the VLP-fraction read coverage values from a specific contig’s pileup file. If a pattern perfectly matches the read coverage values and positions on a contig, then the values will ‘zero-out’ if subtracted from each other. No predefined pattern will perfectly match the read coverage pattern associated with a transduction event, but the lower the match-score is to zero, the better the pattern-match. The patterns are formatted as if they were pileup files with columns for ‘coverage’ and ‘position’ values. The coverage values define the pattern shape, either Prophage-like, Sloping, or No Pattern and the positions replicate the window sizes used in the pileup file. The patterns are built using the specific length and VLP-fraction read coverage values of the contig being assessed. Certain pattern variations are translated across contigs in 1 kbp sliding windows and at each translation, a match-score is calculated. During translation, the overall patterns are kept the same length as the contig being assessed by simultaneously removing 1 kbp (equating to 1 row of the pileup file) from the ends of the pattern dataframes and adding 1 kbp to the beginnings. After translation is complete, certain aspects of the pattern are altered (i.e. height, width, slope). This process of translation and pattern re-scaling is repeated until a large number of pattern variations are tested. After pattern-matching is complete, the pattern class associated with the best (lowest) match-score is used for contig classification. Contigs are classified as ‘Prophage-like’, ‘Sloping’, or ‘NoPattern’ during pattern-matching.

#### Prophage-like patterns

There are three pattern variations in the prophage-like class. One variation represents an integrated MGE in which both borders of the element lie on the contig while the other two variations represent elements where one of the borders trails off the left or right side of the contig, respectively. The Prophage-like patterns contain a vector of repeated values that represent the elevated prophage-like region next to a vector of repeated values that represent the baseline region outside of the Prophage-like region. The value used for the baseline region of the pattern is defined by the minimum VLP-fraction read coverage value. The values iterated through for the elevated Prophage-like region are spaced at equal intervals between the contig’s maximum VLP-fraction read coverage value and one quarter of the maximum value. During each iteration, the Prophage-like pattern is first re-built so that the vector of values representing the elevated Prophage-like region spans almost the entire length of the contig being assessed and is then translated across the contig. After each translation across the contig, the Prophage-like region width is decreased by 1 kbp and the translation is repeated. The decrease in width and translation of the Prophage-like pattern is repeated until the minimum width, 10 kbp, is reached and the entire process is repeated with a new elevated Prophage-like region value. All Prophage-like patterns are translated across the contig being assessed. The 10 kbp minimum was chosen based on commonly used limits for viral genome identification in the literature^48–50^.

#### Sloping patterns

Generalized, lateral and GTA transduction events can span tens to hundreds of kbp and a single contig may not capture an entire event. We designed Sloping patterns that represent contigs that either contain or do not contain the start or packaging initiation associated with a sloping transducing event. The Slope patterns themselves consist of a sequence of numbers between the top and bottom slope values. Positive and negative sequences are generated for pattern variations that slope left and right, respectively, for a total of four Sloping pattern variations. The top and bottom values used to build the Sloping patterns are defined based on the read coverage values of the contig being assessed and are iterated over to change the pattern’s slope value. The values iterated through for the top of the Slope range from a value slightly greater than the maximum VLP-fraction read coverage to half of this value. The values iterated through for the bottom of the Slope range from the minimum VLP-fraction read coverage value to half of the maximum value. The pattern variations that contain the packaging initiation event have a short ‘baseline’ region of repeated values equal to the contig’s minimum VLP-fraction read coverage value added before the maximum value in the sloping sequence which creates the characteristic jump in read coverage that occurs at the start of some transduction events. During the iterative pattern-matching process, these pattern variations are first re-built so that the sequence of values representing the sloping region spans almost the entire length of the contig being assessed. They are then translated across the contig followed by having their width (the sequence of values representing the sloping region) decreased by 1 kbp. The decrease in width and translation of the sloping pattern is repeated until the minimum width, 20 kbp, is reached and the entire process is repeated with a new slope value. The 20 kbp minimum was chosen as it captures Sloping patterns that are only partially contained on the contig while limiting the misclassification of Prophage-like patterns with sloping elements. The pattern variations that do not contain the packaging initiation event only have the slope value changed during pattern-matching and are not translated or decreased in width. The iteration through top and bottom values breaks early if the pattern’s calculated slope value is less than 0.001 (change of 10x read coverage over 10 kbp).

#### No pattern

There are two noPattern pattern variations which consist of a horizontal line the same length as the contig being assessed at either the average or median read coverage for a contig. This pattern is not re-scaled or translated in any way.

### Final contig classification and characterization

#### NoPattern classifications with high VLP-fraction:whole-community read coverage ratios

If a contig receives a noPattern classification, it proceeds to an additional classification step which may re-classify the contig as having a high VLP-fraction:WC read coverage ratio (‘HighCovNoPattern’). Contigs with the HighCovNoPattern classification have even read coverage across the contig (i.e. no notable read coverage pattern) and the VLP-fraction read coverage is notably higher than the WC read coverage. Contigs with noPattern pattern-matches that have median 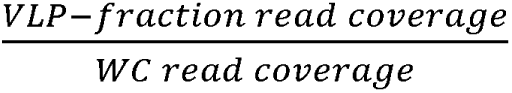 read coverage ratios greater than 2 are re-classified as HighCovNoPattern by default. Any bacterial WC contig covered by VLP reads, regardless of WC coverage, is theoretically the result of transduction since the assumption is that all DNA in the VLP-fraction is contained within VLPs. The default 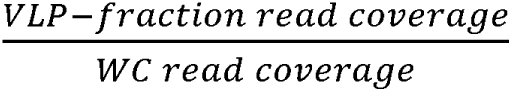 ratio of 2 is arbitrary and simply indicates that the contig has double the coverage in the VLP-fraction compared to the WC and may therefore be a frequent transducer. However, the ratio can be changed based on the user’s preferences and/or can be made relative to the number of sequencing reads obtained for the WC and VLP-fraction.

### TrIdent performance testing

#### Original transductomics data generation

The original transductomics data TrIdent was initially compared against was generated in Kleiner et al (2020). In brief, the colon contents from one mouse was homogenized and split into two aliquots for WC and VLP-fraction DNA extraction. The aliquot being used as the WC fraction was treated with lysozyme and bead beat prior to DNA extraction with phenol:chloroform and ethanol precipitations. The aliquot being used as the VLP-fraction was centrifuged to pellet debris and the supernatant was 0.45 um filtered. The filtered supernatant was treated with DNase and RNase before being loaded onto a three layer CsCl density gradient and ultracentrifuged overnight. The band between the 1.35 and 1.5 g/ml density layers was extracted and buffer exchanged into SM-buffer using centrifugal filters (10 kDa MWCO). The buffer-exchanged sample was treated with Proteinase K and SDS to disrupt VLPs and release enclosed nucleic acid. The DNA was extracted with phenol:chloroform and ethanol precipitation. Sequencing libraries were prepared with the WC and VLP-fraction DNA samples and the libraries were sequenced with Illumina. The WC-fraction was sequenced in two separate runs-the first in paired-end mode with 76M 150 bp reads and the second in paired-end mode with 313M 75 bp reads. The VLP-fraction was also sequenced in two separate runs-the first in paired-end mode with 97M 150 bp reads and the second in paired-end mode with 262M 75 bp reads. The WC sequencing reads were decontaminated of reads that mapped to the mouse, human and Illumina control phiX reference genomes using BBSplit. The unmapped reads were trimmed for quality and adapter removal using BBDuk. The cleaned reads were assembled with SPades and the assemblies were filtered for contigs > 30 kbp. The WC and VLP-fraction reads were mapped onto the WC assembled contigs using BBMap. The resulting bam files were sorted and indexed with SAMtools. The sorted and indexed bam files were used to create pileup files with BBMap’s pileup.sh using binsize=100.

#### Re-labeling original transductomics data

The classifications given to the original transductomics dataset had to be cleaned for direct comparison to the TrIdent classification labels. So as not to bias the new classifications, we did not re-inspect the coverage patterns prior to re-labeling and instead derived a set of criteria for consistent re-labeling. Any label which contained the word(s) ‘prophage’, ‘chromosomal island’, ‘retrotransposon/transposon/transposase/transposable element’, or ‘specialized’ were re-labeled as ‘Prophage-like’. Any label which contained the word(s) ‘gene transfer agent’, ‘plasmid’, or ‘bacterium’ were re-labeled as ‘HighCovNoPattern’ and labels which contained the word(s) ‘generalized’ were re-labeled as ‘sloping’. Finally, labels which contained the word(s) ‘false hit’ or ‘none’ were re-labeled as ‘NoPattern’ and labels which contained the word(s) ‘weird’, ‘unknown’, ‘mixed’ or contained the word(s) associated with several other categories were re-labeled as ‘unknown’ (S1 table). The pileup files described above were imported into R where they were processed with TrIdent’s TrIdentClassifier() function with default parameters.

#### Comparing TrIdent to manual classifiers

To assess the efficacy of TrIdent when compared to the industry standard of manual classification of transductomics datasets, example datasets were classified automatically by TrIdent and manually by two independent experts. Over the course of 5x 2-hour classroom sessions, each expert received extensive training in phage transduction, metagenomics and transductomics data analysis prior to performing the final classifications. Once the experts had a sufficient knowledge base to classify transductomics data, they were given a practice dataset to work through with the trainer. After it was confirmed that the experts’ classifications on the practice dataset were sufficient, they were given the final transductomics datasets to be used for benchmarking. Each classifier was provided with three transductomics datasets from experimental replicate 2 across each of the three sampling conditions described in the case study. The case study data had not been analyzed with TrIdent prior to training or data provision. Each classifier was instructed to use predefined labels for their classifications (‘prophage-like’, ‘sloping’, ‘no pattern’, ‘high coverage no pattern’, and ‘unknown’), work independently, and to keep track of their time spent classifying.

We merged the classification results for CL1, CL2 and TrIdent in R and compared the classifications with plots generated using the ggplot2 and ggVennDiagram R packages. We calculated percent agreement between all three classifiers by dividing the number of contigs in which the classifiers made the same classification by the total number of contigs in each dataset. We considered each of the three datasets a replicate and compared the agreement percent for each dataset between all three classifiers, CL1 and CL2, CL1 and TrIdent, and CL2 and TrIdent. We ran a paired Wilcoxon test to determine if there were statistically significant differences (p<0.05) between the percent agreement for each classifier comparison group. We also calculated unweighted Cohen’s Kappa scores between each of the classifiers to determine if there were differences in scores between comparison groups.

#### Transductomics experiment used to compare TrIdent to manual classifiers and to exemplify TrIdent’s use in the case study

##### Animals and Housing

C57BL/6J mice (male, 4-5 weeks old) purchased from Jackson Laboratories were used for the experimental infections. Mice were housed with autoclaved bedding and water and irradiated food. Cage changes were performed weekly in a laminar flow hood. All mice were subjected to a 12-hour light and 12-hour dark cycle. Animal experiments were conducted in the Laboratory Animal Facilities located on the NCSU CVM campus. The animal facilities are equipped with a full-time animal care staff coordinated by the Laboratory Animal Resources (LAR) division at NCSU. The NCSU CVM is accredited by the Association for the Assessment and Accreditation of Laboratory Animal Care International (AAALAC). Trained animal handlers in the facility fed and assessed the status of animals several times per day. Those assessed as moribund were humanely euthanized by CO_2_ asphyxiation. This protocol is approved by NC State’s Institutional Animal Care and Use Committee (IACUC).

##### Sampling

We generated transductomics datasets from murine fecal samples collected during an experiment in which mice (n=4) were treated with antibiotics prior to challenge with *C. difficile*^36^. We collected fecal pellets from the mice immediately before and after an antibiotic treatment in which cefoperazone (0.5 g/mL) was provided in the drinking water for 5 days. After 2 days of antibiotic-washout with fresh drinking water, the mice were orally gavaged with ∼1.0e5 *C. difficile* (strain 630) at which point we started monitoring the mice for clinical signs of *C. difficile* infection (CDI). On day 4 of CDI, when toxin mediated inflammation is severe, we collected a final round of fecal pellets from all 4 mice. Each sample collection consisted of two fecal pellets from each mouse replicate. The pellets were manually homogenized in 1.2 mL of SM buffer (100mM NaCl, 8mM MgSO4-7H2O, 50mM TrisHCl) using sterile inoculating loops. After homogenization, the samples were brought up to 2 mL total volume using SM buffer. The fecal homogenates were split into two aliquots, 1.5 mL and 0.5 mL, for WC and VLP-fraction DNA extraction, respectively. Each of the VLP-fractions was further split into two 0.75mL aliquots and one of the aliquots from each sample was processed and purified with CsCl density gradient ultrancentrifugation immediately. The WC samples were stored at-80C until DNA extraction.

##### Whole-community DNA extraction

The DNA from the WC aliquots was extracted with the Qiagen PowerFecal Pro kit according to the manufacturer’s instructions. We followed the manufacturer’s recommended modifications for cells that are difficult to lyse. Unless specified, all centrifugation steps were carried out at 15,000 xG for 1 minute. Briefly, we added the homogenized stool samples to the provided bead beat tubes, added 800uL of CD1 buffer and briefly vortexed to mix. The samples were incubated at 65C for 10 minutes before being vortexed at max speed for 10 minutes. The tubes were centrifuged and the supernatant was transferred to a clean tube. We added 200uL of CD2 buffer and briefly vortexed before centrifuging and transferring supernatant to a clean tube. Next, we added 600uL of CD3 buffer and briefly vortexed before loading the sample on to the provided MB column spin tube and centrifuging. After the entire sample was loaded, the flow through was discarded and the column was placed into a clean collection tube. We added 500uL of EA to the spin tubes and centrifuged before discarding the flow-through. Then, we added 500uL of C5 buffer to the columns, centrifuged, and then placed the column into a clean collection tube where it was centrifuged again at 16,000 xG for 2 minutes. The columns were moved to clean collection tubes where we added 50uL of C6 buffer to the spin columns and centrifuged. The final extracted DNA product was quantified with Qubit dsDNA HS assay and stored at-80C before being sent for sequencing.

##### VLP-fraction purification

The VLP-fractions were centrifuged at 2,500 xG for 5 minutes to pellet large debris. The supernatant was removed and centrifuged for 5,000 xG for 5 minutes to pellet any remaining debris. The supernatant was 0.2 um filtered and treated with 100 U of DNase 1 (Thermo Scientific, catalog no. PI90083) and incubated at 37 C for 2 hours. We set up density-gradients using 1.7, 1.5, and 1.35 g/mL cesium chloride (CsCl) density layers prepared in SM buffer in Beckman ultra-clear tubes (14×89 mm, catalog no. 344059) and marked the interface between the 1.5 and 1.35 g/mL density layers. We loaded the digested VLP-fractions on top of the gradients and ultracentrifuged the tubes at 68,600 xG for 24 hours at 4C. After centrifugation, there was no visible band in the density gradients so we used a 27 gauge needle to puncture the tubes ∼2mm below the marked interface and extracted ∼3mL of liquid. We used a 10 Kda Amicon centrifugal filter to wash the purified VLPs 3x with SM buffer and concentrate the samples down to ∼500 uL each.

##### VLP-fraction DNA extraction

To extract DNA from purified VLPs, we performed a proteinase K digestion followed by a phenol:chloroform extraction and DNA concentration with MinElute tubes. First, the VLP samples were lyophilized overnight followed by resuspension in 100uL of PCR-grade water. Next, the resuspended VLPs were combined 1:1 with lysis mix (2% SDS, 90 ug/mL Proteinase K (Thermo Scientific, catalog no. FEREO0491)) and incubated at 55C for 1 hour. The samples were cooled to room temperature before being transferred to pre-spun 5prime light phase-lock tubes. We added 500uL of phenol:chloroform:isamyl (25:24:1) and inverted the tubes to mix before centrifuging at 12,000 xG for 5 minutes. The aqueous phase was poured into a new pre-spun phase lock tube where we added 500 uL of chloroform:isoamyl (24:1) and inverted to mix before centrifuging at 12,000 xG for 5 minutes. We poured the aqueous phase into a clean microcentrifuge tube where we used the MinElute PCR Purification kit according to the manufacturer’s instructions to clean and concentrate the extracted DNA. Briefly, we diluted the DNA samples 5:1 in buffer PB and applied the diluted sample to a MinElute spin column and centrifuged at 17,900 xG for 1 minute. After the samples were completely loaded onto the columns, they were washed with 750uL of buffer PE and centrifuged at 17,900 xg for 2 minutes. The columns were placed into clean collection tubes where 20uL of buffer EV was added to the column membranes. The columns were allowed to sit for 1 minute prior to centrifuging at 17,900 xG for 1 minute. The final DNA product was quantified with the Qubit dsDNA HS assay and stored at-80C before sequencing.

##### Sequencing for transductomics analysis

Our final sample collection consists of a WC and associated VLP-fraction from 4 mice under three different sample conditions (Pre-ABX, Post-ABX, CDI) thus generating a total of 24 samples that were sequenced with the Illumina NovaSeq X Plus platform at 2×150bp. We received between 50-160M reads for the WC samples and between 35-180M reads for the VLP-fractions. We used TrimGalore^51^ to trim the raw reads of Illumina adaptors and polyG sequences identified through FastQC. Next, we used BBSplit with ambig2=split to remove reads that mapped to the host mouse (GCF_000001635.27) or PhiX (NCBI: NC_001422.1) genome. While the WC of replicate 4 sequenced well (Table S2) and was used for binning and taxonomic classification of the WC, the VLP-fraction lost the majority of sequencing reads after quality filtering and contaminant removal and was therefore excluded from the analysis (Table S2).

##### Generation and taxonomic classification of whole-community metagenome assembled genomes (MAGs)

We assembled the WC metagenomes using MEGAHIT^52^ with default parameters and ran geNomad’s^39^ end-to-end pipeline on each of the Pre-ABX assemblies for virus and plasmid identification. We mapped all the trimmed and decontaminated WC reads from each replicate in all conditions to each of the WC assemblies using BBMap with default parameters. The resulting.bam files were sorted with SAMtools^53^ then binned with MetaBAT2’s^54^ runMetaBat using default parameters. We performed taxonomic classification of the MAGs using GTDB-Tk’s^55^ classify_wf with the skip-ani-screen parameter.

##### Relative abundance calculation for whole-communities and transductomes

We mapped WC and VLP-fraction reads to the MAGs from the respective WC sample using BBMap (ambiguous=random, minid=0.97). To obtain the relative abundance of each MAG in the Pre-ABX WCs and transductomes, we summed the reads by MAG for each WC and VLP metagenome and divided by the total number of trimmed and decontaminated reads for the respective sample. MAGs with less than 0.05% relative abundance in a given sample were removed to avoid spurious MAG detections that could be derived from incorrect read mapping. We removed viral and prophage-containing contigs (classified with geNomad) prior to determining relative abundances.

##### Transductomics for TrIdent performance analysis

We filtered the assemblies generated from the second mouse replicate for contigs greater than 30 kbp and mapped both the WC and VLP-fraction sequencing reads to the filtered assemblies using BBMap with ambiguous=random and minid=0.97. We generated pileup files from the resulting.bam files using BBMap’s pileup.sh command with binsize=100. The pileup files were imported into R where they were processed with TrIdent’s TrIdentClassifier() function with default parameters. The.bam files were also sorted and indexed with SAMtools sort and index functions, respectively, and converted to.tdf files with Igvtools using the count command, the mean window function and window size=100. The.tdf files were used for read coverage visualization in IGV by the manual classifiers.

##### Taxonomy and abundance of transduction events in the transductomes

To determine taxonomy of contigs classified by TrIdent, we ran CAT^56^ with the GTDB-Tk database to assign taxonomy to any contigs that had not already been binned when determining the taxonomy of the WC (described above). We determined transduction event abundance by extracting the maximum pattern-match values for each of the classified contigs. These values are stored in TrIdent’s output list. We normalized the maximum pattern-match values by the number of mapping reads from each of the respective VLP-fraction samples to obtain a final abundance value.

## List of Abbreviations

HGT: Horizontal Gene Transfer
VLP: Virus-Like Particle
WC: Whole-Community
MV: Membrane Vesicle
GTA: Gene Transfer Agent
ABX: Antibiotics
CDI: *C. difficile* Infection
CL: Classifier

## Availability and requirements

**Project name:** TrIdent

**Project home page:** https://bioconductor.org/packages/release/bioc/html/TrIdent.html

**Operating system(s):** Platform independent

**Programming language:** R

**Other requirements:** None

**License:** GPL-2

## Declarations

### Ethics approval and consent to participate

Not applicable

### Consent for publication

Not applicable

### Availability of data and materials

The raw sequence datasets generated for the benchmarking of TrIdent against human classifiers and the transductomics case study are available in the Sequence Read Archive (SRA) repository under BioProject PRJNA1404639. The assemblies and MAGs used to generate taxonomy are deposited in Dryad (DOI: 10.5061/dryad.76hdr7t9b). The data analyzed during the comparison of TrIdent to a previously generated transductomics dataset is included in the 2020 Kleiner et al. article [and its supplementary information files].

### Competing interests

The authors declare that they have no competing interests.

### Funding

This research was supported by a seed grant from the North Carolina State University Data Science Academy and by the National Institutes of Health under Award Numbers R35GM138362 and R01Al171046.

### Authors’ contributions

**Jessie Maier:** Conceptualization, Data Curation, Formal Analysis, Investigation, Methodology, Software, Visualization, Writing-original draft, Writing-editing and review

**Craig Gin:** Conceptualization, Software

**Jorden Rabasco:** Software

**Wynter Spencer:** Data Curation

**Avery Bass:** Data Curation

**Breck A. Durekop:** Conceptualization, Funding Acquisition, Writing-editing and review

**Ben Callahan:** Conceptualization, Software, Funding Acquisition, Writing-editing and review

**Manuel Kleiner:** Conceptualization, Software, Funding Acquisition, Resources, Writing-editing and review

## Acknowledgements

We would like to thank Casey Theriot and Cypress Perkins for their resources and assistance with running the CDI murine model. We would also like to thank all Kleiner Lab members for their consistent support and feedback throughout this research work.

## Notes

https://www.ncbi.nlm.nih.gov/bioproject/?term=(PRJNA1404639)%20AND%20bioproject_sra[filter]%20NOT%20bioproject_gap[filter]

https://datadryad.org/dataset/doi:10.5061/dryad.76hdr7t9b

### Competing Interest Statement

The authors have declared no competing interest.

### Summary of Updates

We changed the relative abundance calculation used to determine WC and transductome composition. Figure 5, results and methods text have been updated to reflect the changes.

