## Supplemental Figures for "TrIdent - An R package to automate transductomics analysis of virus-like particle mediated DNA mobilization"

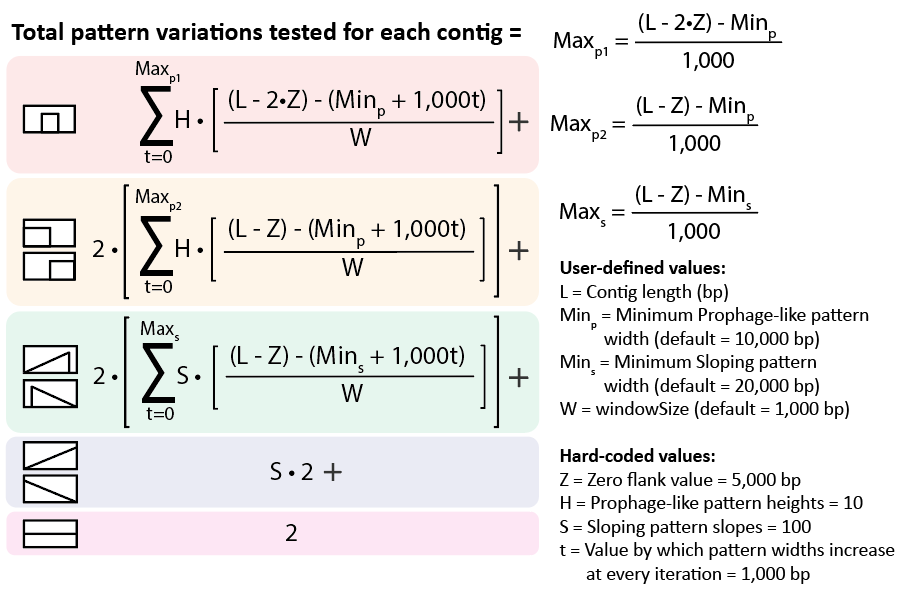

**Supplementary figure 1- Formula to calculate number of pattern variations tested against each contig in a dataset.** Patterns that are translated across the contig require summation (**Σ)** in which a series of possible pattern widths are iterated through from minimum (Min_s_, Min_p_) to maximum (Max_s_, Max_p1_, Max_p2_) increasing by 1,000 bp at each iteration (t). At each iteration, the number of possible pattern translations is calculated for the given pattern width, taking into account the size of the pattern’s zero flank regions (Z) (i.e. the regions flanking the patterns that have a value of 0 or close to 0), the minimum pattern width (Min_s_, Min_p_), the contig length (L) and the windowSize used for pattern translation (W). This value is multiplied by the number of Prophage-like pattern height (H) or Sloping pattern slope (S) values tested to determine the total pattern variations tested for the respective pattern. Patterns that are not translated do not require summation. The number of total pattern-variations tested for a contig may differ if non-default values are used (i.e. if the minimum Prophage-like pattern width is set to 50,000 bp, a 40,000 bp contig will generate no Prophage-like pattern variations). Contigs must be greater than 25,000 bp for pattern-matching.

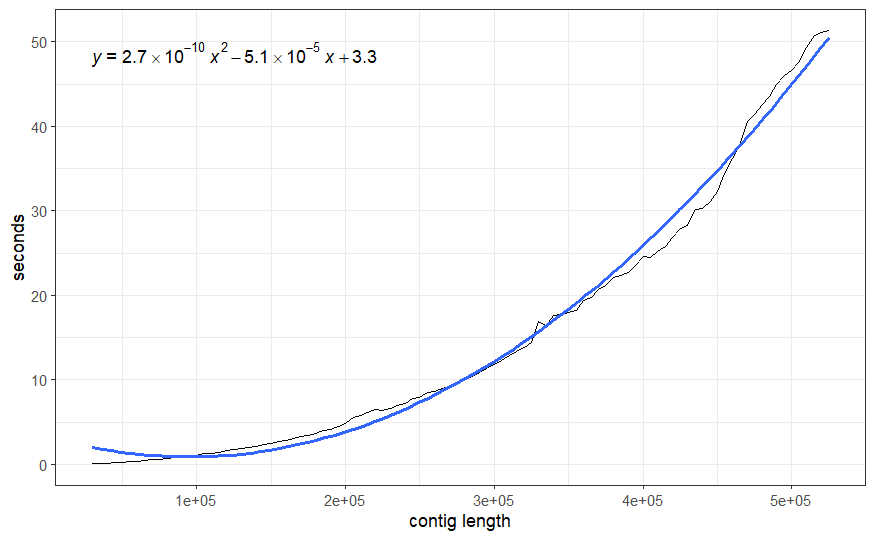

**Supplementary figure 2- Average (n=3) computational time for pattern-matching and classification at increasing contig length with default TrIdentClassifier() parameters.** Run on a system with an 11th Gen Intel(R) Core(TM) i7-1165G7 processor @ 2.80Ghz with 32 GB of RAM (3200 MT/s). Average computational time is plotted in black and a quadratic regression line fit to the data is overlaid in blue. The equation for the regression line is displayed in the top left: $y=2.7*10^{-10}x^{2}-5.1*10^{-5}x+3.3$.

**Note- multipage figure, caption below**
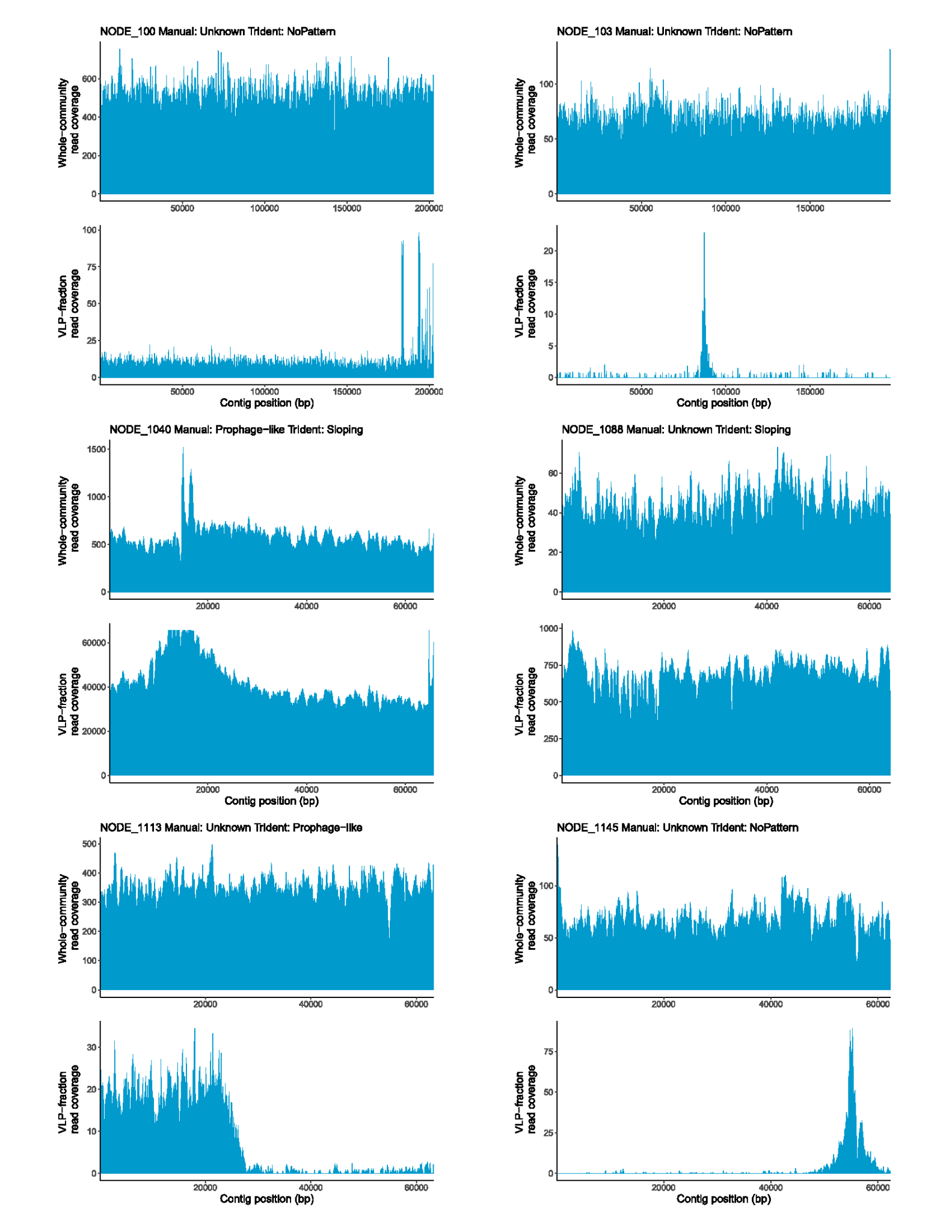

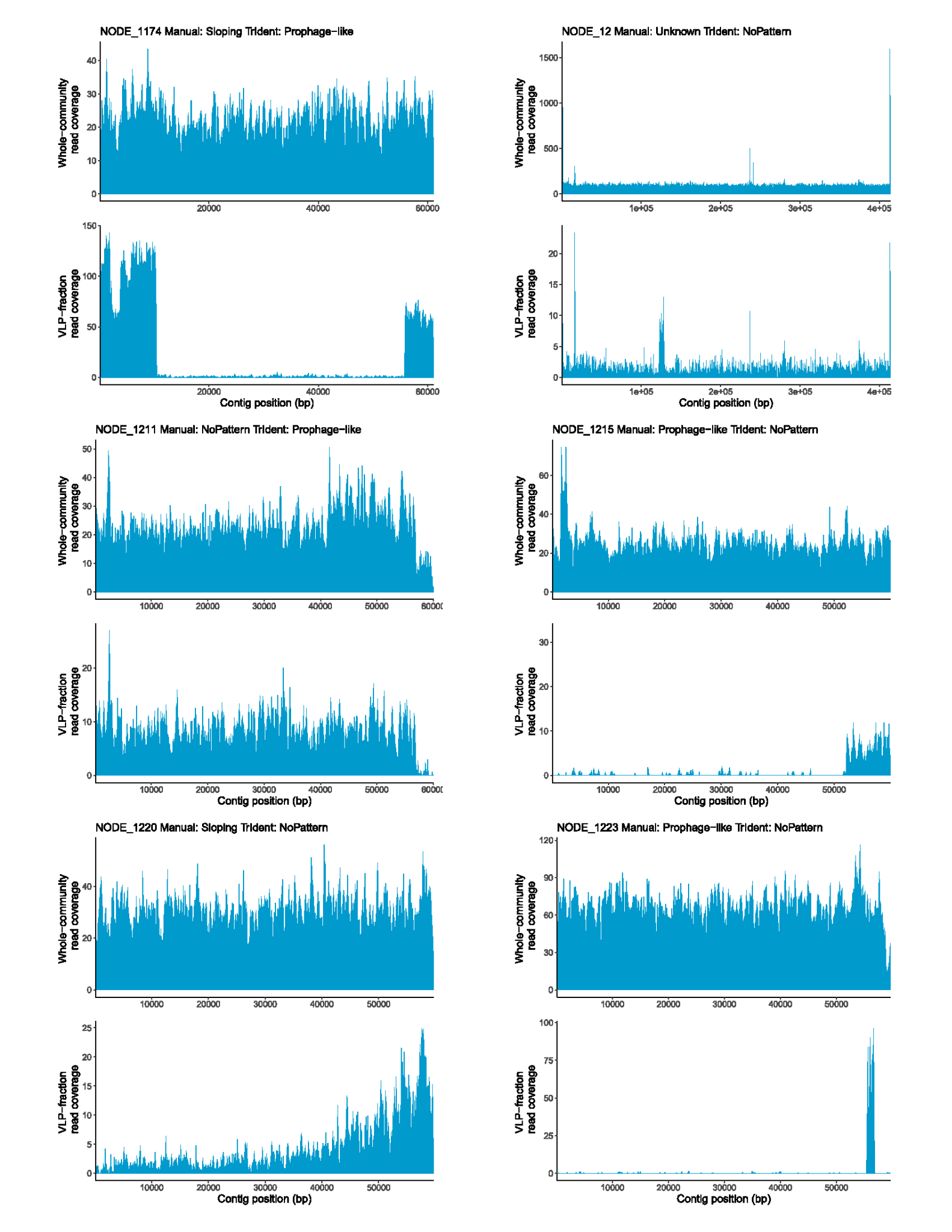

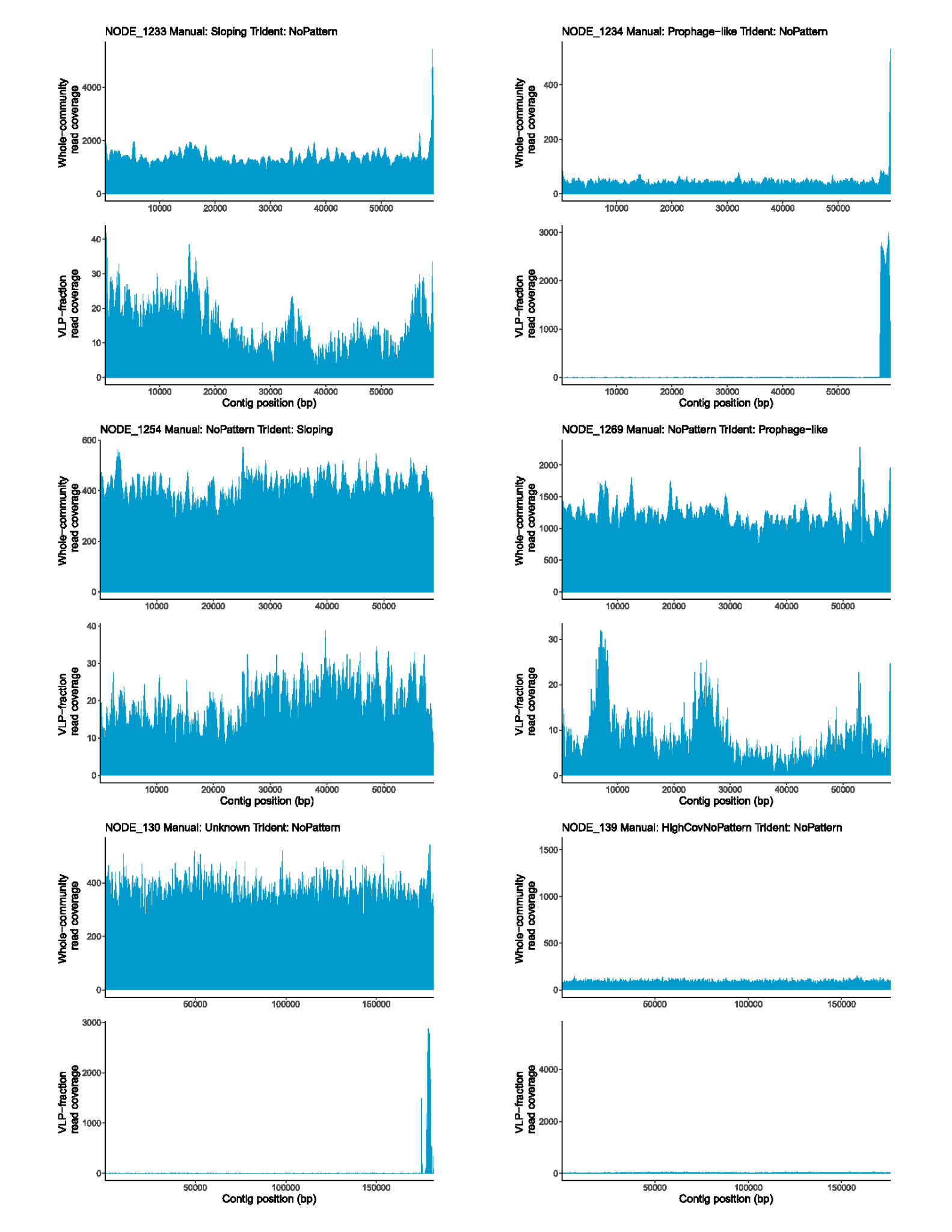

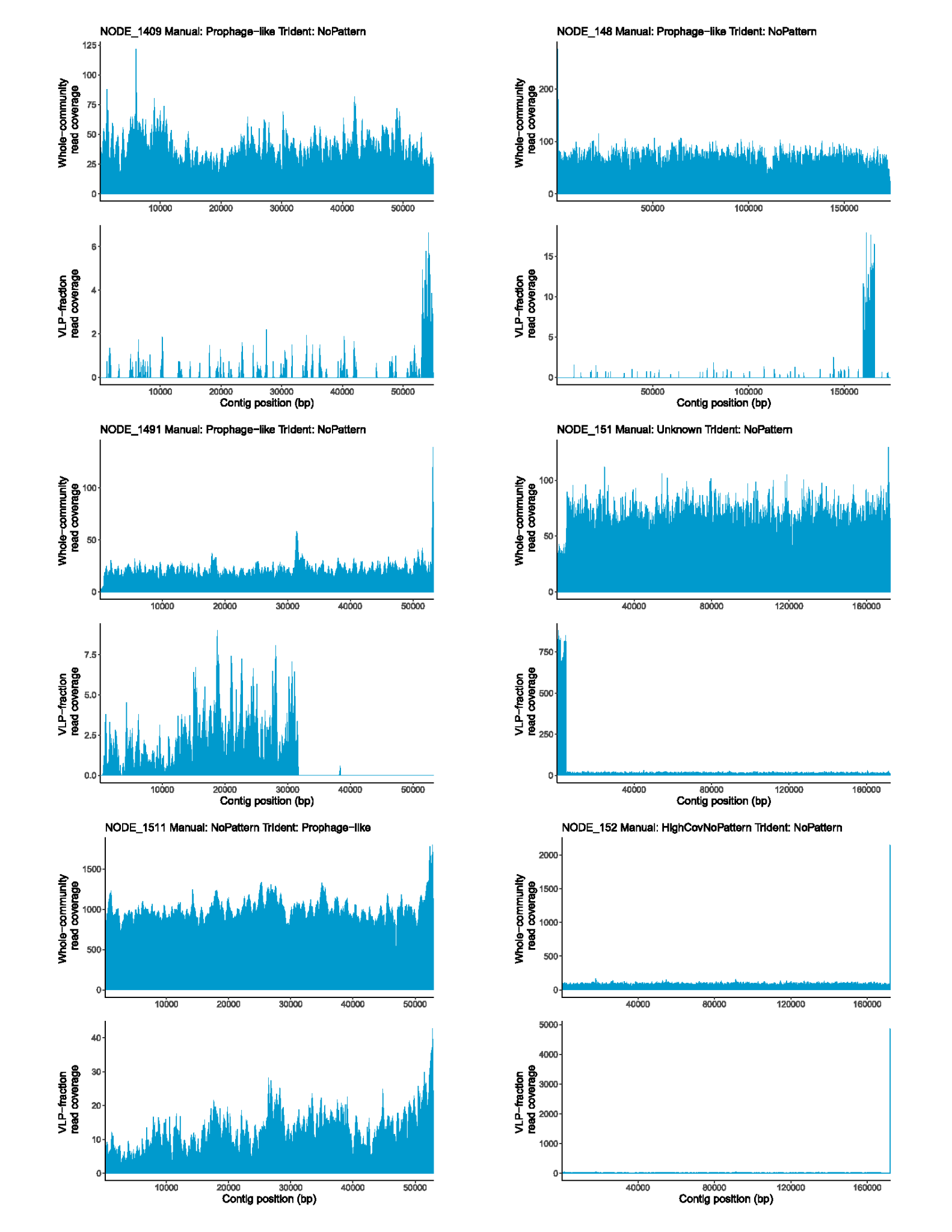

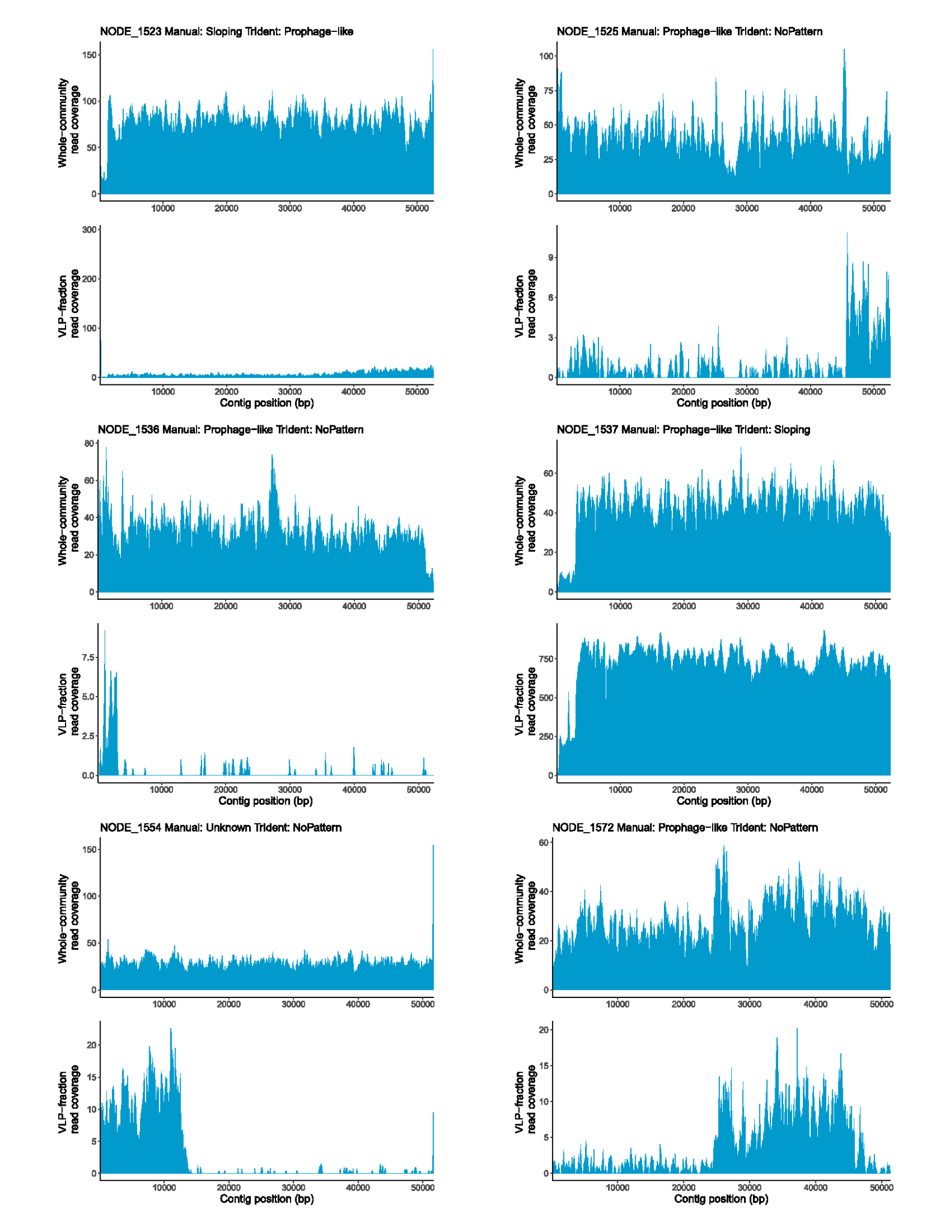

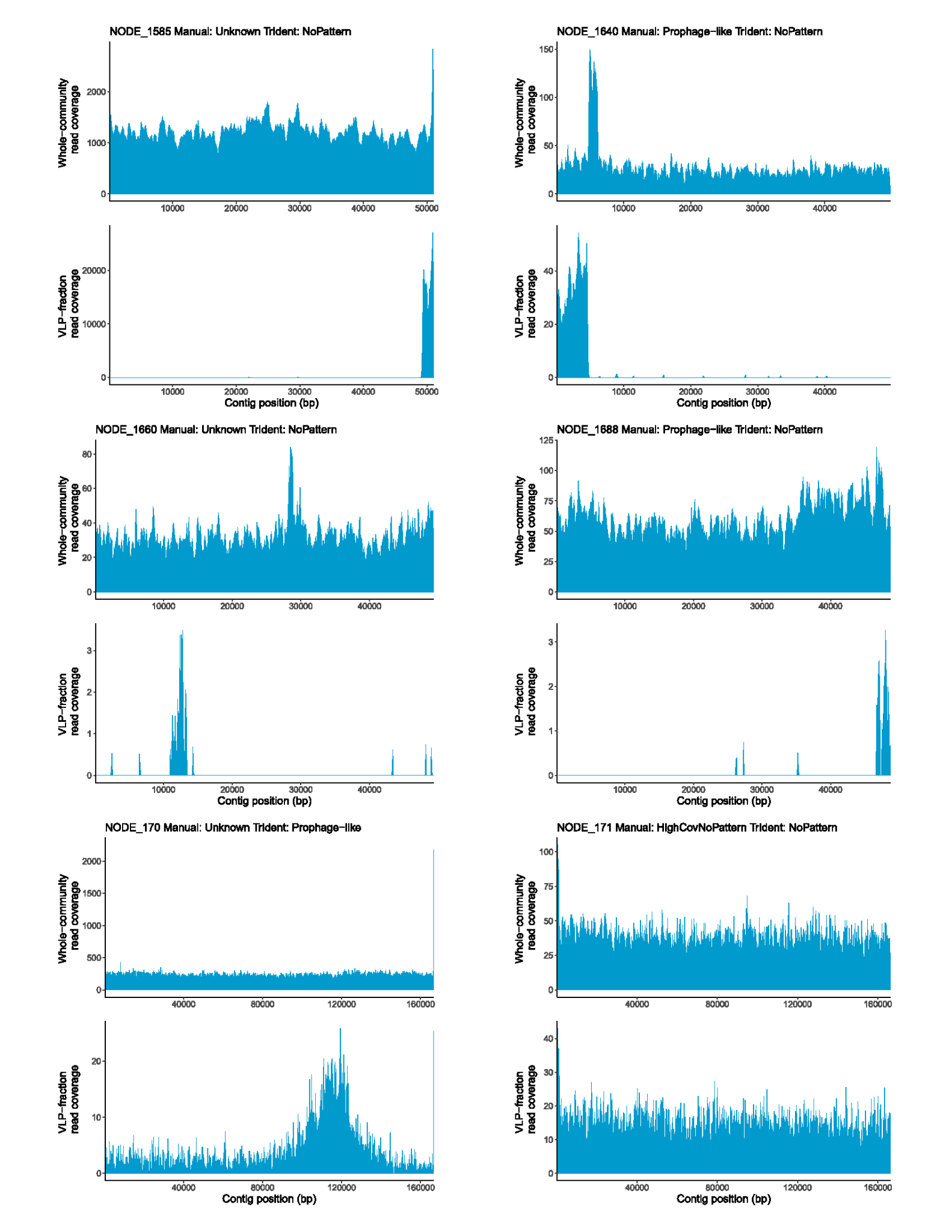

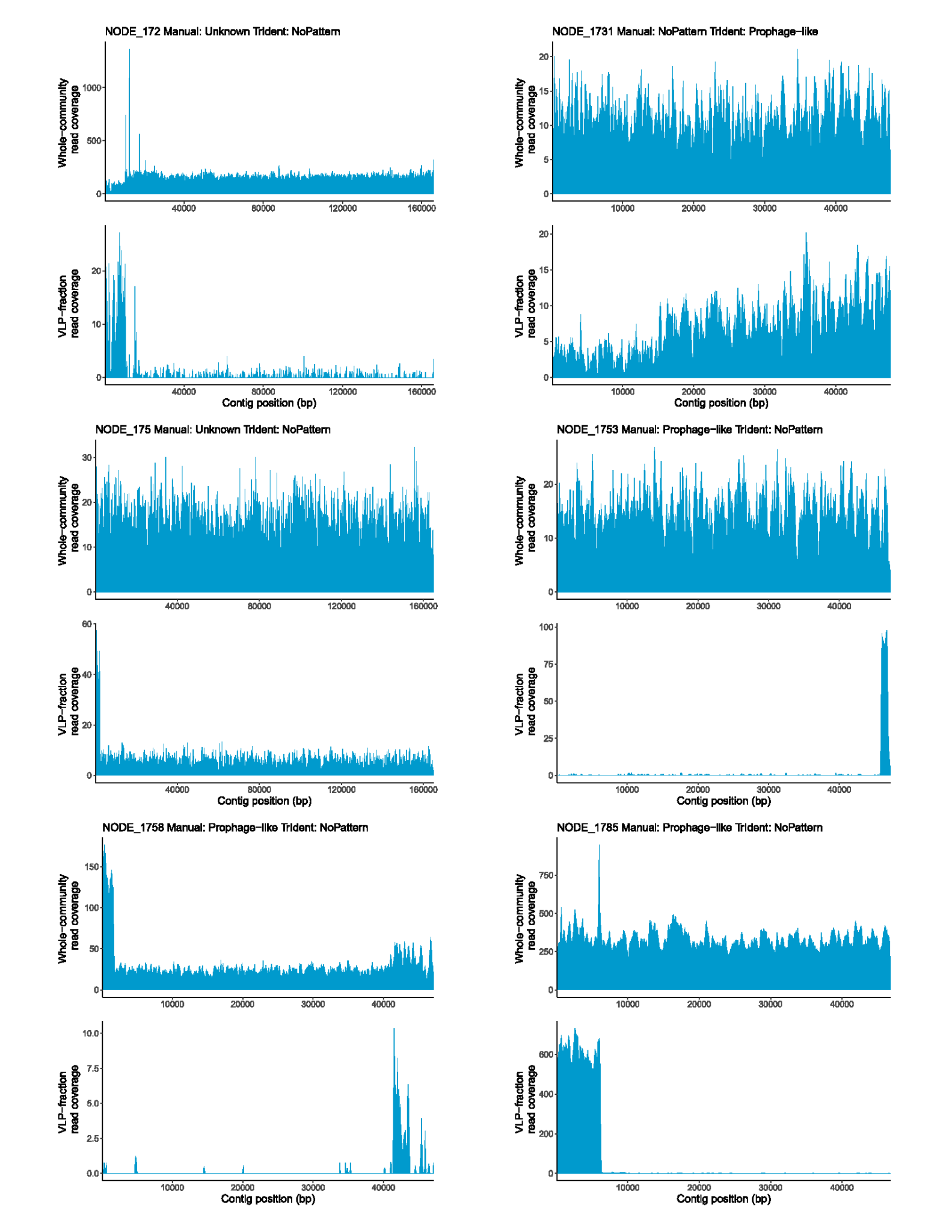

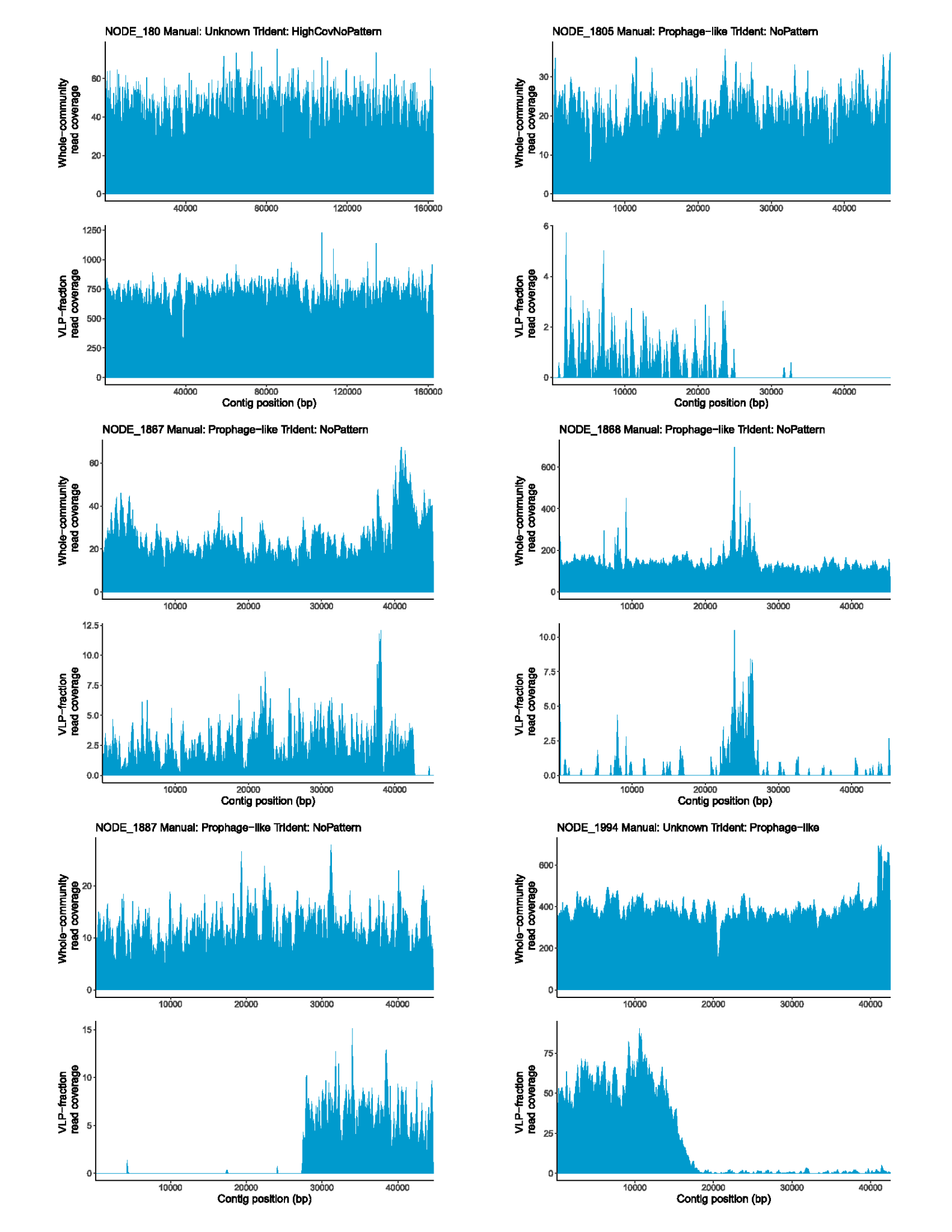

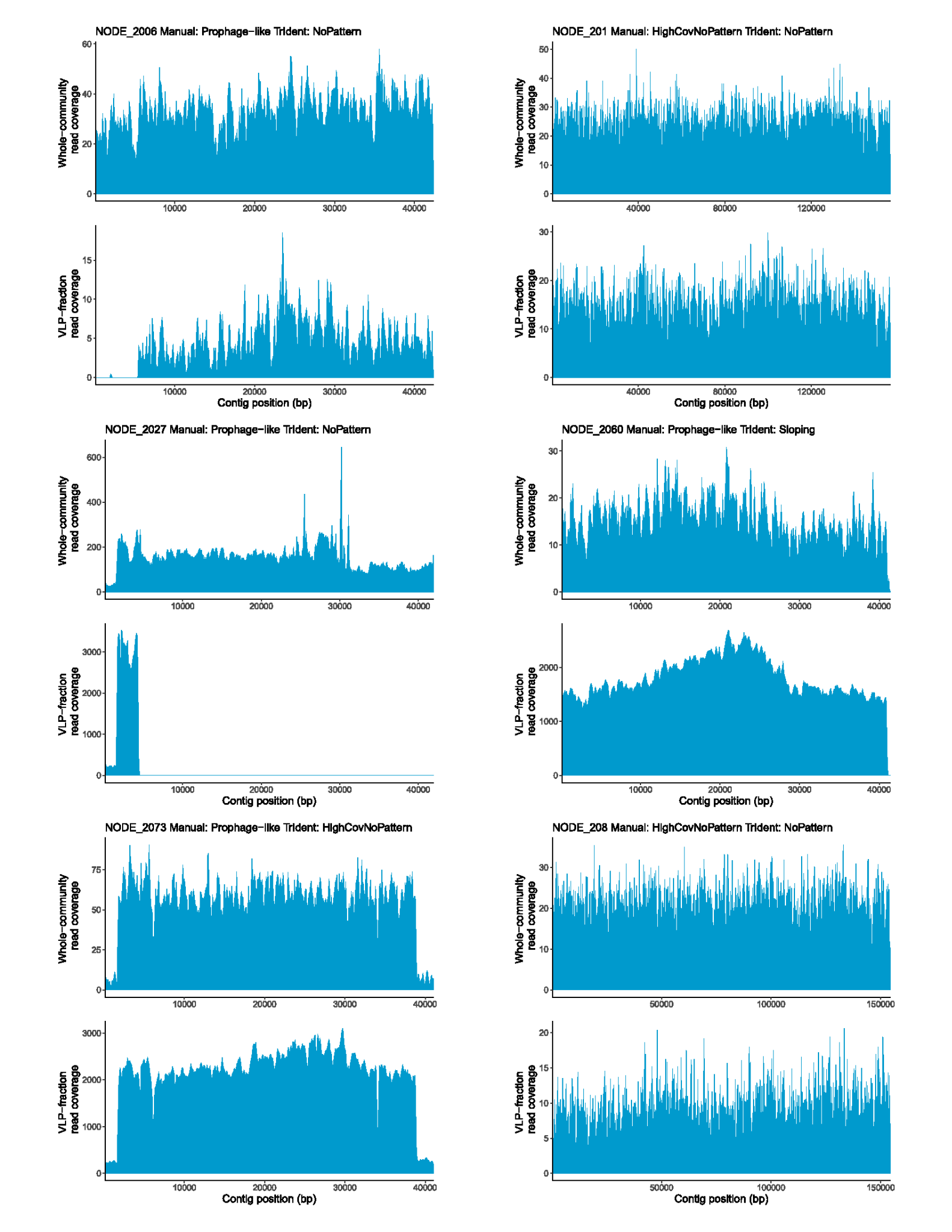

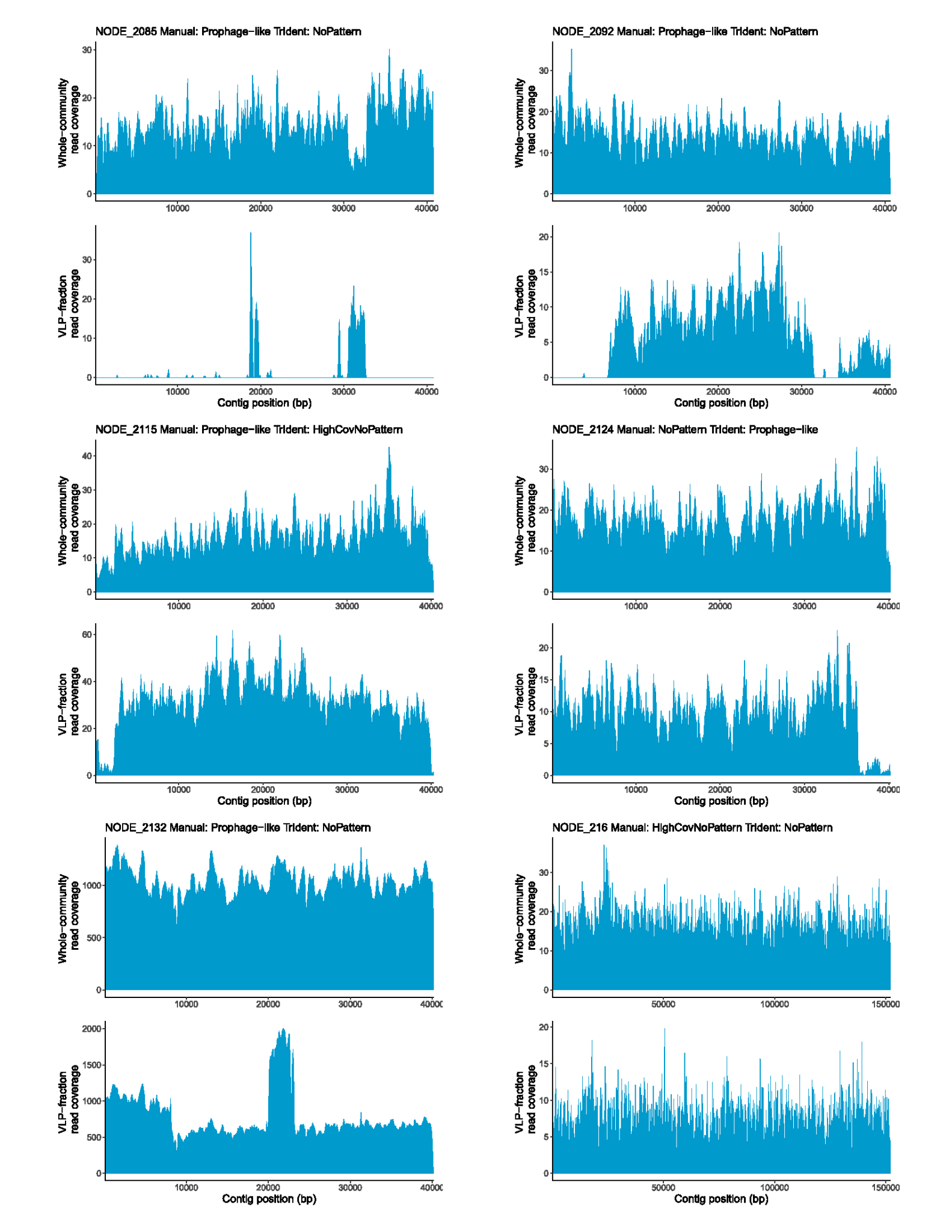

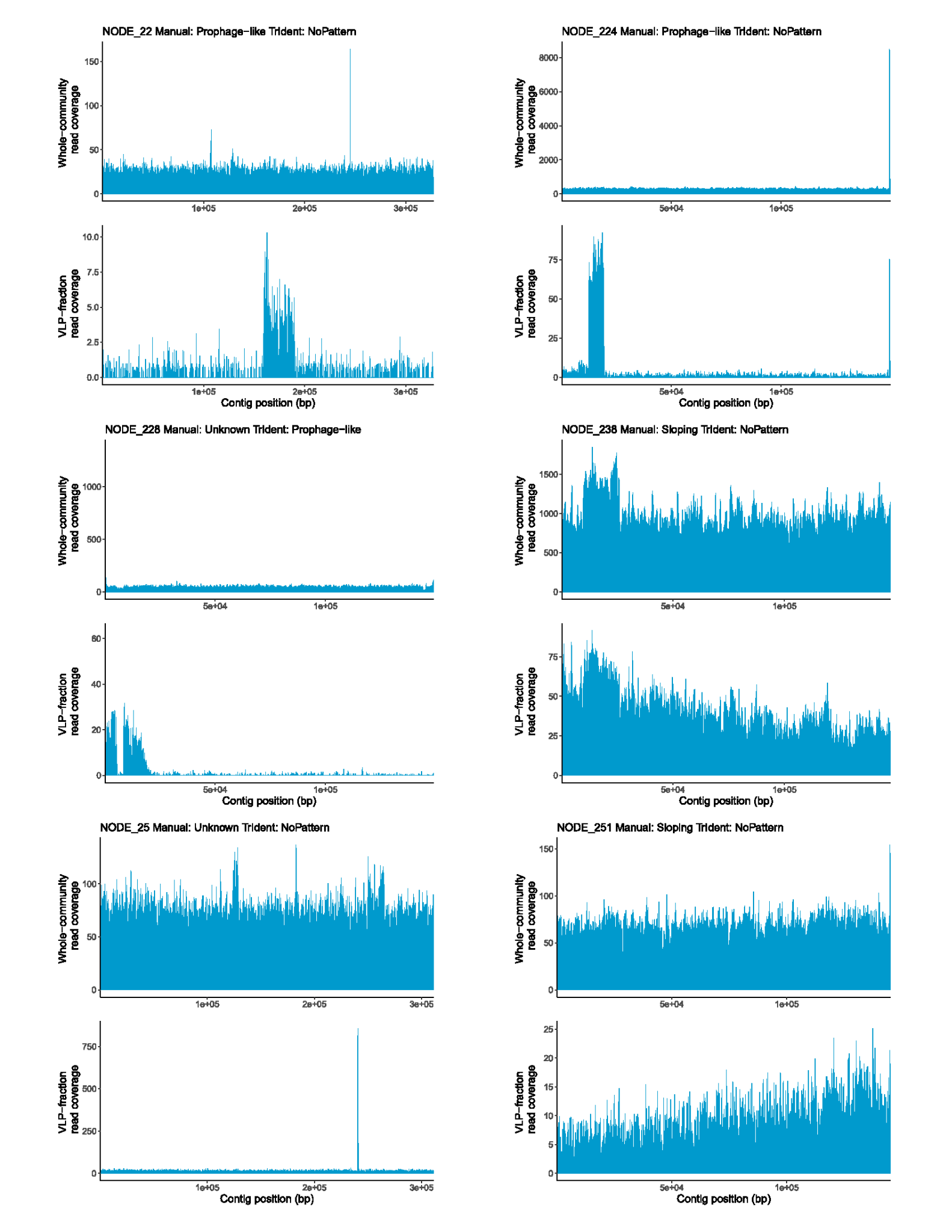

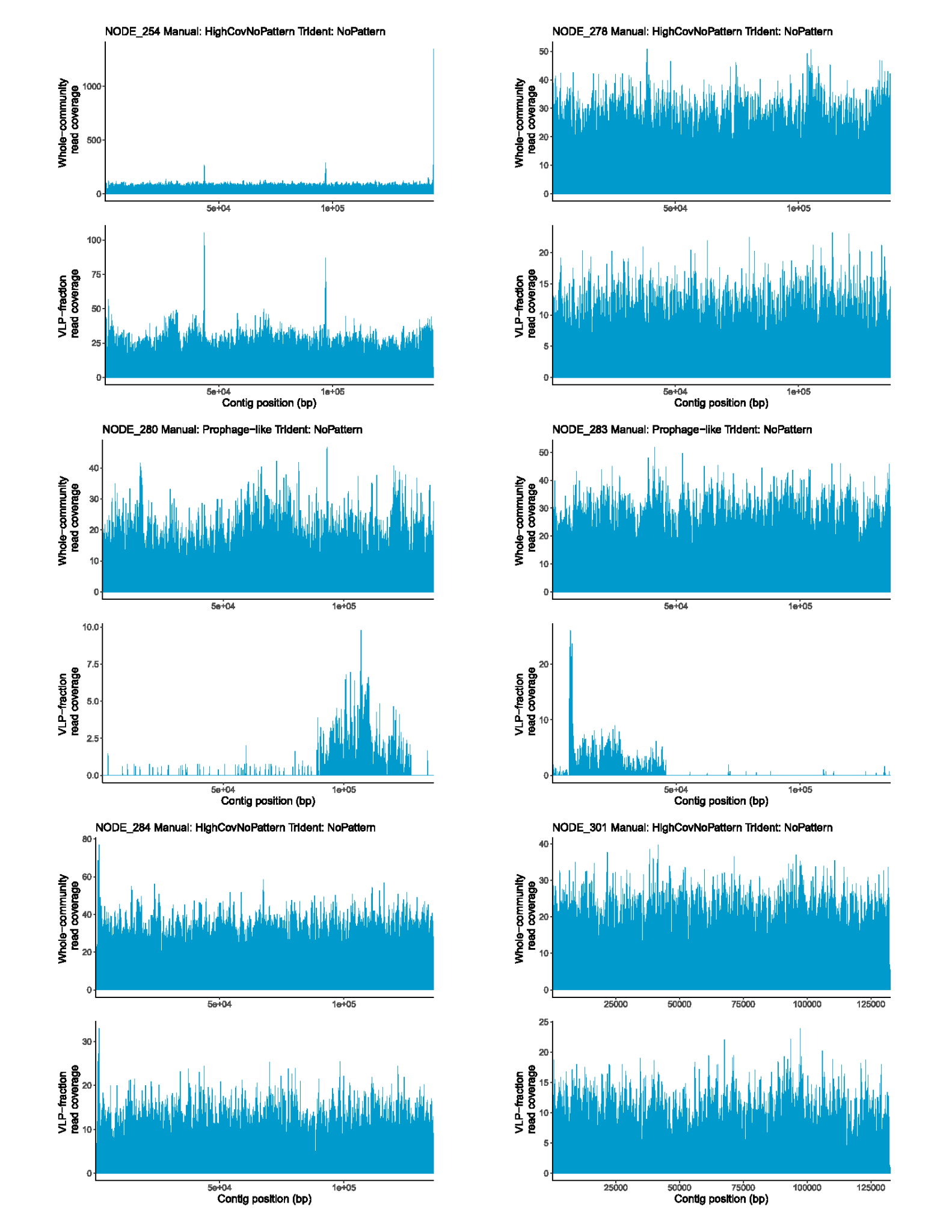

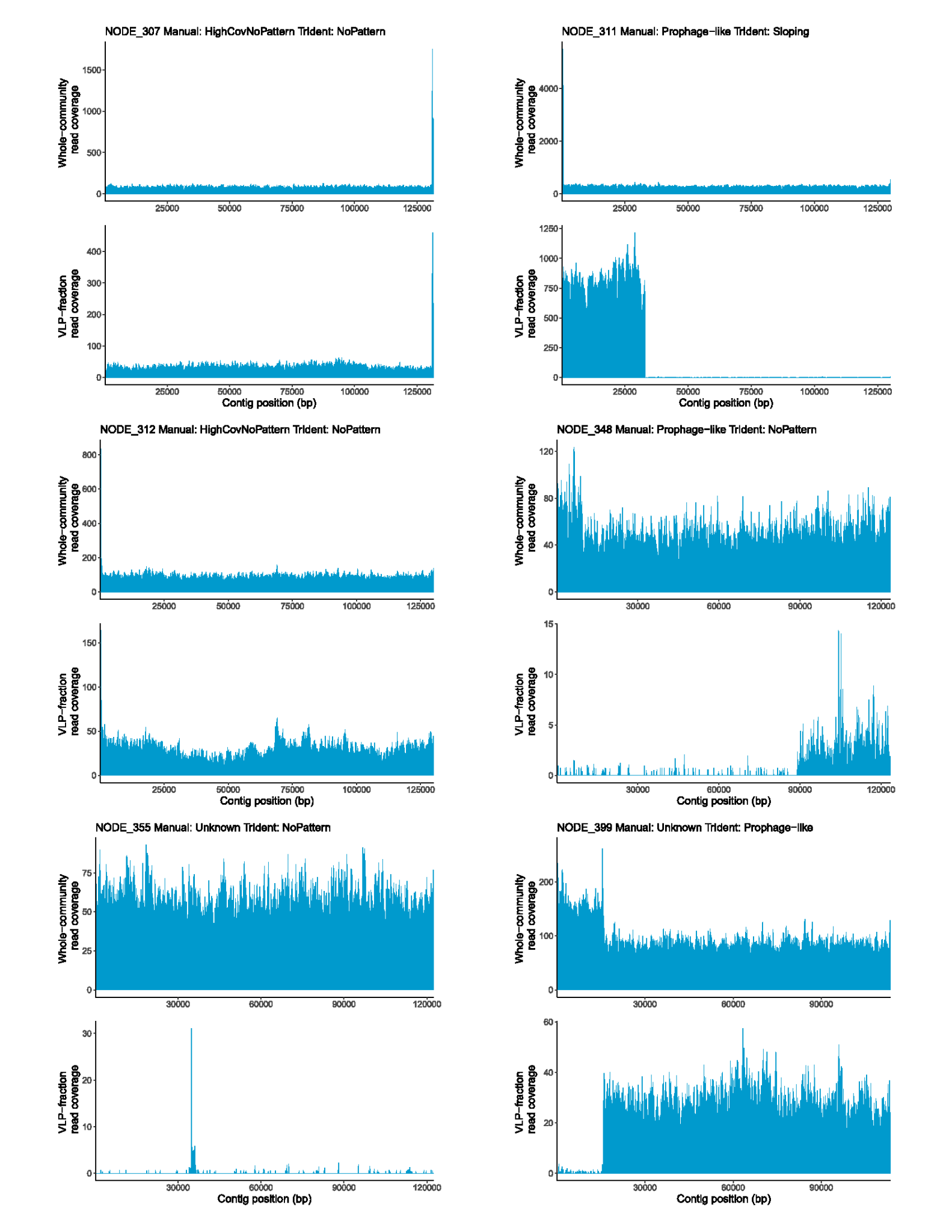

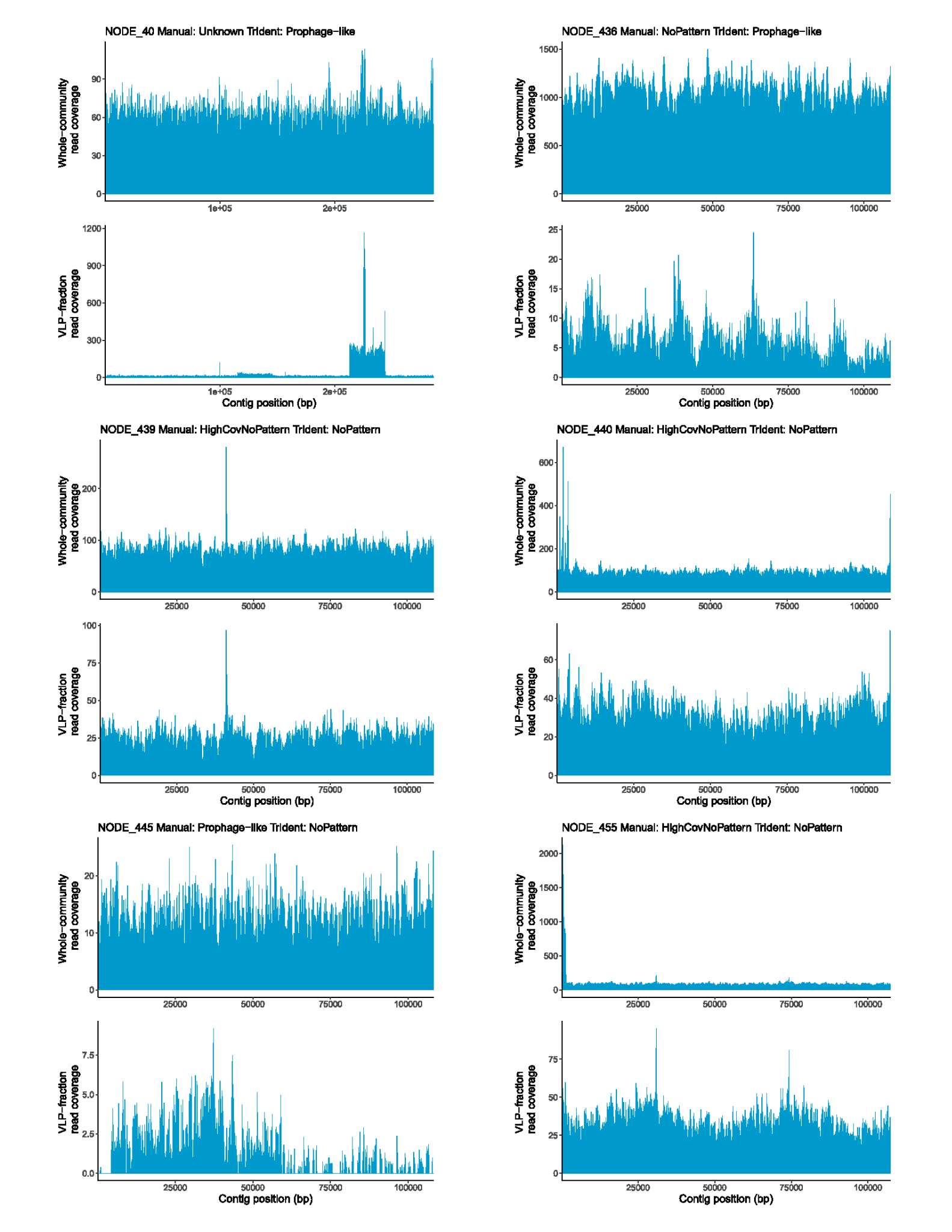

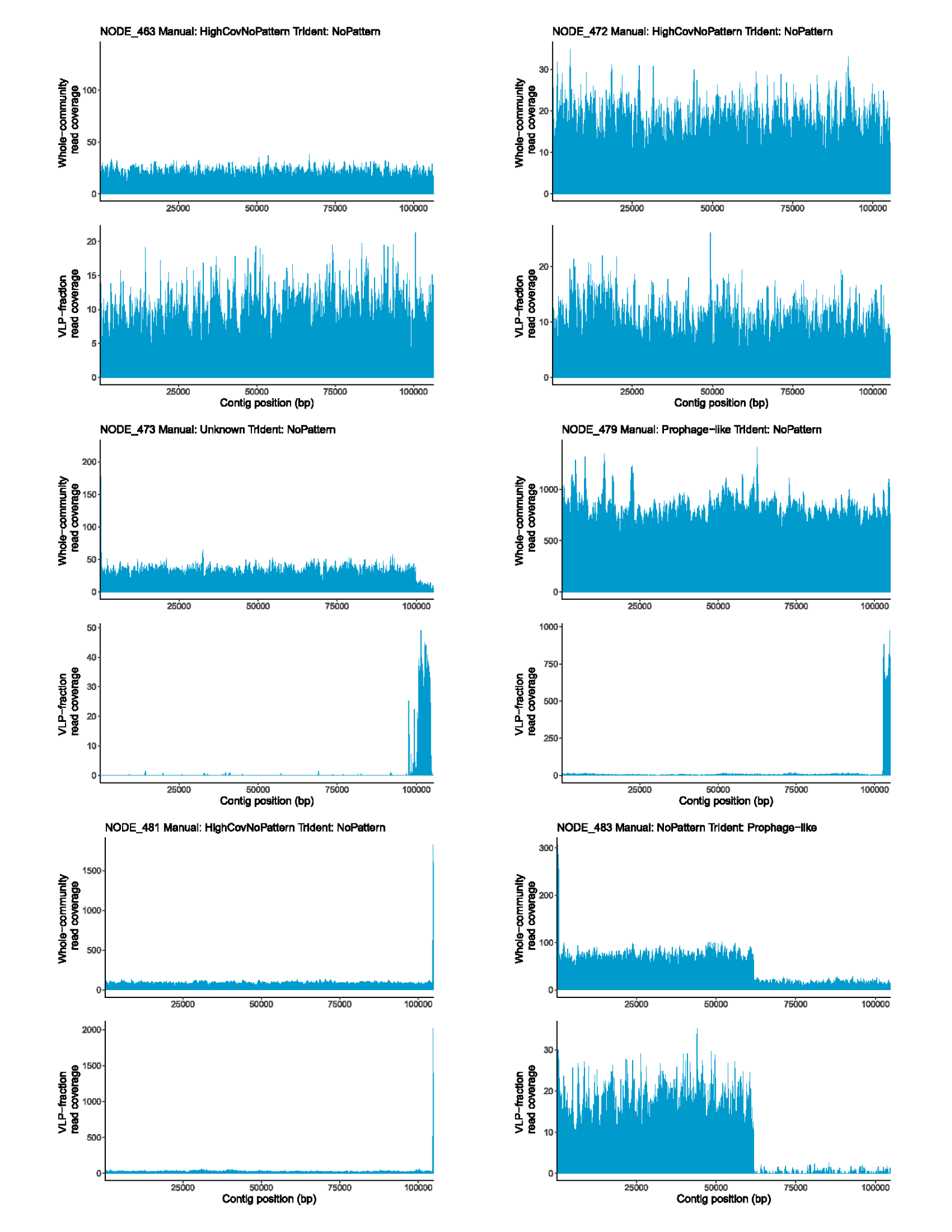

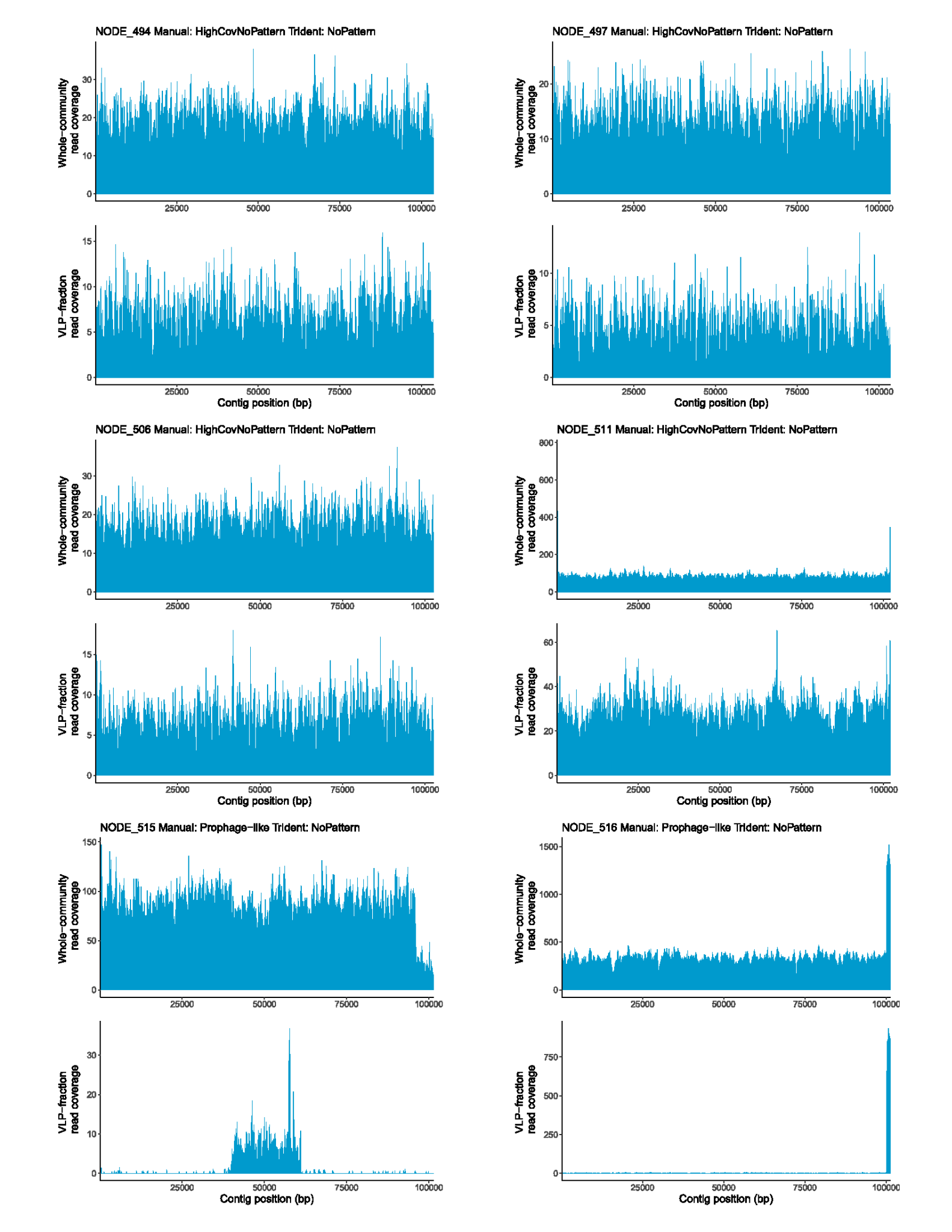

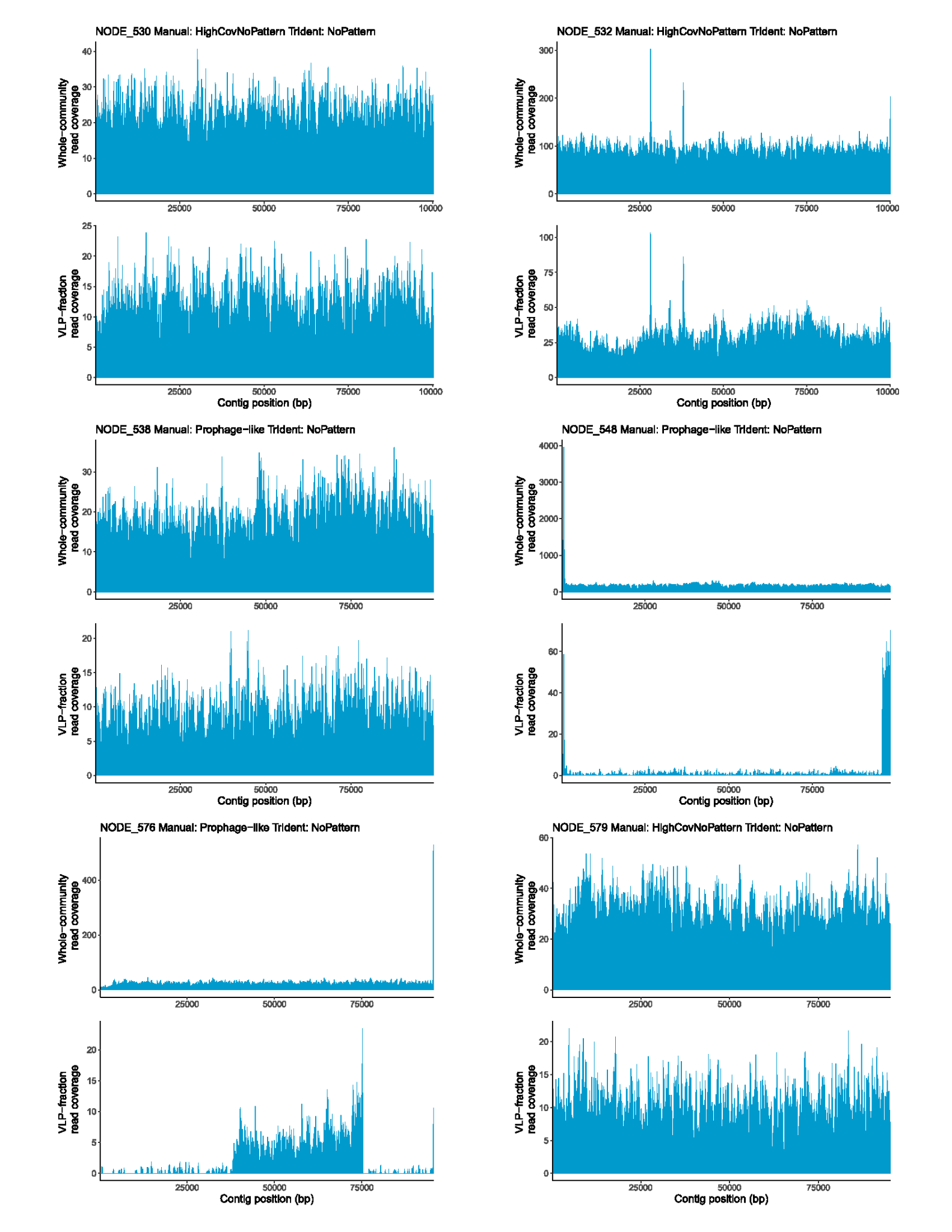

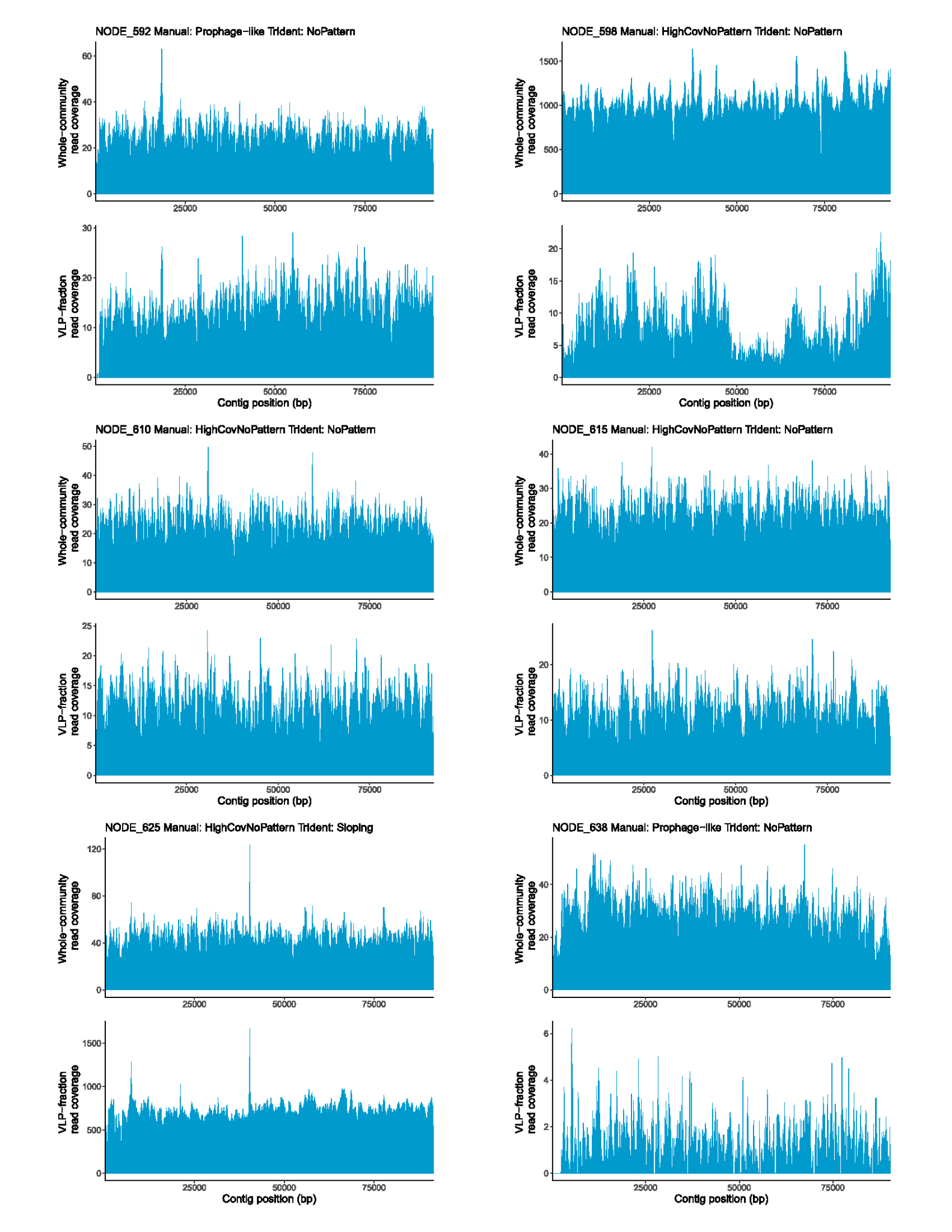

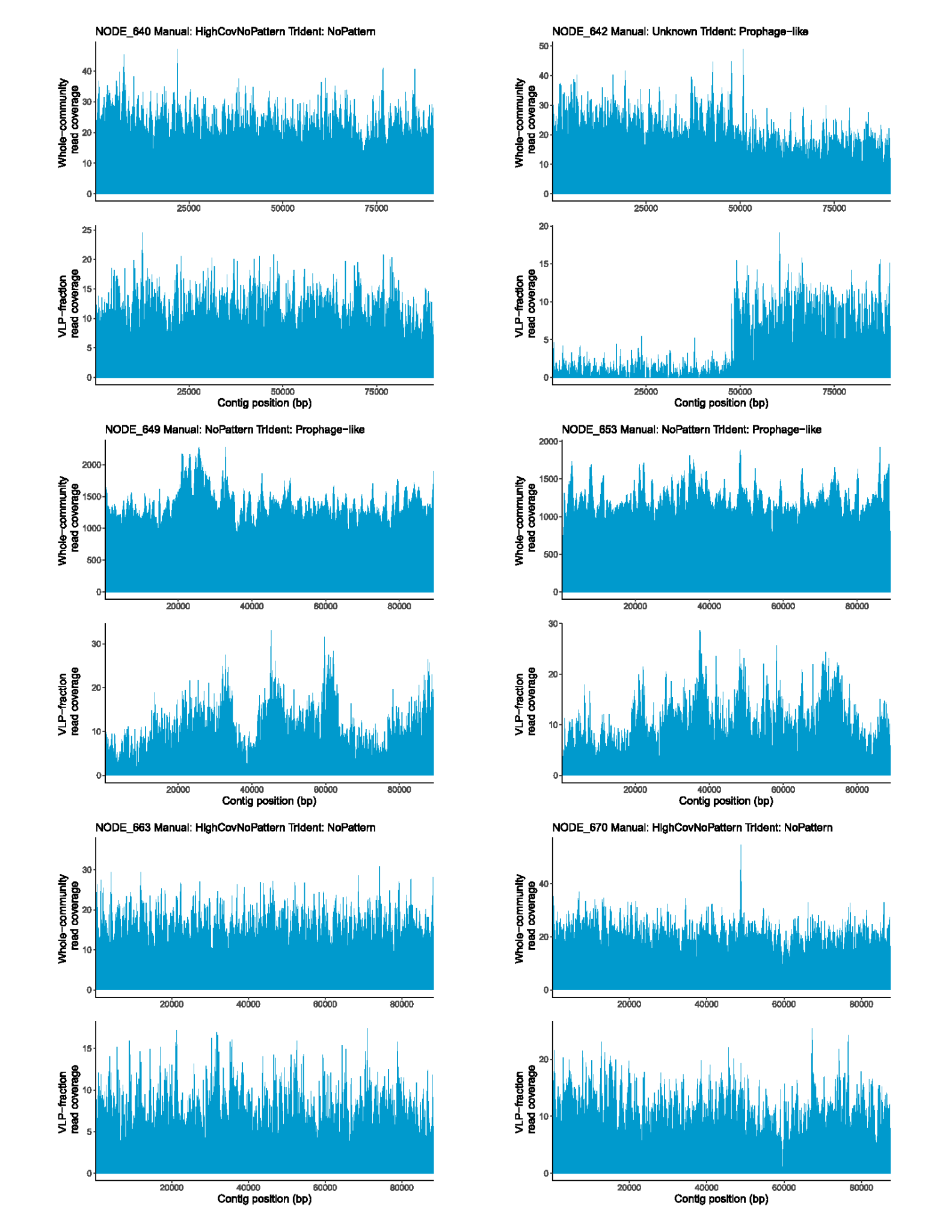

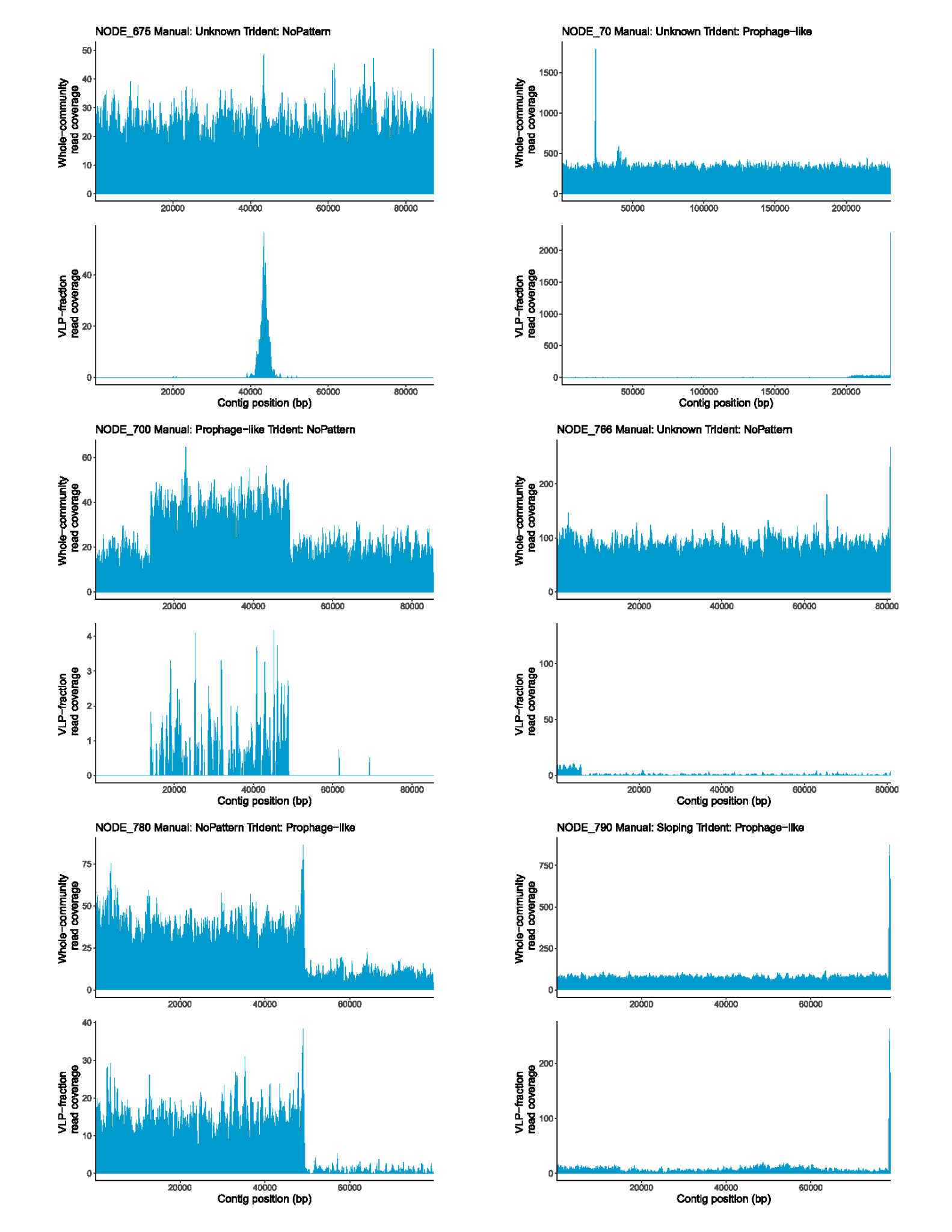

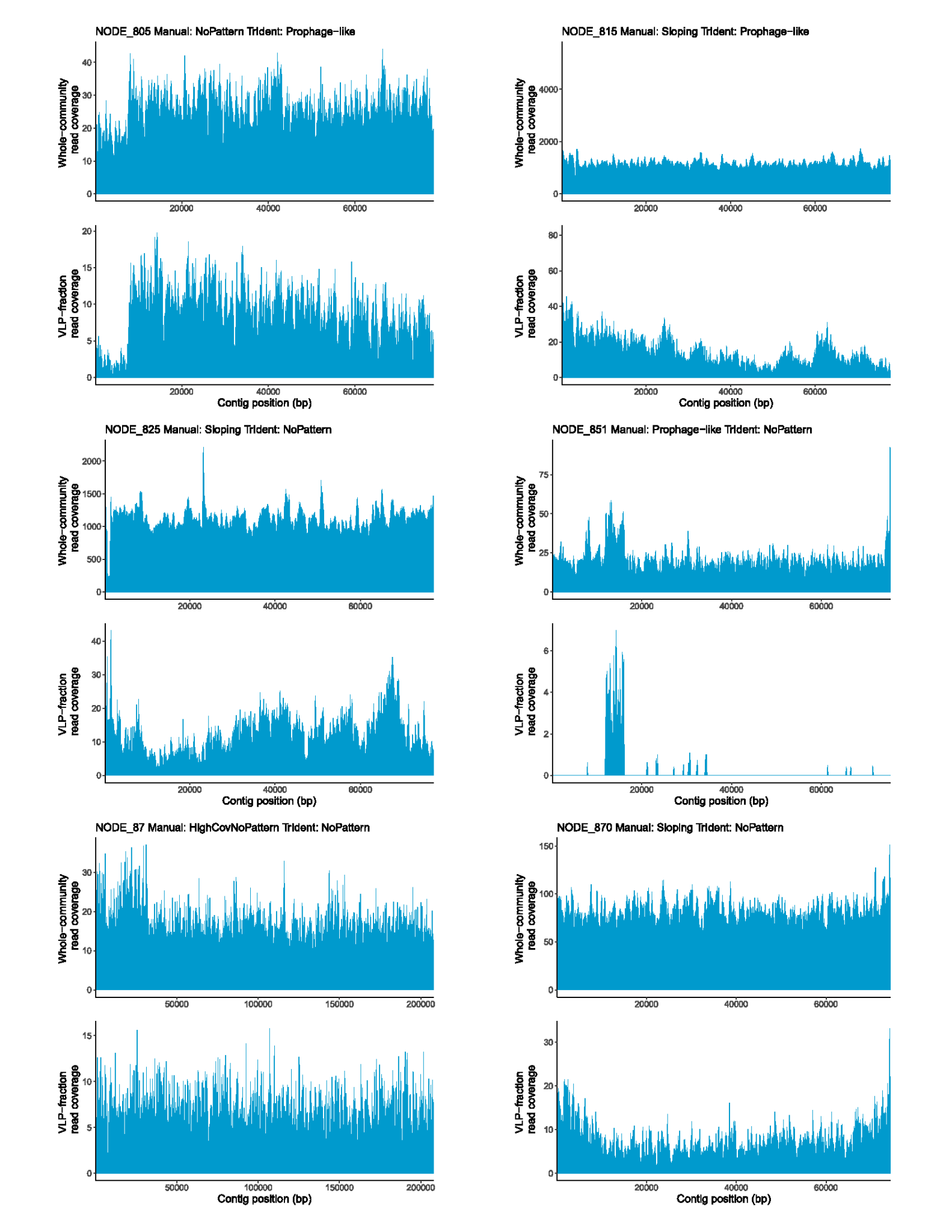

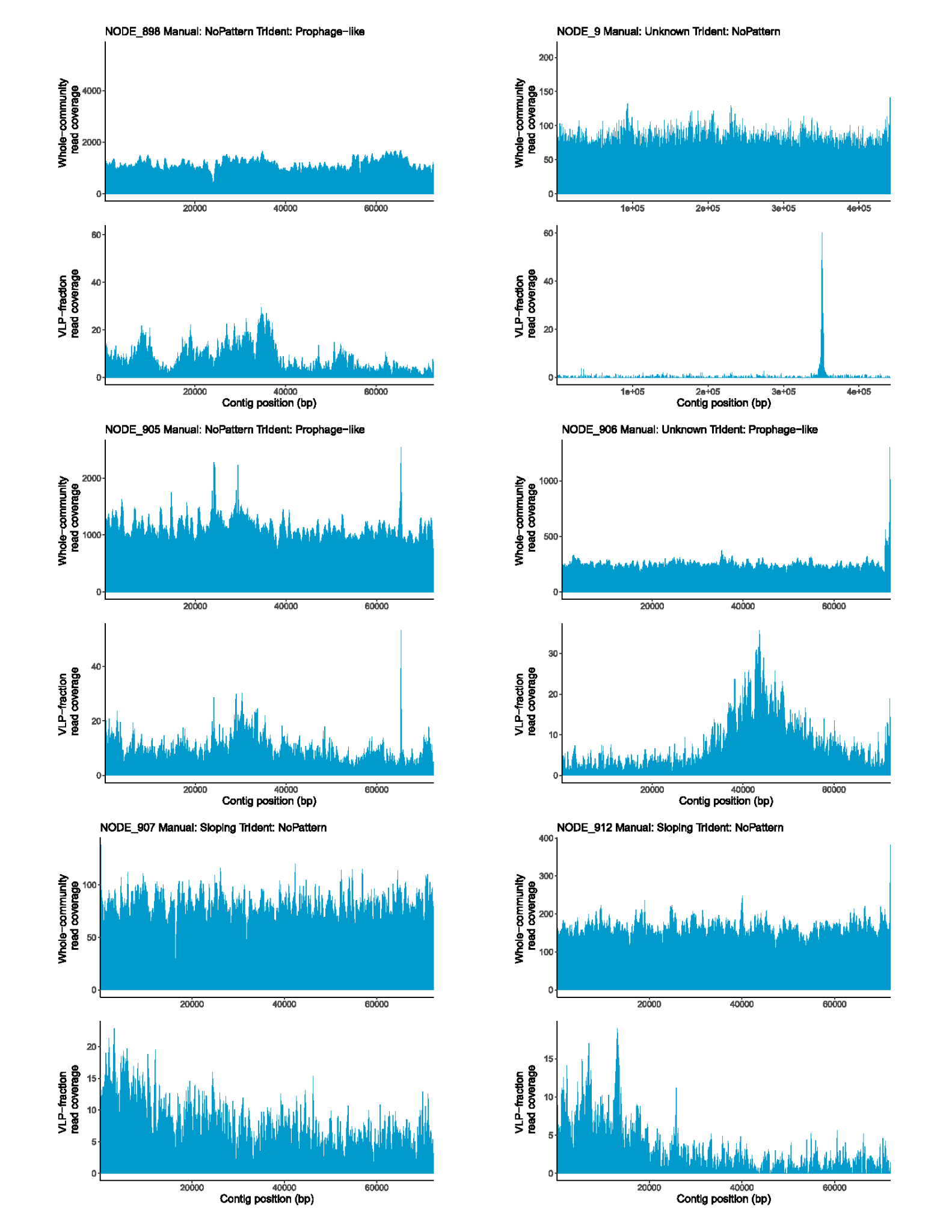

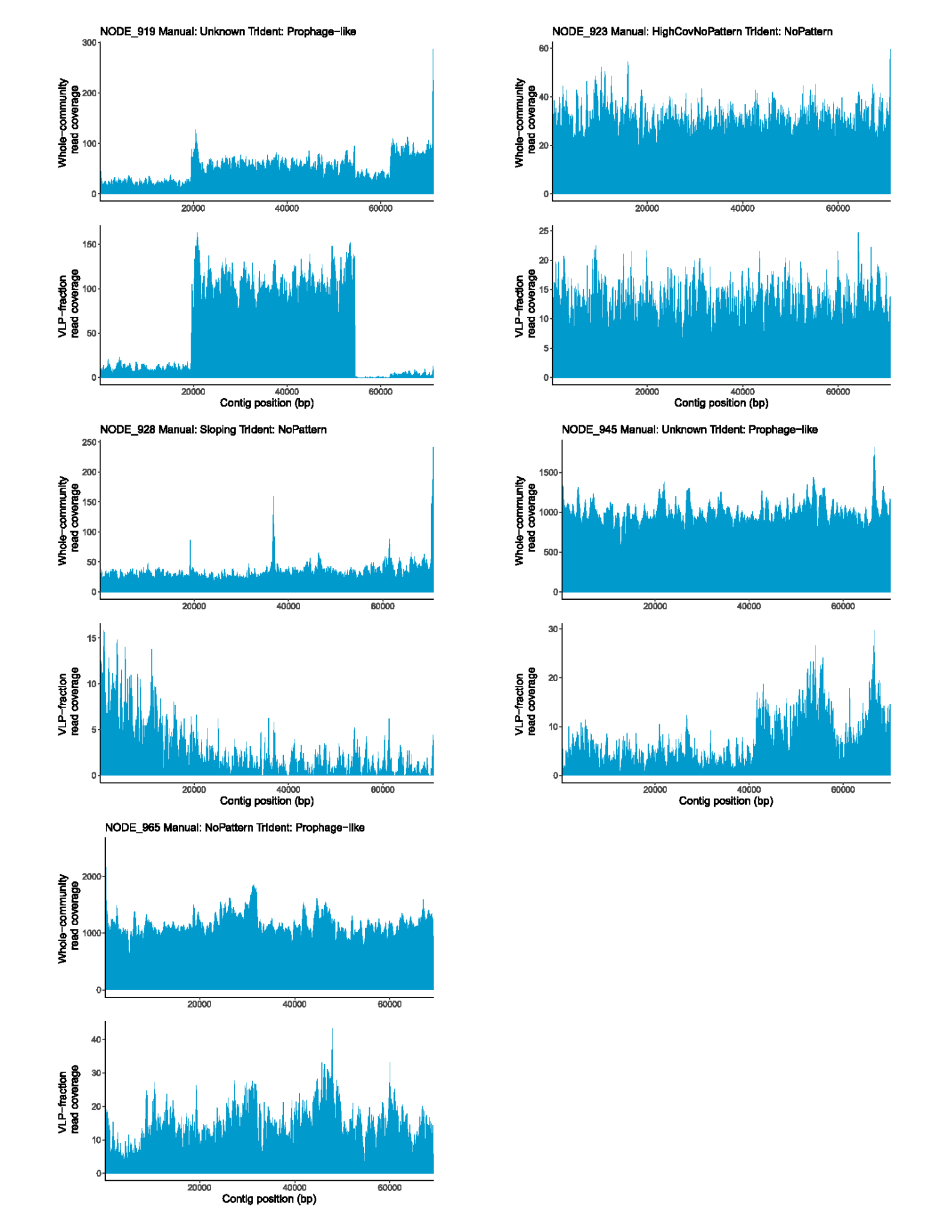

**Supplementary figure 3- Read coverage patterns of contigs with mismatched classifications in previously generated transductomics data.**

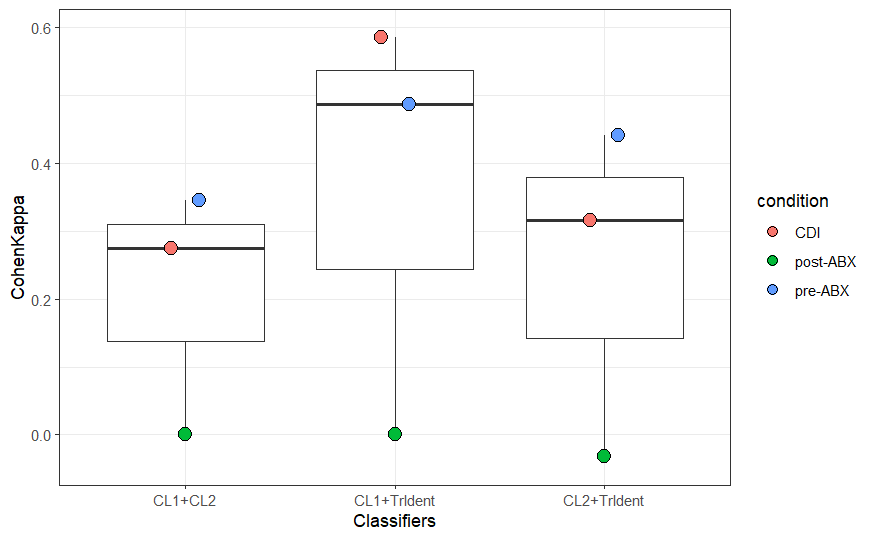

**Supplementary figure 4 - Cohen’s Kappa scores generated between each classifier for Pre-ABX, Post-ABX and CDI transductomics datasets.** Cohen’s Kappa is a measure of interrater reliability that takes into account agreement based on random chance. There are no significant differences (p < 0.05) between Cohen’s kappa scores between classifiers (Wilcoxon test).

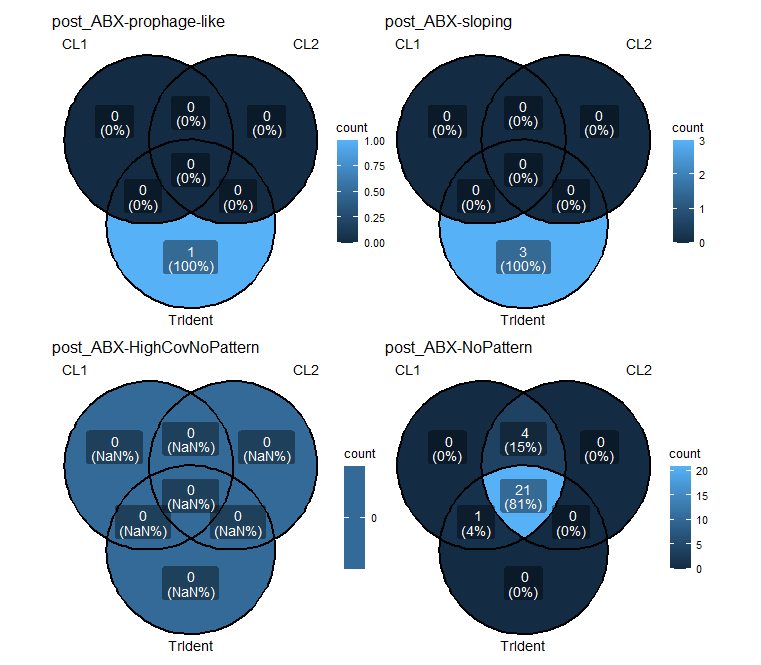

**Supplementary figure 5 - Classification Venn diagrams for ABX datasets.** Comparison of classifications made by the three classifiers (CL1, CL2, and TrIdent) in the ABX dataset for each of the four TrIdent pattern-classes - NoPattern, Prophage-like, Sloping, and HighCovNoPattern.

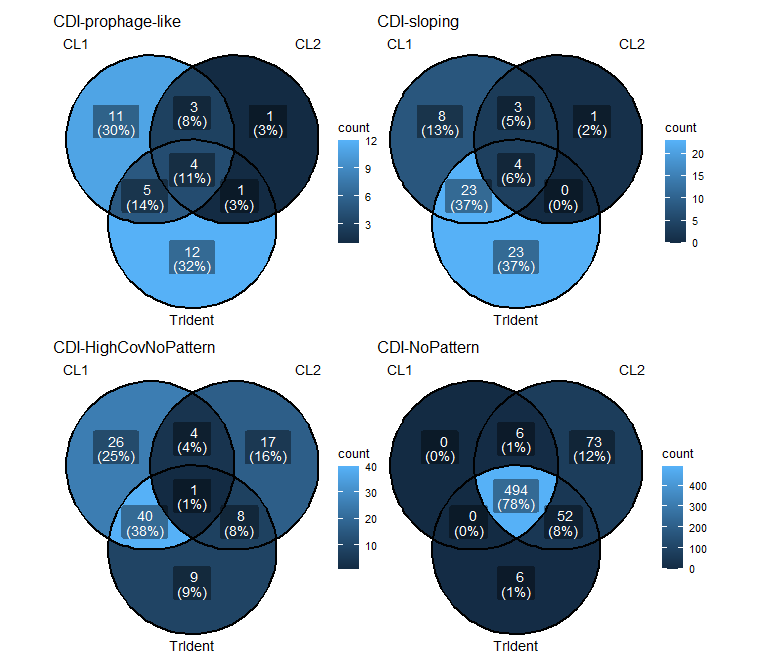

**Supplementary figure 6 - Classification Venn diagrams for *Clostridioides* difficle Infection (CDI) datasets.** Comparison of classifications made by the three classifiers (CL1, CL2, and TrIdent) in the CDI dataset for each of the four TrIdent pattern-classes - NoPattern, Prophage-like, Sloping, and HighCovNoPattern.

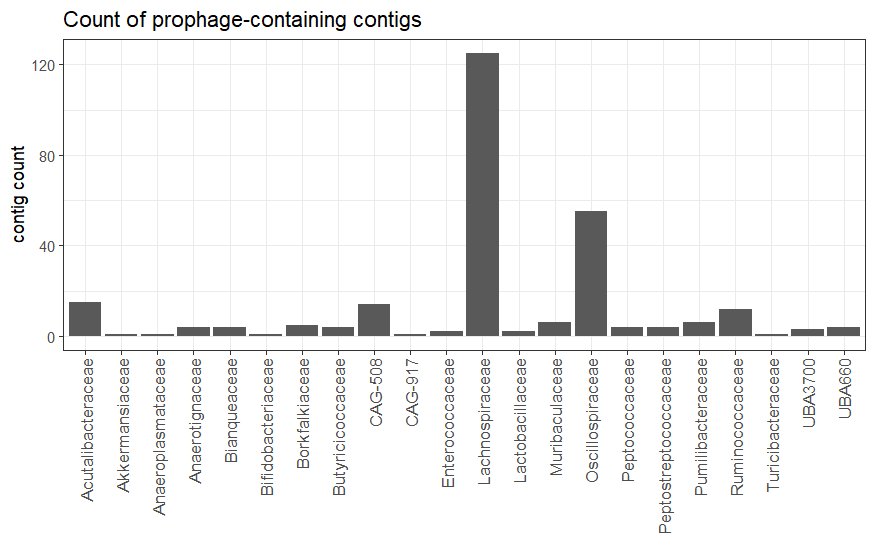

**Supplementary figure 7 - TrIdent classifications and familial taxonomy of contigs classified as ‘provirus’ by geNomad.** The count of contigs classified as ‘provirus’ by geNomad.

**Note- multipage figure, caption below**
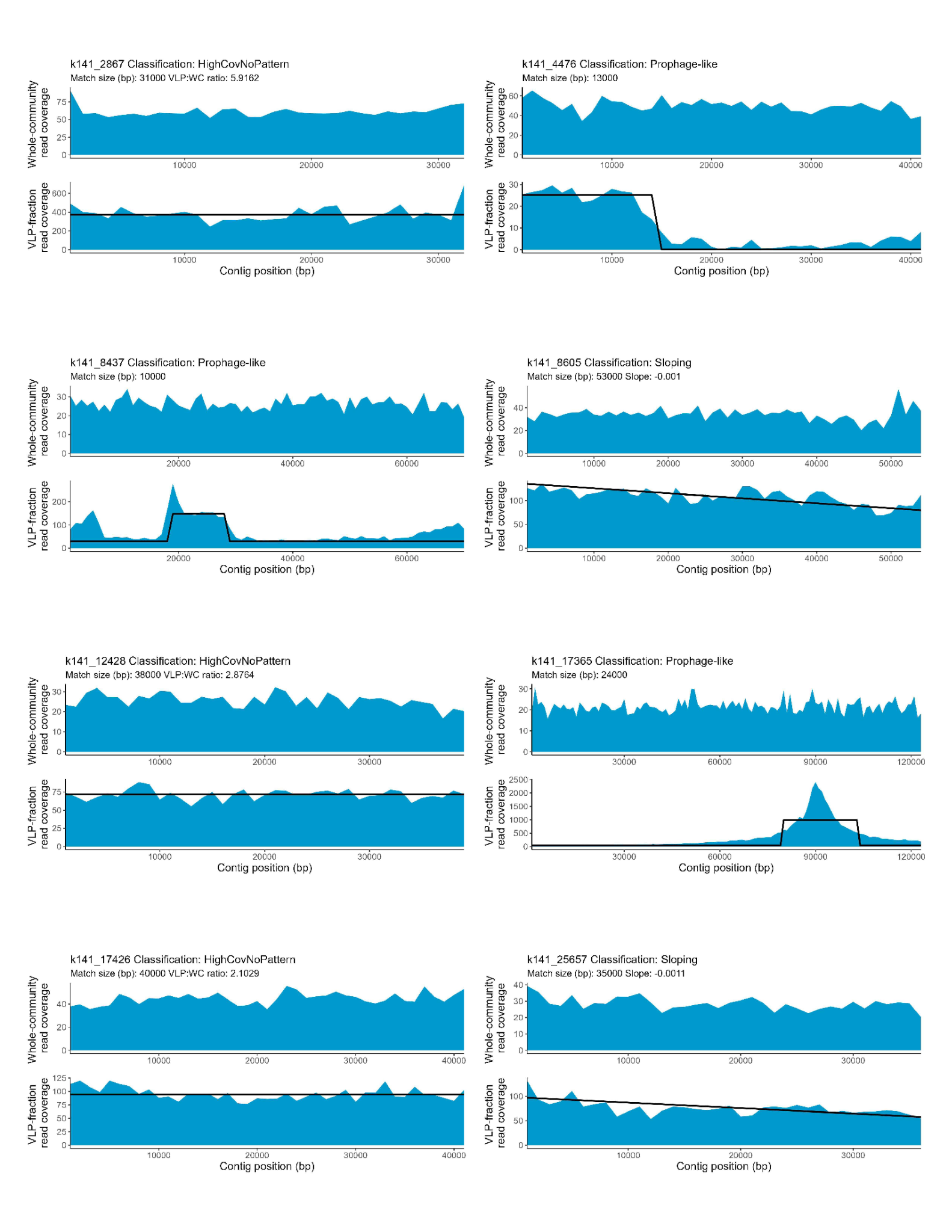

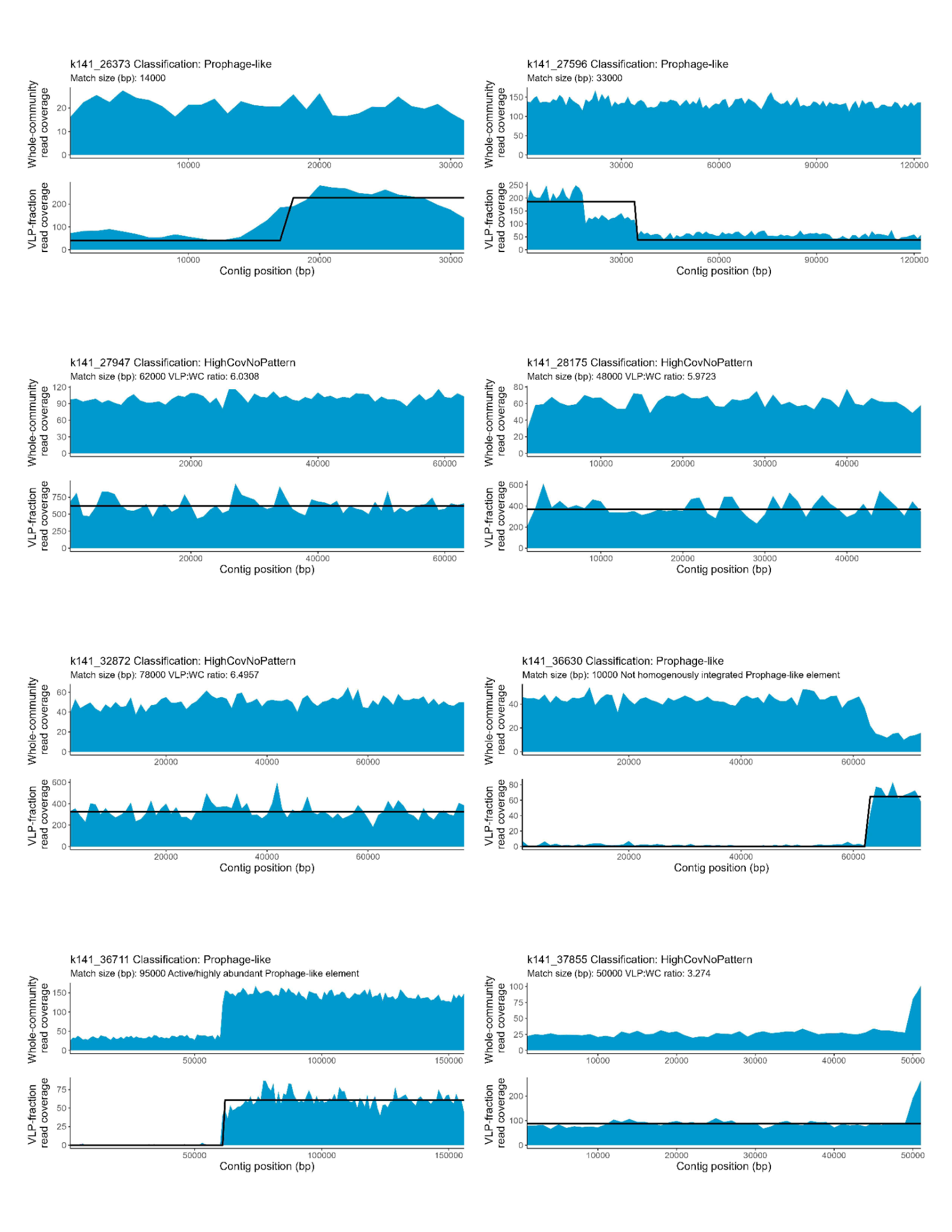

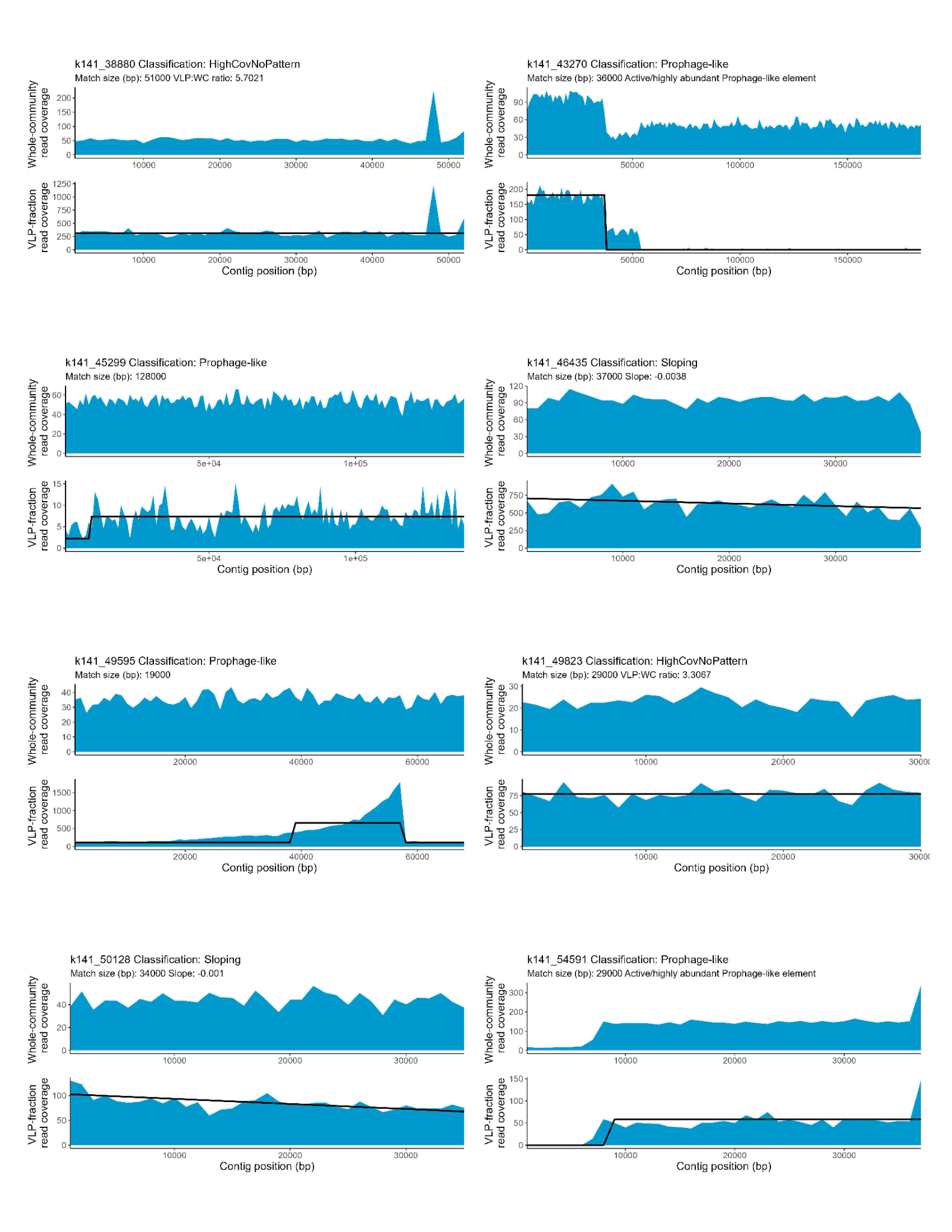

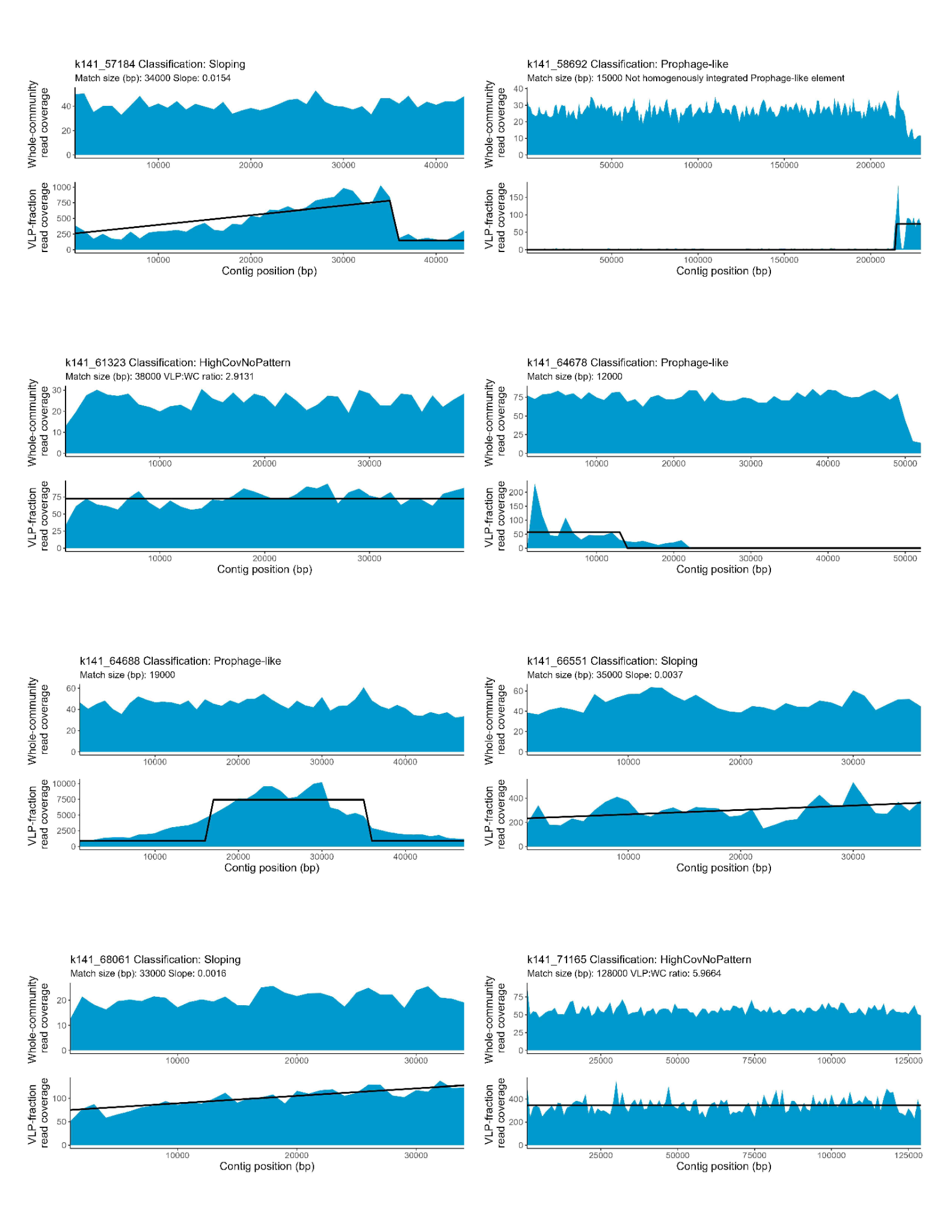

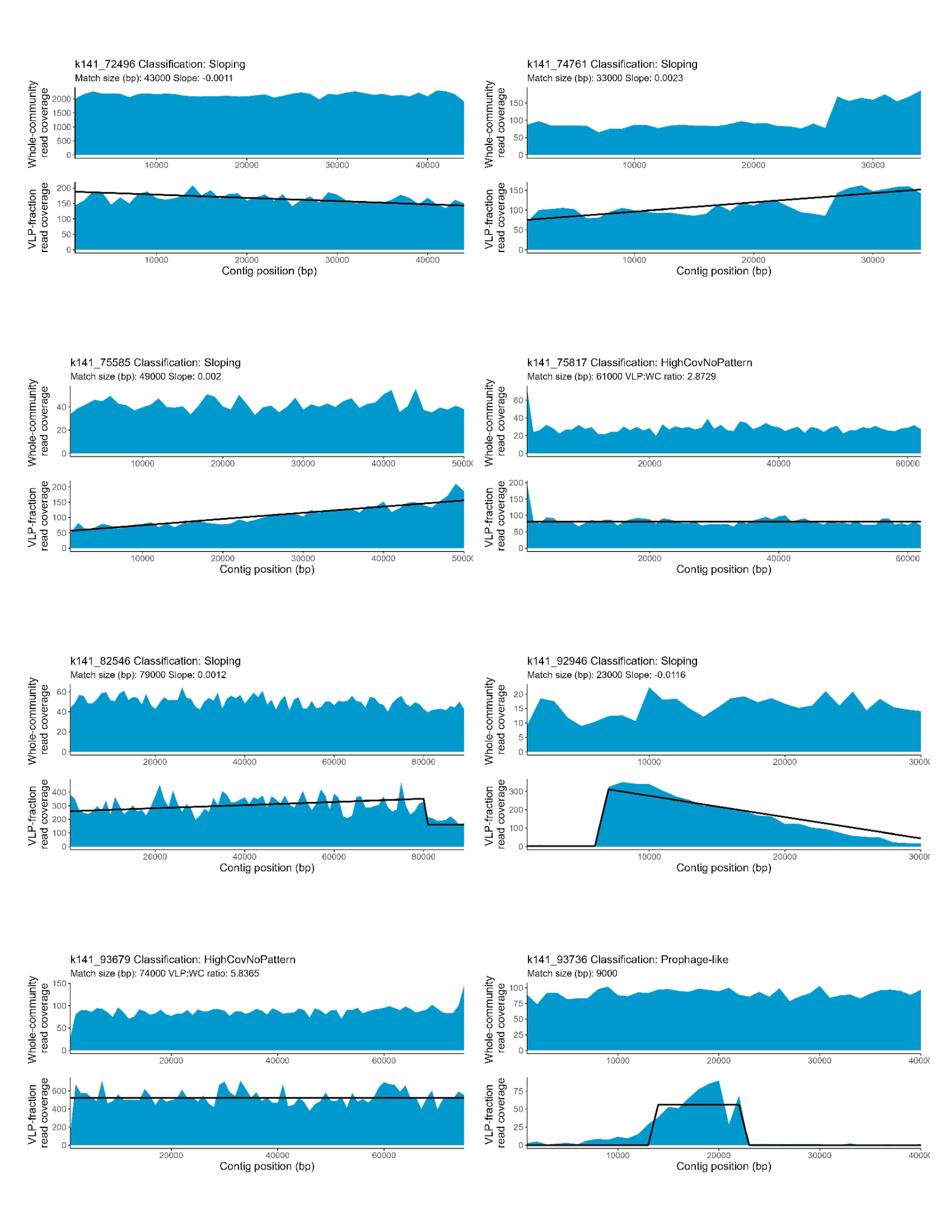

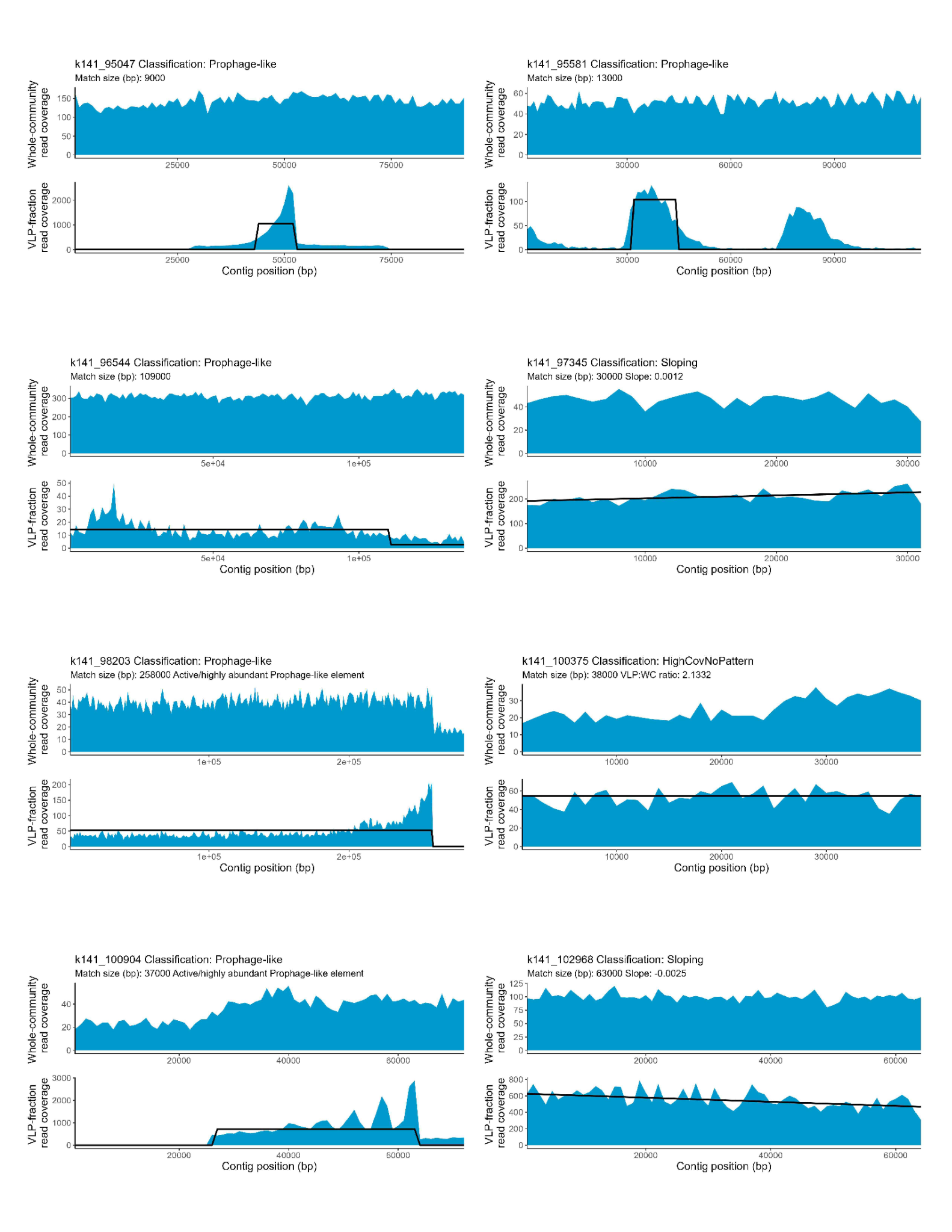

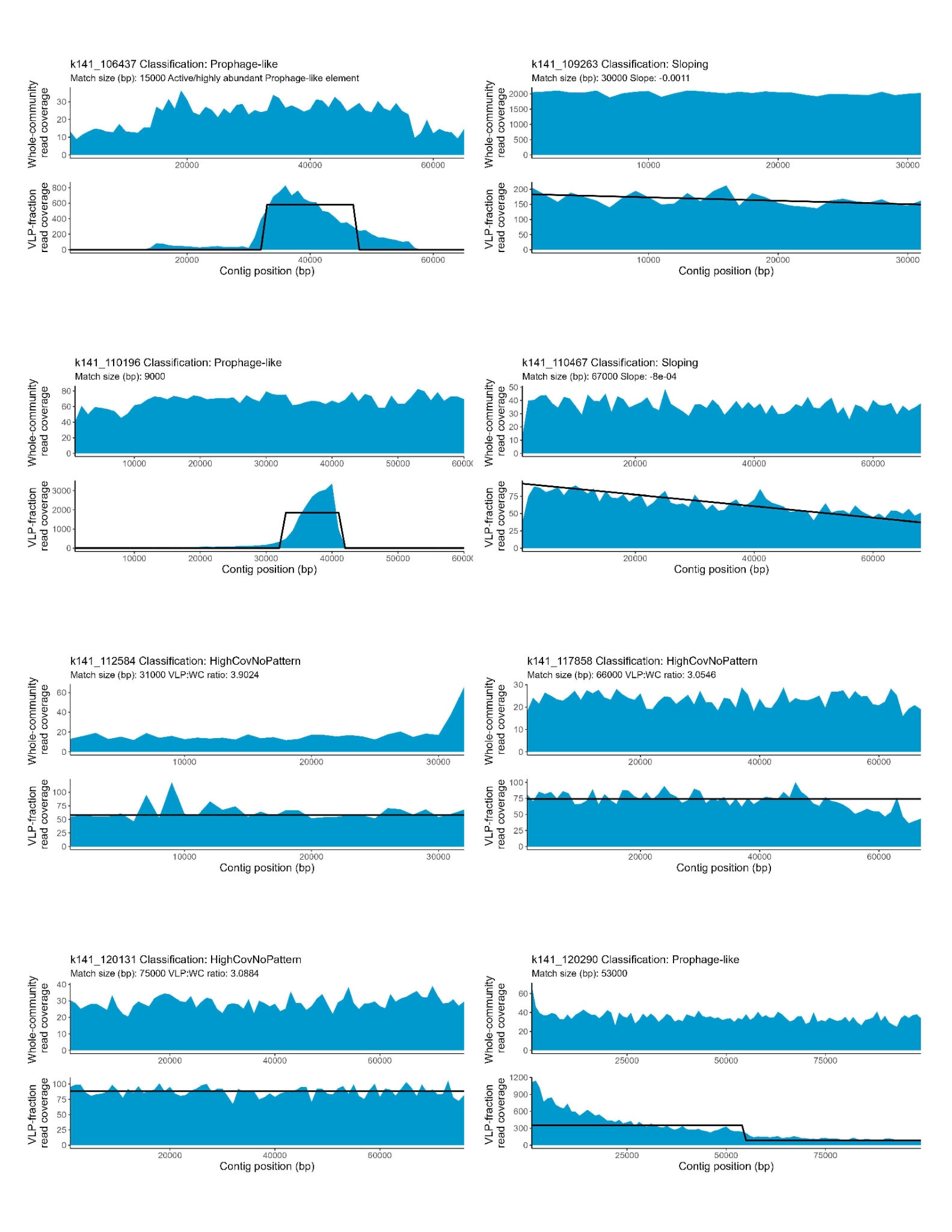

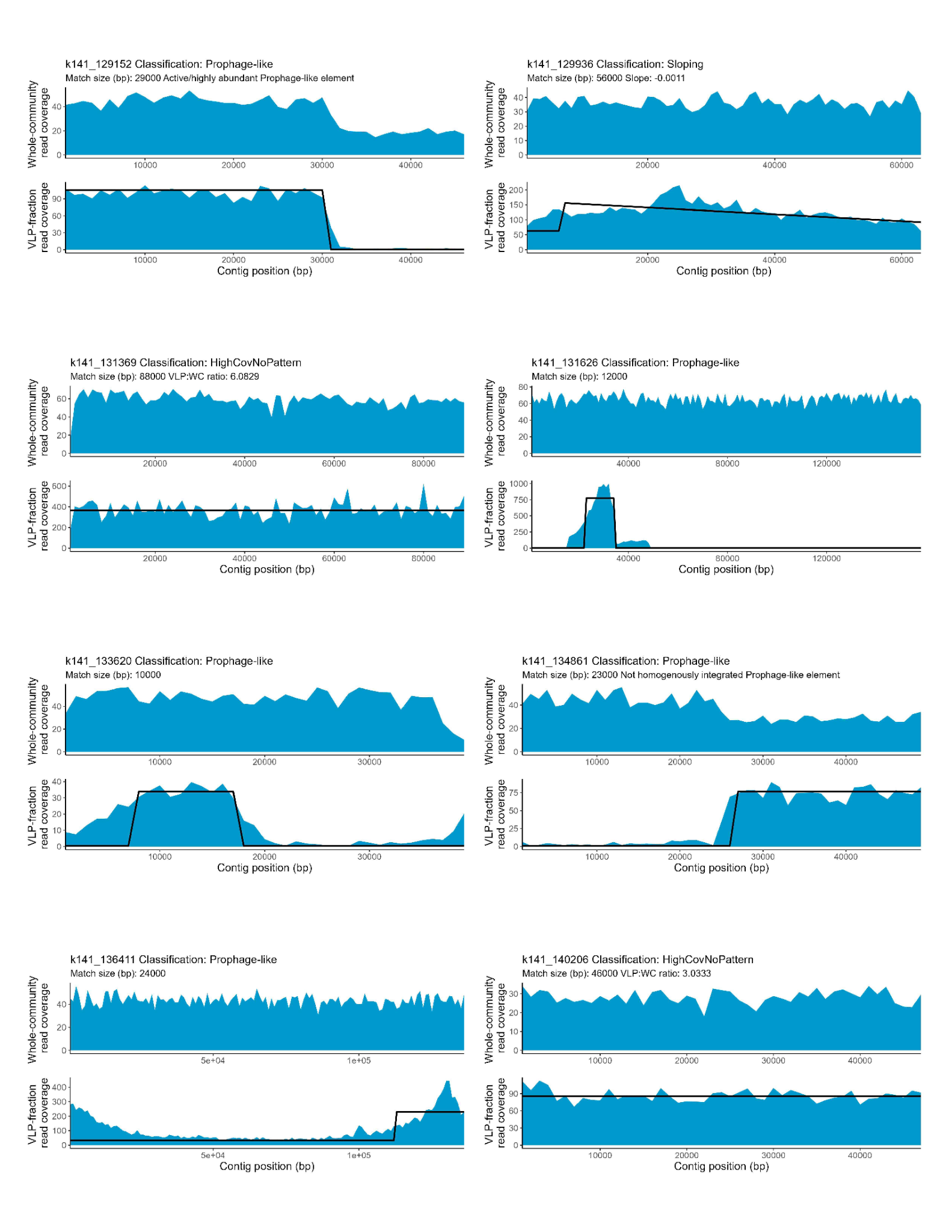

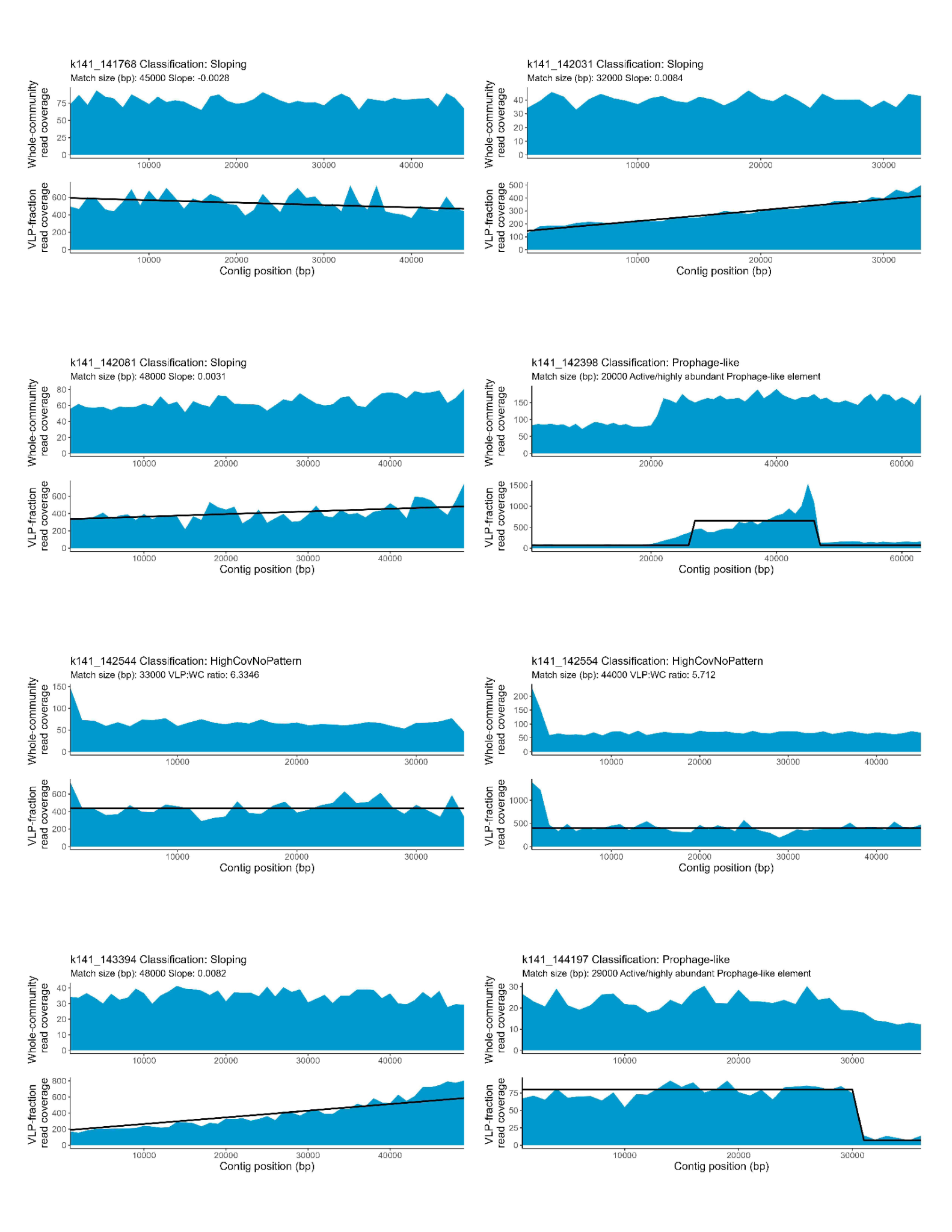

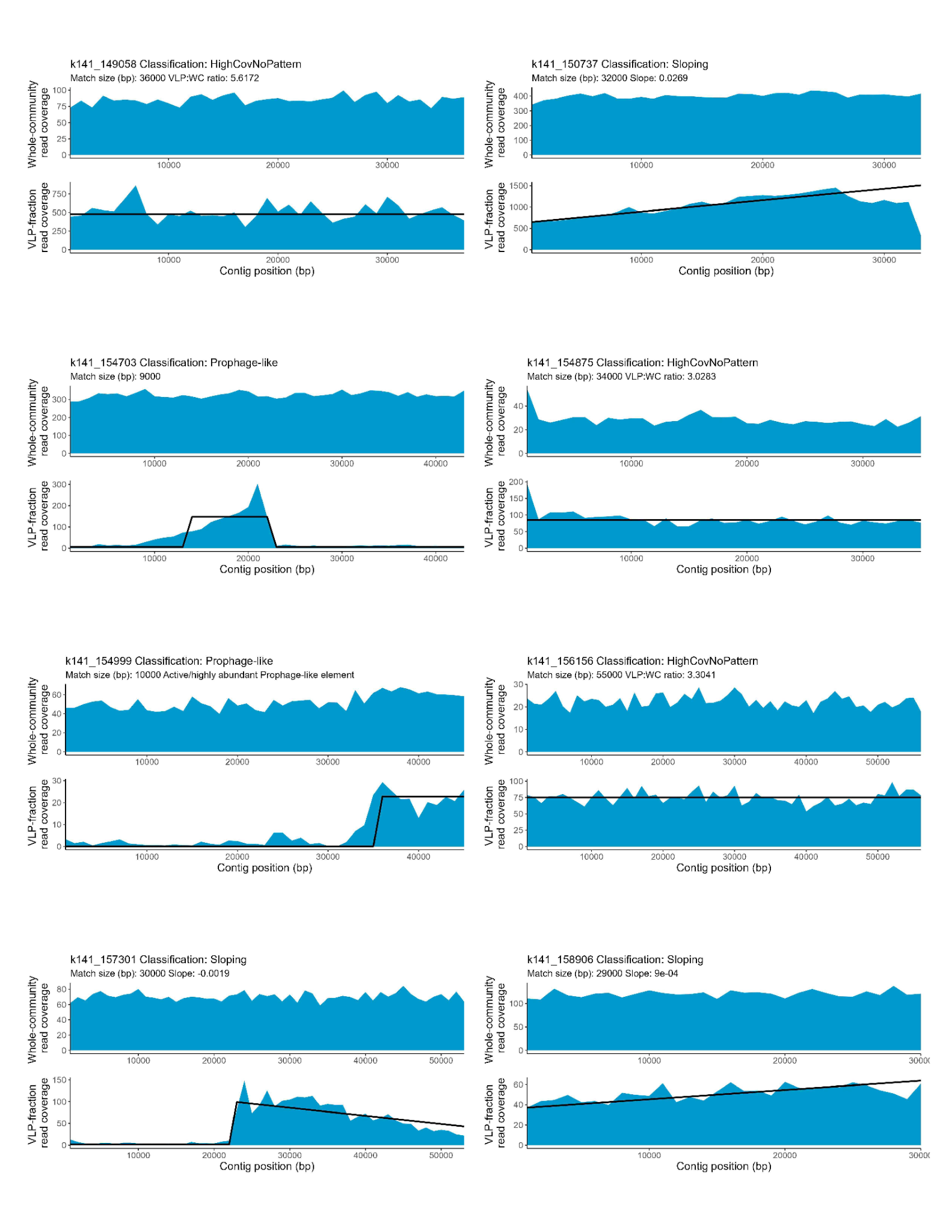

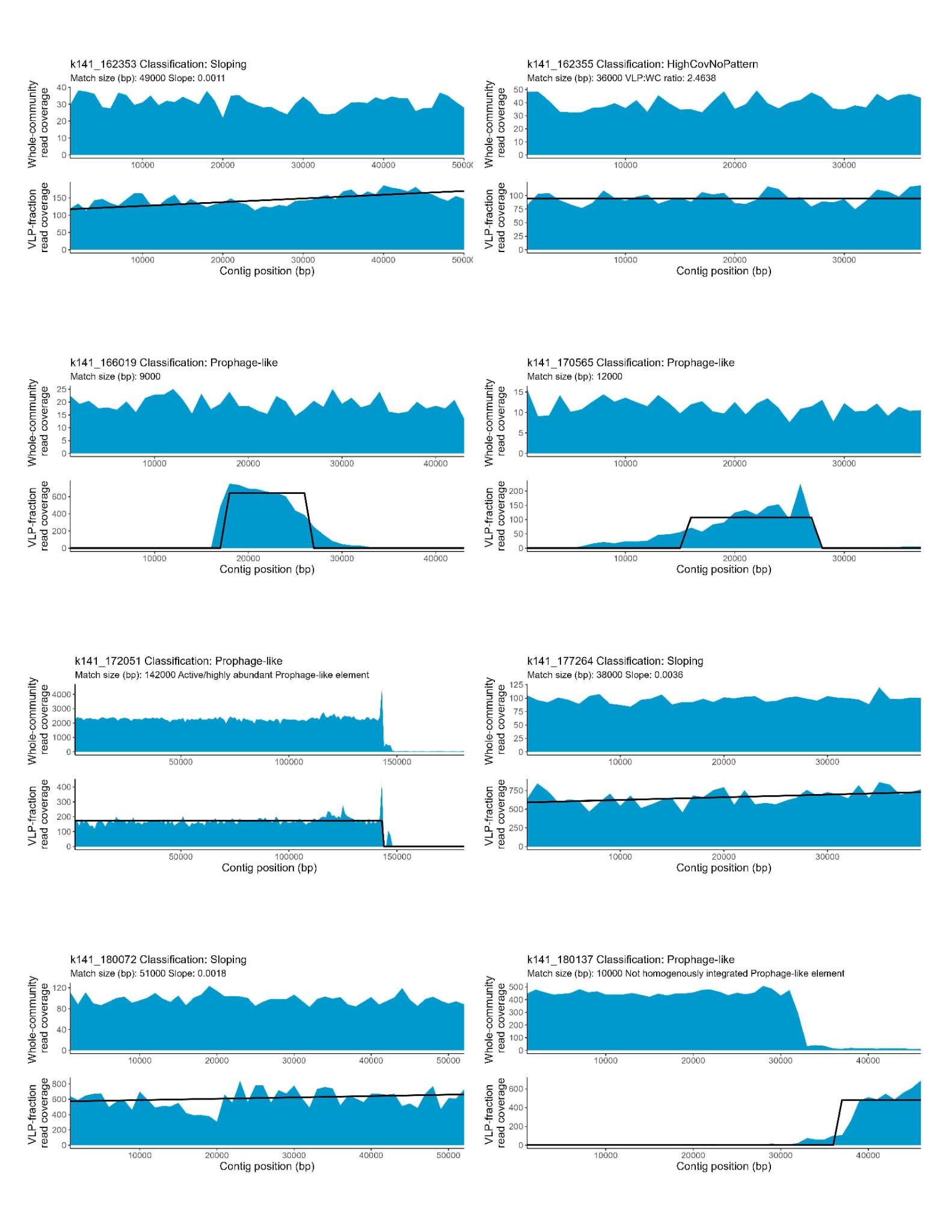

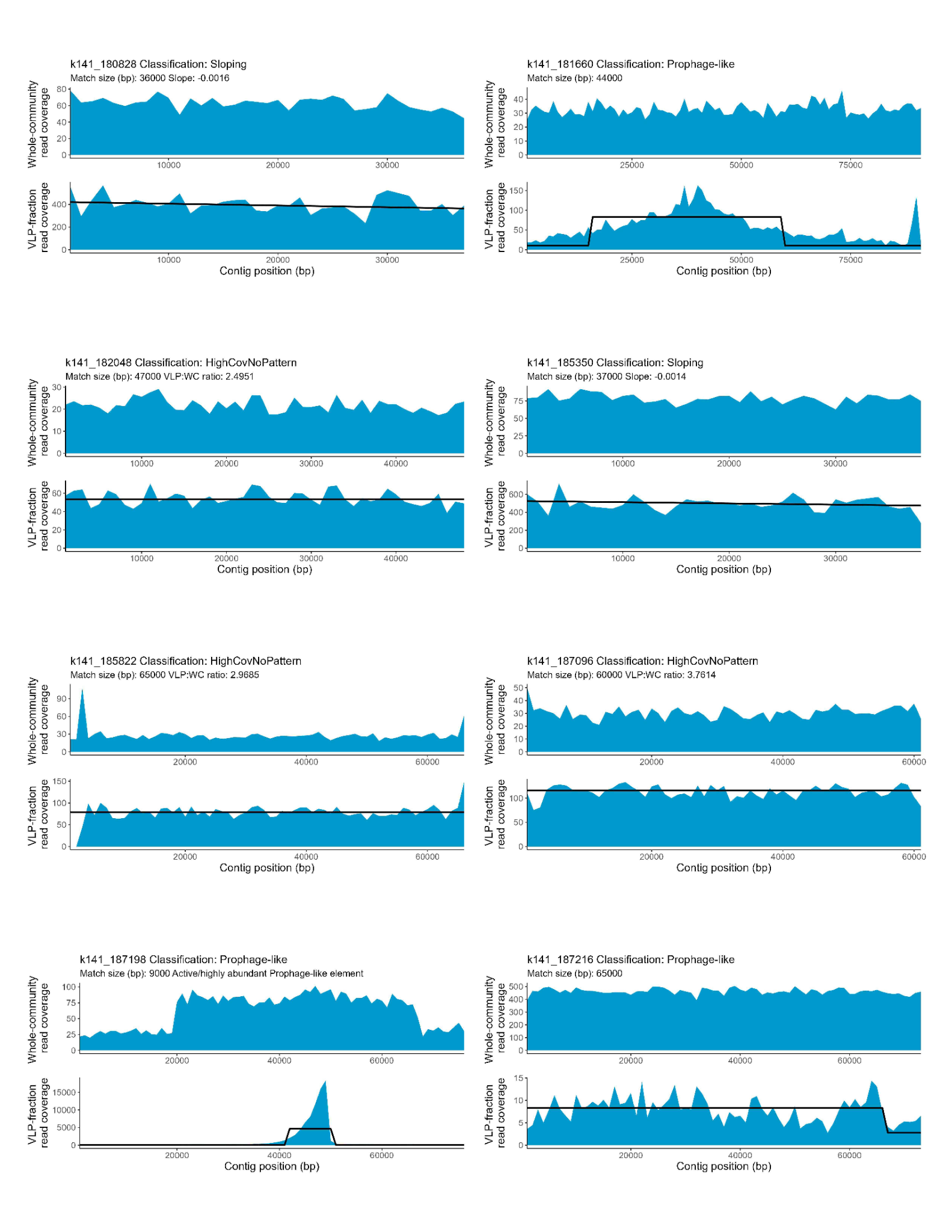

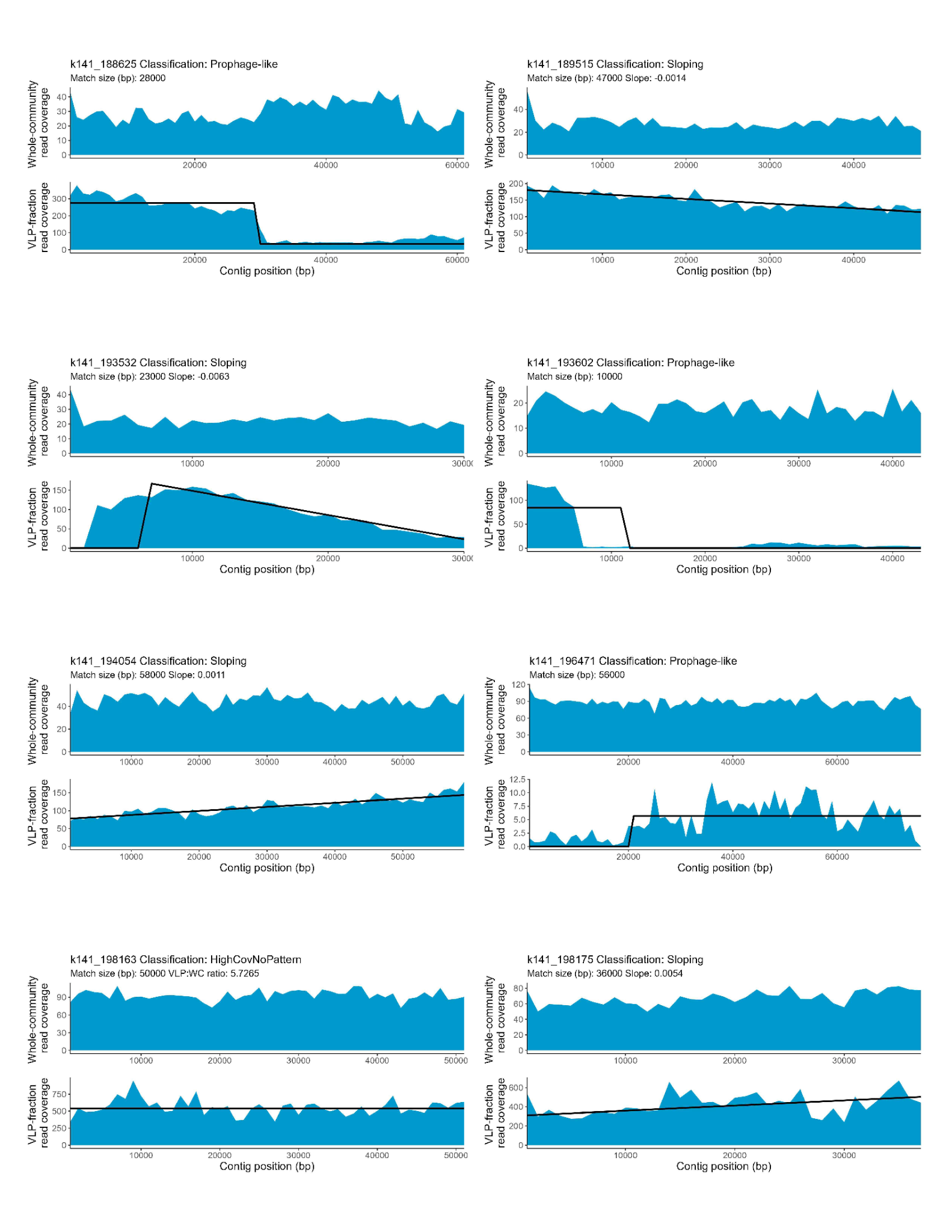

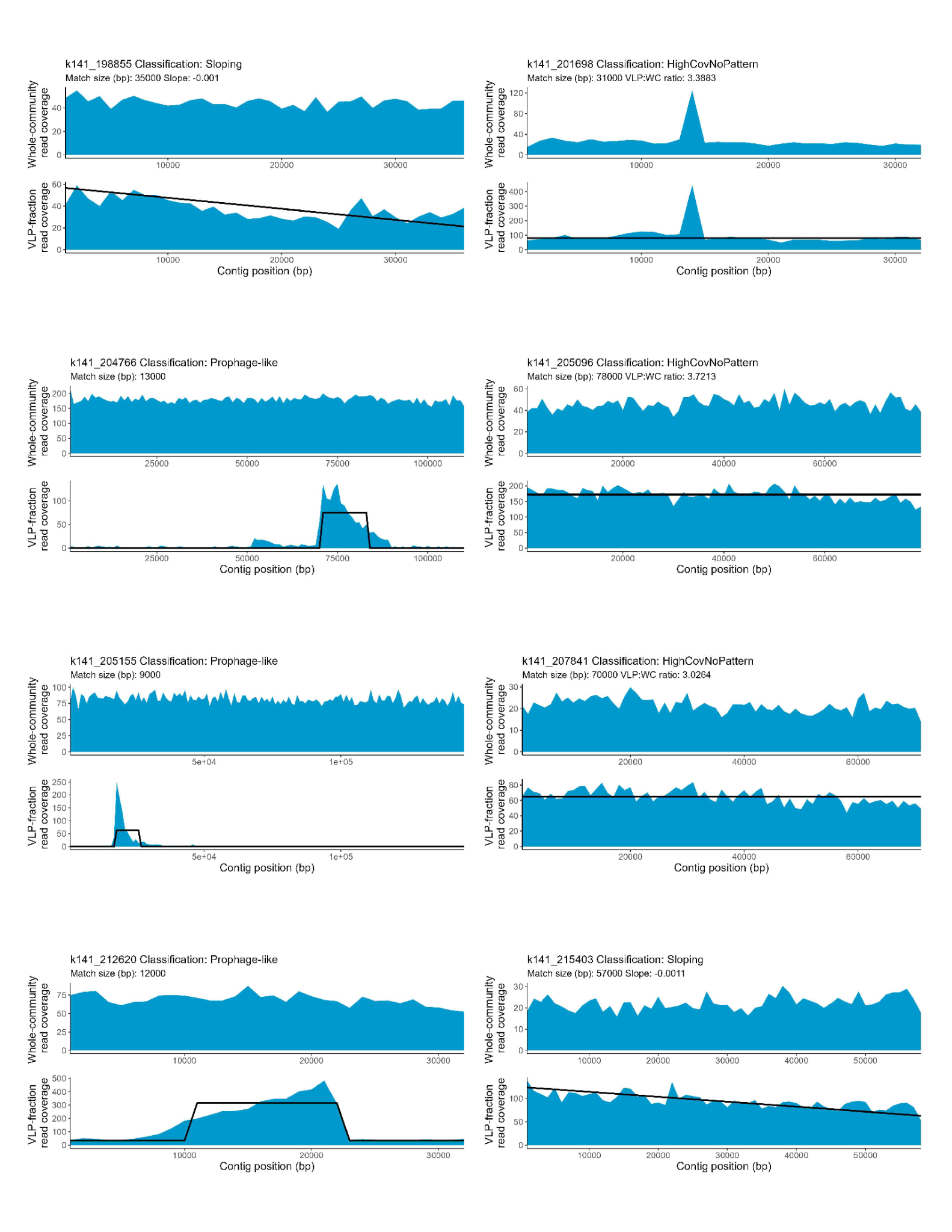

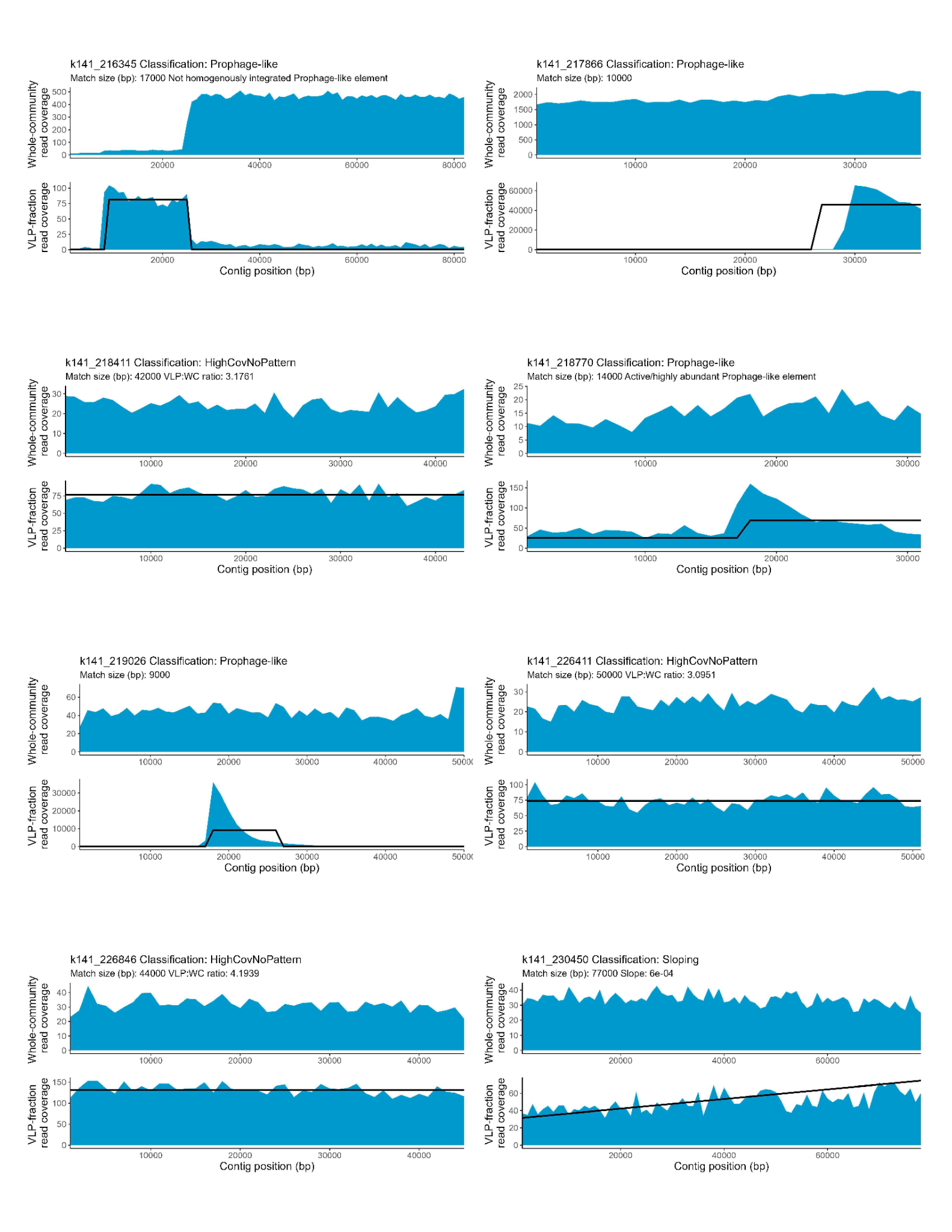

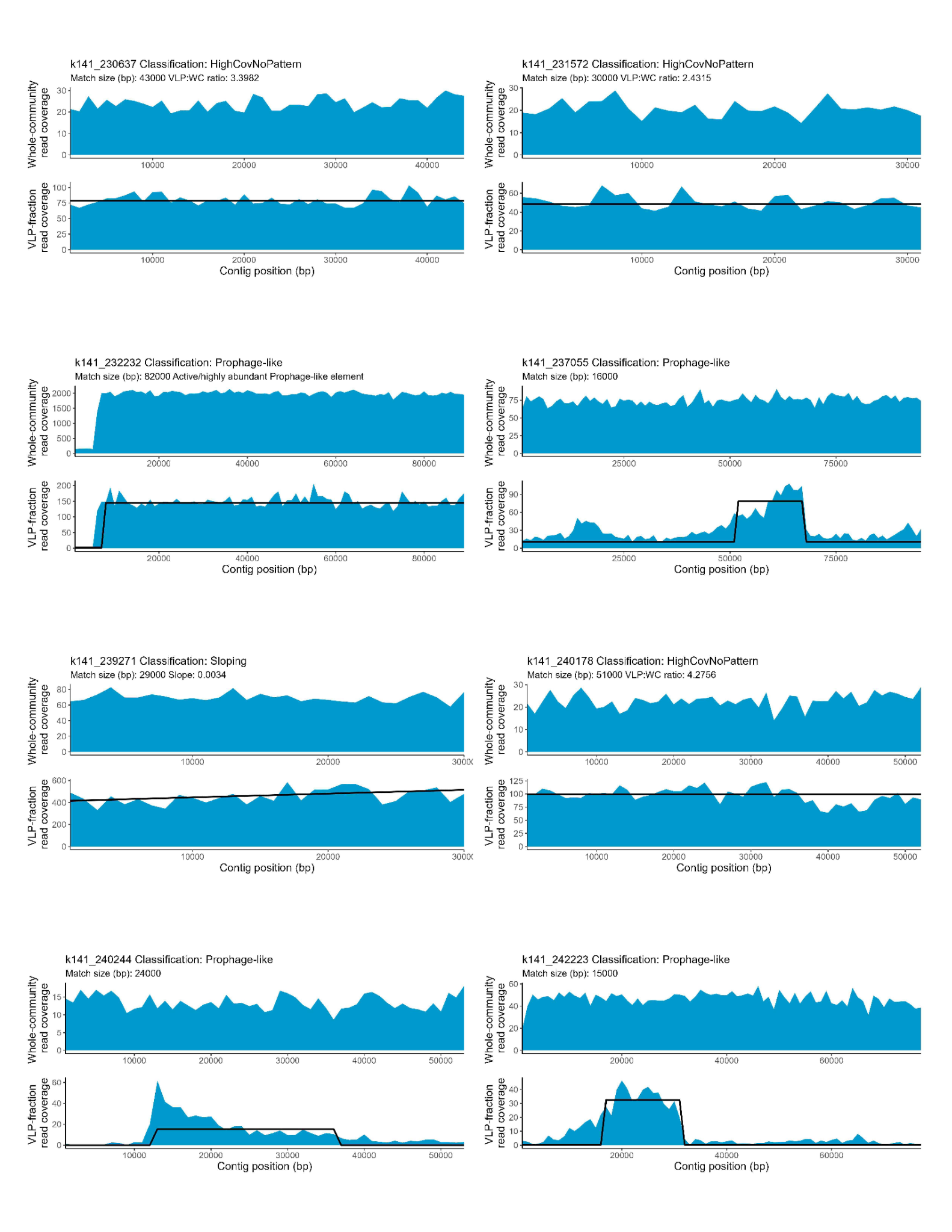

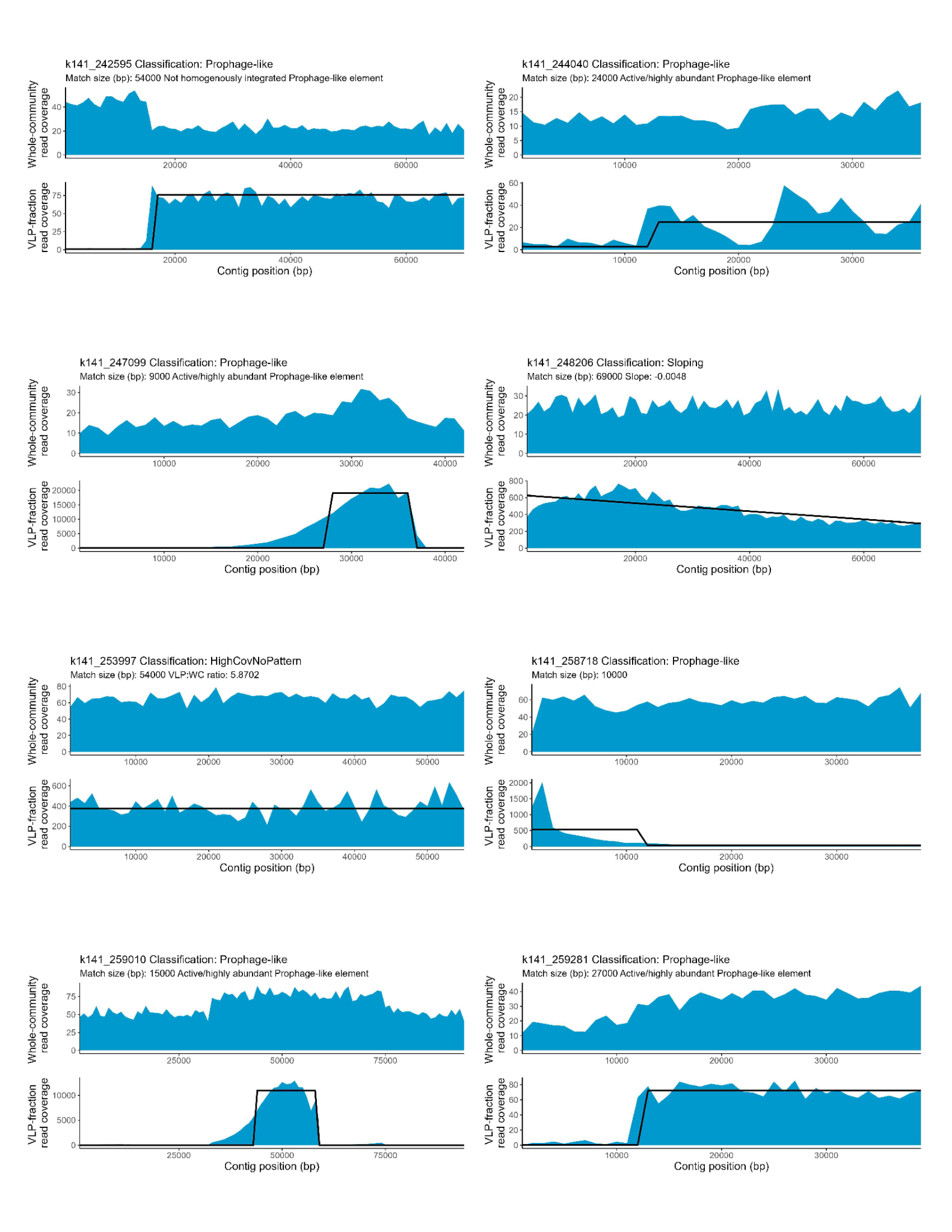

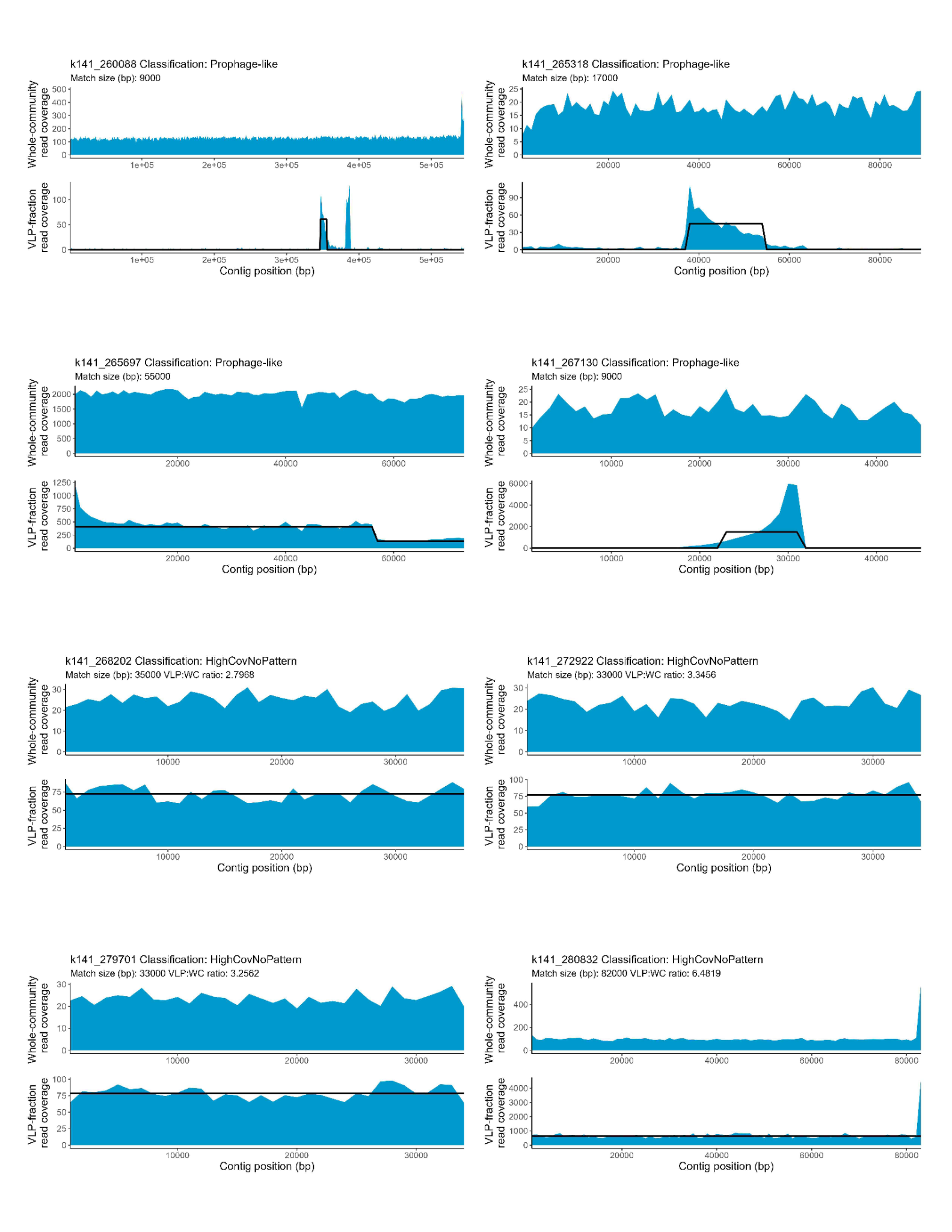

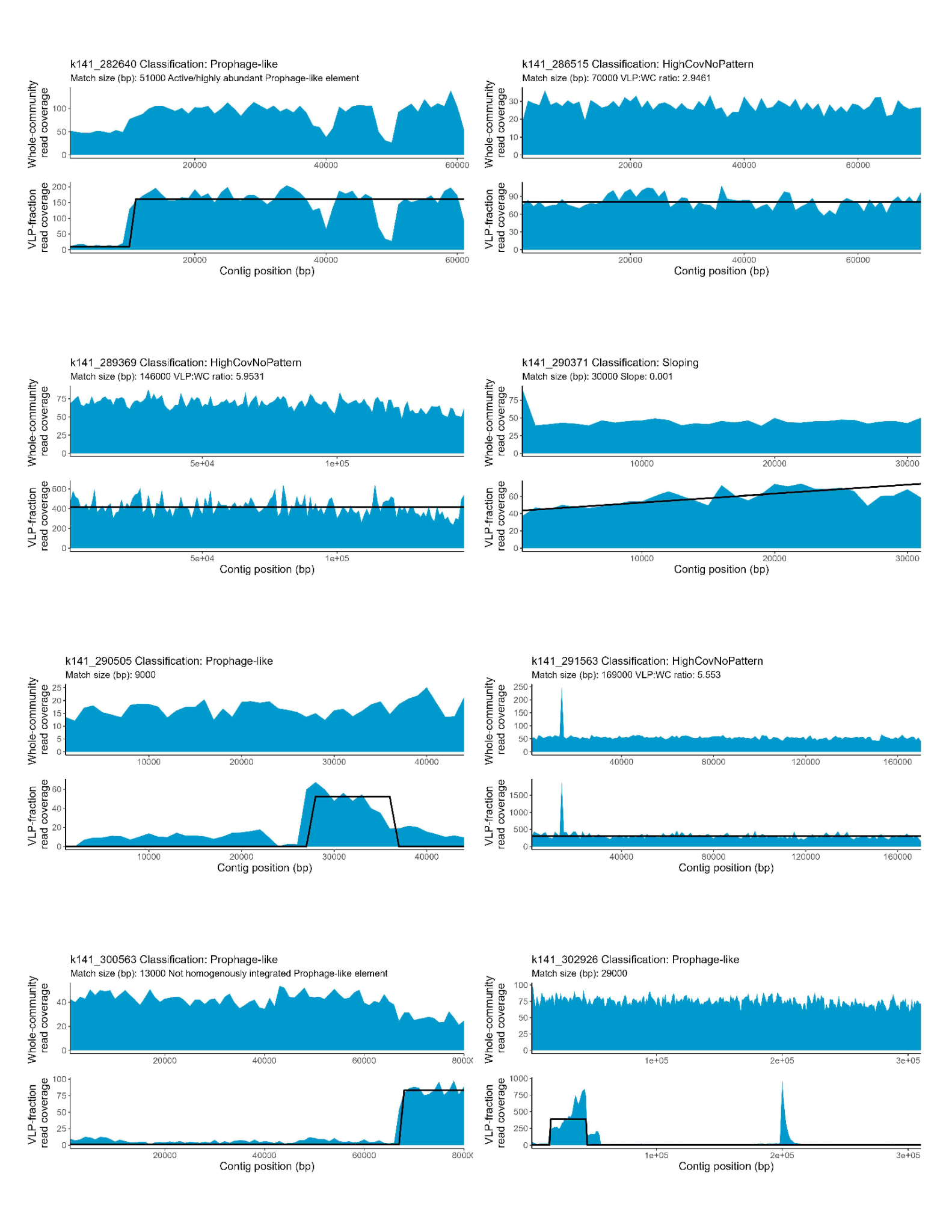

**Supplementary figure 8- TrIdent pattern-match results for replicate 1 in case study data.** Read coverage plots generated with the plotTrIdentResults() function in the TrIdent R package.
